# Developmental timing of gene expression predicts dosage dominance, which tracks neuronal–glial fate in the developing human brain

**DOI:** 10.64898/2026.11.22.768478

**Authors:** In Gyeong Koh, Dong-Min Shin, Beomsu Kim, Seojin Chung, Soo-Whee Kim, Hyeji Lee, Ji Hyun Son, Jeewon Lee, Guiyoung Bong, Jinju Han, Changuk Chung, Hee Jeong Yoo, Joon-Yong An

**Author notes:** Corresponding authors. (H.J.Y.), (J.-Y.A.). These authors contributed equally to this work (In Gyeong Koh, Dong-Min Shin).

## Abstract

Genes implicated in brain disorders have diverse functions, but how altered dosage shapes their effects on developmental cell fate remains unclear. In a cell-state dynamics model of the developing human brain, genes expressed earlier in development responded more strongly to knockout than to overexpression, and this dosage dominance tracked neuronal–glial fate. Fate responses of 14,044 genes formed 31 modules across 21 lineages. Stronger perturbations could reverse fate responses despite correlated predicted expression changes across neuronal destinations. Disease-risk genes occupied shared and distinct developmental programs. Within autism, module timing distinguished genes associated with and without intellectual disability or developmental delay, while de novo and common variation implicated different modules. Knockout signatures of selected autism-associated modules matched the neuronal-down, glial-up pattern that postmortem autism cortex shares with other psychiatric disorders. These findings connect heterogeneous genetic risk through developmental programs whose fate responses depend on dosage and cellular context.

## Main Text

Autism spectrum disorder (ASD) and other neurodevelopmental disorders (NDDs) have a heterogeneous genetic basis, with rare coding variants implicating synaptic, chromatin, transcriptional and signaling processes^1,2^. Recurrent copy-number variants (CNVs) and common variants across the genome also contribute to risk, with genetic overlap extending across clinical presentations and diagnostic boundaries^1,3–10^. Risk genes differ in their expression across progenitor and neuronal populations during brain development^1,2^. Relating genetic risk to the formation and maturation of brain cell types could help explain both shared disease associations and differences in the developmental effects of individual genes.

Neurodevelopmental risk genes show distinct expression patterns across cell populations and stages of brain development^1,2,11^. Genes more strongly associated with developmental delay are enriched in progenitor and immature neuronal transcriptomes, whereas genes more strongly associated with ASD are enriched in maturing neurons^1^. Experimental perturbations of different NDD genes can converge on transcriptional networks whose composition depends on cell type and developmental stage^12–14^. Convergent changes are also evident in neuronal and glial transcriptional programs in postmortem ASD cortex^15–18^. Although these studies identify shared molecular responses, whether such responses lead to similar fate outcomes across dosage levels and developmental states remains unresolved.

Reduced and increased gene dosage can have distinct effects on neural development^19,20^, and neural cell types differ in their transcriptional responses to perturbations of neurodevelopmental risk genes^13,21^. How dosage dominance, defined here as the relative magnitude of responses to reduced and increased expression, relates to developmental timing, gene function, and neuronal– glial fate remains unclear. Comparisons across perturbation strengths would further reveal whether larger molecular changes are accompanied by proportionately greater fate shifts across developmental lineages.

Computational models trained on unperturbed single-cell data can simulate gene knockout or overexpression and predict the resulting shifts in cell identity^22,23^. Such simulations have been applied to risk transcription factors of brain disorders in the human prefrontal cortex^24^, and to nearly all protein-coding genes in brain organoids, where gene clusters defined by predicted expression responses were enriched for ASD risk genes^25^. In these applications, responses were read out as expression changes, and how modules of genes with shared responses alter the proportions of cells reaching each developmental fate in the same system has not been measured. Here, we used scDiffEq, a published framework for modeling cell-state dynamics, to predict how gene perturbations alter the proportions of cells reaching each fate in a human developmental brain atlas^11,26^. We simulated knockout (KO) and overexpression (OE) of 14,044 genes and quantified fate changes across 21 lineages defined by pairs of starting and endpoint cell populations. Genes with similar effects on differentiation, maturation, and maintenance of cell identity across these lineages were grouped into modules. We related gene and module responses to developmental expression and genetic risk, and varied perturbation strength to compare predicted expression changes with fate outcomes. We compared selected model predictions with findings from experimental studies of altered gene dosage and transcriptomic analyses of postmortem ASD cortex.

### Results

### Expanded perturbation landscape defines 31 perturbation-response modules

We constructed a genome-wide perturbation workflow within a human developmental brain cell-state space (**Fig. 1a**), expanding the atlas from eight broad cell classes to 21 annotated cell types (**Extended Data Fig. 1a** and **Supplementary Table 1a**)^11^. Radial glia and neuronal precursor populations differed in composition across sampled brain regions (**Extended Data Fig. 1b**). Marker expression patterns further supported the annotated progenitor, regional, neuronal, and glial identities, including distinct oligodendroglial differentiation states (**Extended Data Fig. 1c**).

**Fig. 1.**
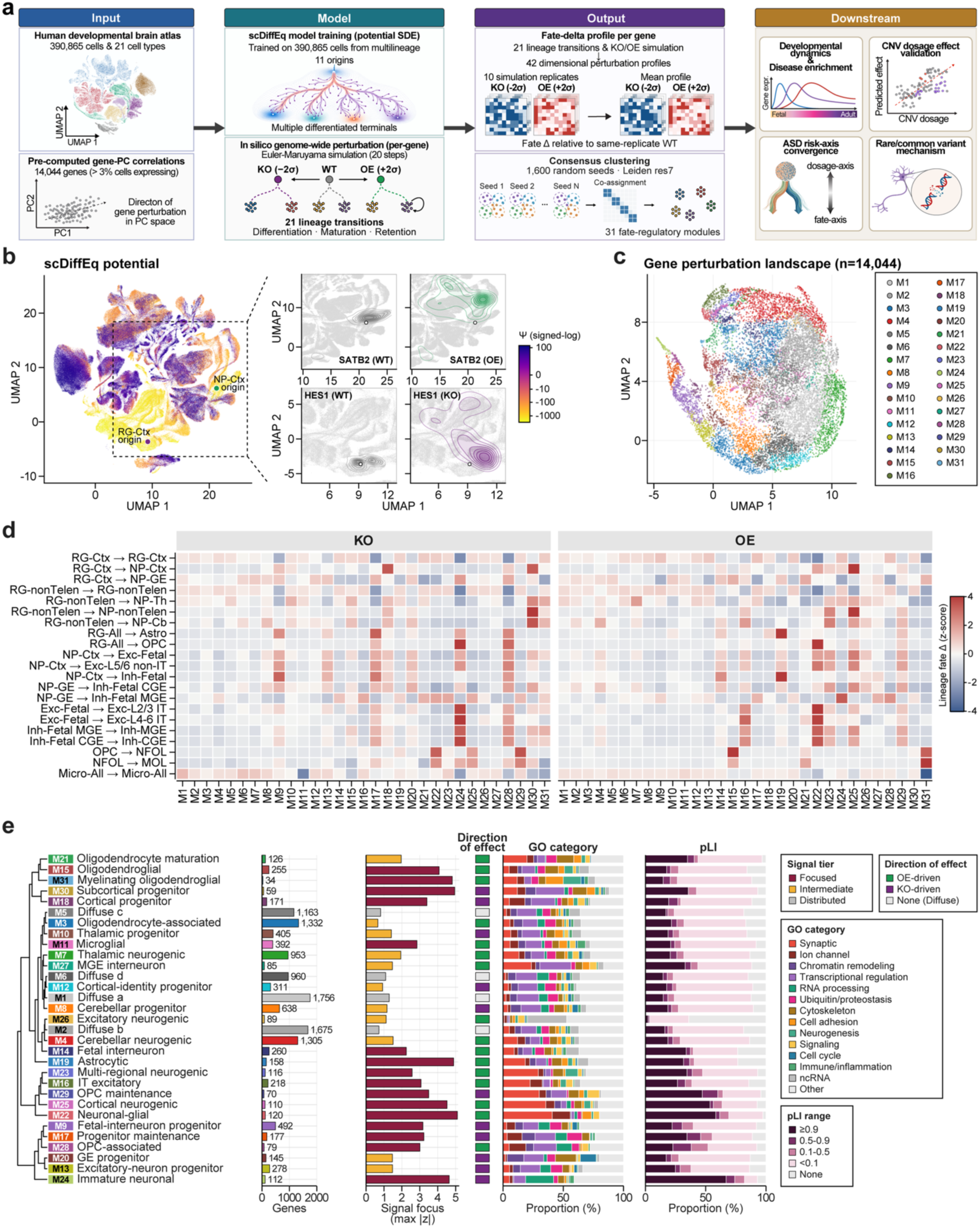
Genome-wide perturbation profiles resolve lineage-associated modules. **a**, Overview of the perturbation workflow. **b**, Learned potential and representative *SATB2* OE and *HES1* KO simulations. Contours show endpoint densities for 500 starting cells per origin. Potential ticks give original units on a signed-log scale. **c**, Gene perturbation profiles grouped into 31 modules. **d**, Module-mean effects standardized across modules for each lineage and condition. **e**, Module annotations, ordered by perturbation-profile similarity. OE-driven and KO-driven indicate the condition that increases occupancy on the representative transition. RG, radial glia; NP, neuronal precursor; Ctx, cortex; GE, ganglionic eminence; Th, thalamus; Cb, cerebellum; nonTelen, non-telencephalic; Exc/Inh, excitatory/inhibitory neuron; MGE/CGE, medial/caudal ganglionic eminence; IT, intratelencephalic; OPC, oligodendrocyte precursor cell; NFOL/MOL, newly formed/mature oligodendrocyte; Astro, astrocyte; Micro, microglia.

Along diffusion pseudotime, cell composition shifted from predominantly radial-glial and neuronal-precursor populations to predominantly neuronal populations, and neuronal and gliogenic markers showed distinct expression profiles across this ordering (**Extended Data Fig. 1d,e**). The refined states provided distinct origins and destinations for evaluating responses across neuronal and glial lineages.

We defined 21 lineages as pairs of starting and endpoint populations, spanning differentiation, maturation, and retention of the starting-cell identity across 11 origin populations (**Extended Data Fig. 2a** and **Supplementary Table 1b**). Using the scDiffEq model trained on this annotated cell-state space, we simulated knockout (KO) and overexpression (OE) of 14,044 genes^26^. For each gene, changes in target-state arrival relative to wild type (WT) across the 21 lineages formed a 42-dimensional KO–OE profile (**Supplementary Table 1c**). These profiles captured both the affected lineages and the direction of each effect. KO–OE effects were negatively correlated across all 17 lineages away from the WT floor or ceiling boundaries (**Extended Data Fig. 2b**), with a pooled correlation of −0.373 (**Extended Data Fig. 2c**). KO and OE generally produced opposing effects, although their responses were not uniformly mirrored. As a lineage-specificity check, microglial endpoint changes from five radial-glial or neuronal-precursor origins exceeded the WT noise threshold for no KO perturbations and only 0.10–0.44% of OE perturbations (**Supplementary Note 1**).

Representative simulations illustrated effects on state retention and neuronal-state arrival within the learned potential landscape (**Fig. 1b**). Simulated *HES1* KO reduced radial-glial retention in cortical (Δ = −0.52) and non-telencephalic populations (Δ = −0.55). *SATB2* OE increased the arrival of cortical neuronal precursors in the fetal excitatory-neuron state (Δ = +0.37) (**Fig. 1b** and **Supplementary Table 1c**). These effects had the same sign in all 10 replicate simulations. Radial-glial retention measures endpoint state occupancy rather than self-renewal or proliferation. To identify shared patterns of perturbation response across lineages, we grouped the joint KO– OE profiles into 31 perturbation-response modules (**Fig. 1c**). The 27 lineage-associated modules comprised 16 focused and 11 intermediate modules. Four additional modules had distributed profiles without a distinctive lineage-associated signature (**Fig. 1d,e** and **Supplementary Table 1d**). Module assignments were highly consistent across clustering-seed batches on the fixed perturbation map (mean adjusted Rand index, ARI = 0.982), whereas agreement between independently averaged simulation profiles was lower (mean ARI = 0.624), with recovery varying across individual modules (**Supplementary Note 2**, **Supplementary Fig. 1** and **Supplementary Table 1e**).

Integrating lineage signatures with cell-type expression and functional annotations yielded a taxonomy of neuronal, glial, and progenitor-associated modules (**Fig. 1e**, **Extended Data Fig. 2d** and **Supplementary Table 1d**). The 27 lineage-associated modules comprised 16 OE-driven and 11 KO-driven groups, based on their representative lineage effects. M15 Oligodendroglial was characterized by a distinctive OE-associated increase in the influx of oligodendrocyte precursor cells (OPCs) into newly formed oligodendrocyte states, whereas M3 Oligodendrocyte-associated showed less pronounced lineage effects and was named primarily for elevated relative expression in mature oligodendrocytes. M22 Neuronal-glial showed high relative expression in excitatory neurons, and synaptic genes constituted the largest functional class within the module. OE-associated increases extended across neuronal and glial transitions, from radial glia to OPCs and from fetal excitatory neurons to layer 2/3 IT neurons.

### Dosage dominance links developmental gene expression to neuronal–glial fate

We mapped perturbation-defined modules onto age- and lineage-resolved expression trajectories to link KO-to-OE dosage dominance to developmental programs in the human brain. Dynamic trajectories were classified as rising, transiently rising, transiently falling, or falling (**Fig. 2a** and **Supplementary Fig. 2a–c**). Falling trajectories predominated in M24 Immature neuronal (100%) and M28 OPC-associated (86.6%), whereas rising trajectories predominated in M14 Fetal interneuron and M21 Oligodendrocyte maturation. Across 27 lineage-associated modules, the late-rising fraction (up and trans-up trajectories) correlated inversely with dosage dominance (Pearson r = −0.90, *P* = 1.4 x 10^−10^) (**Fig. 2b, Extended Data Fig. 3a** and **Supplementary Table 2a).** Earlier developmental expression was associated with larger KO responses relative to OE, whereas later-rising expression was associated with a shift toward OE dominance.

**Fig. 2.**
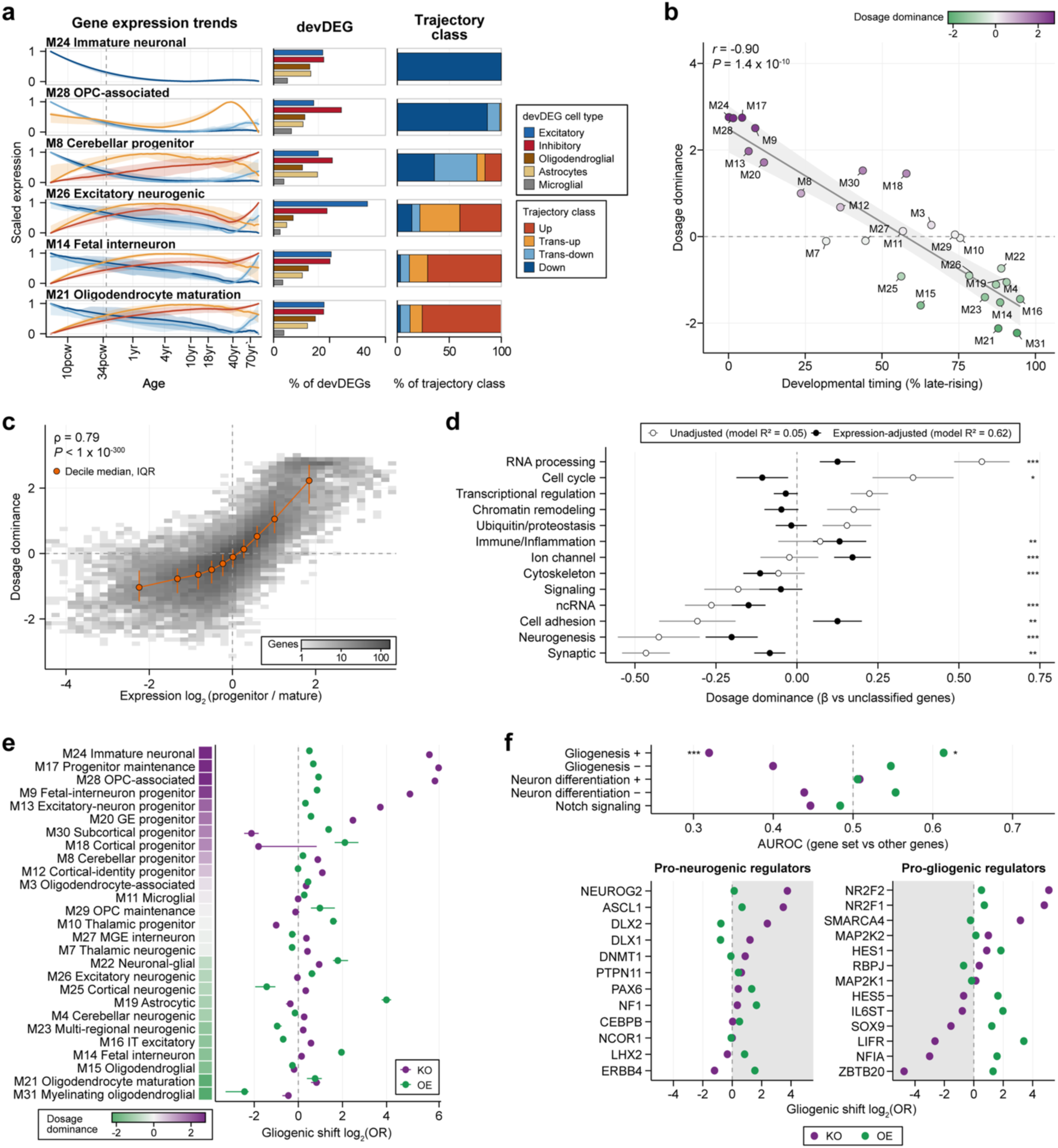
Developmental expression and neuronal–glial fate changes in relation to dosage dominance. Positive dosage dominance indicates larger KO than OE responses. **a**, Developmental expression trajectories of genes in six modules. Curves and shading, medians and interquartile ranges (IQRs); dashed line, birth. **b**, Relationship between developmental timing and dosage dominance across 27 modules. Line and shading, linear fit and 95% confidence band. **c**, Relationship between the progenitor-to-mature expression ratio and dosage dominance across 14,044 genes. Orange points and bars, decile medians and IQRs. **d**, Functional-class coefficients relative to unclassified genes, before and after expression adjustment. Bars, 95% confidence intervals. Asterisks in d and f indicate adjusted *P* < 0.05 (*), <0.01 (**) or <0.001 (***). **e**, Module gliogenic shifts, defined as WT-relative changes in log2 astrocyte/OPC-to-neuronal-precursor odds. Points and bars, gene medians and bootstrap 95% CIs. **f**, Top, Differences in gliogenic shifts between each GO set and other genes. AUROC >0.5 indicates higher gliogenic shifts. Bottom, Gliogenic shifts under KO and OE for 25 regulators. Gray shading indicates the loss-of-function direction reported for the regulator or a combined perturbation.

The timing–dominance relationship was reflected in expression across progenitor and mature cell populations. Genes expressed more highly in progenitor and immature cells than in mature cells showed higher dosage dominance (Spearman ρ = 0.79 across 14,044 genes) (**Fig. 2c** and **Supplementary Table 2b**). The corresponding module-level expression ratio showed the same relationship (Pearson r = 0.93) (**Extended Data Fig. 3b**). RNA-processing and cell-cycle genes had the highest dosage dominance relative to unclassified genes (unadjusted mean differences, β = 0.57 and 0.36), whereas synaptic and neurogenesis genes had the lowest (β = −0.47 and −0.43). To distinguish functional-class differences from their associated expression patterns, we adjusted for mean expression and the progenitor-to-mature expression ratio. Adding these covariates increased explained variance from 4.6% to 61.6% in the 13-class model. The synaptic coefficient was attenuated to −0.08 (95% CI, −0.13 to −0.04), whereas ion-channel genes had a positive adjusted coefficient (0.17; 95% CI, 0.12 to 0.23) (**Fig. 2d**, **Extended Data Fig. 3c** and **Supplementary Table 2c**). Thus, much of the synaptic contrast was accounted for by expression, whereas ion-channel genes showed higher dosage dominance than unclassified genes after adjustment.

To connect dosage dominance with developmental fate outcomes, we compared the neuronal and glial states reached by simulated radial-glial start cells under KO and OE separately. The gliogenic shift was measured as the change from WT in the log2 odds of reaching astrocyte or OPC states relative to neuronal-precursor states. Across 14,044 genes, the magnitude of this shift increased with dosage dominance under KO (Spearman ρ = 0.54) and decreased under OE (ρ = −0.26) (**Fig. 2e** and **Supplementary Table 2d**). The KO-dominant M17 and M24 modules showed large gliogenic shifts under KO and smaller shifts in the same direction under OE. By contrast, M30 Subcortical progenitor and M10 Thalamic progenitor shifted toward neuronal precursors under KO and toward gliogenic states under OE. Overall, KO and OE shifts had opposite directions in 9 of 27 modules (**Supplementary Table 2e**).

At the gene level, dosage dominance was more strongly associated with changes in gliogenic share under KO than under OE (Spearman ρ = 0.46 and 0.08) (**Extended Data Fig. 3d**). The KO relationship was also observed within modules (module-centered rank correlation r = 0.39; 8,490 genes) (**Supplementary Table 2d**). In KO-dominant genes, the increase in gliogenic share primarily reflected more simulated cells reaching glial states. In the highest dosage-dominance decile, the median number of simulated cells reaching astrocyte states increased from 46 under WT to 1,038 under KO. The corresponding median neuronal-precursor count changed from 281 to 247. Radial-glial retention fell from 4,489 to 2,335 of 5,000 start cells, accompanying the higher glial endpoint counts (**Supplementary Table 2b**).

We compared the modeled fate shifts with the known roles of genes in promoting neuronal or glial differentiation (**Fig. 2f** and **Extended Data Fig. 3e**). Among five GO sets, genes annotated to positive regulation of gliogenesis showed lower gliogenic shifts than other genes under KO and higher shifts under OE (KO AUROC = 0.32, adjusted *P* = 6.9 x 10^-5^) (OE AUROC = 0.61, adjusted *P* = 2.3 x 10^-2^) (**Supplementary Table 2f**). The other four sets did not reach the corrected significance threshold. Among 25 selected regulators, modeled KO produced 12 matching directions, seven small shifts and six opposite directions (**Supplementary Table 2g**). *NFIA* KO shifted the neuronal–glial balance toward neuronal precursors, whereas *NFIA* OE shifted the balance toward glial states. The OE direction was consistent with glial competence induced by transient *NFIA* expression in human neural stem cells^27^. Conversely, *DLX2* KO shifted the balance toward glial states, consistent with premature oligodendroglial differentiation after combined *Dlx1*/*Dlx2* loss in the developing mouse forebrain^28^. Because modeled fate shifts arise from perturbations along gene-associated directions in expression space, correspondence with known regulators could reflect where each gene is expressed rather than the modeled dynamics. We therefore compared the modeled effects with scores derived directly from expression differences between glial and neuronal-precursor states. These scores reproduced the direction of the association for the gliogenesis gene set (KO AUROC = 0.36, OE AUROC = 0.64, both adjusted *P* = 0.0028) (**Supplementary Table 2f**). Without a magnitude threshold, modeled KO shifts had the reported sign for 16 of the 25 regulators, compared with 15 for the expression-based scores. For ten regulators with reported gain-of-function effects, the corresponding counts were five for modeled OE shifts and seven for the expression-based scores (**Supplementary Table 2g**).

A single-cell CRISPRi screen of 44 transcription factors in primary radial glia from human cortex at 16–18 gestational weeks provided an independent comparison^29^. Among the 43 transcription factors included in the perturbation map, modeled and experimental effects had matching signs for 23 and opposite signs for 20. *ARX* knockdown and modeled KO both reduced inhibitory relative to excitatory output. Across these factors, the correlation between modeled KO shifts in inhibitory-to-excitatory output and measured knockdown effects after 7 days of differentiation was weak (Spearman ρ = 0.15, one-sided permutation *P* = 0.17). For a score based on the expression difference between fetal excitatory and inhibitory neurons, the corresponding correlation was 0.30 (*P* = 0.026). The model–experiment partial correlation after adjustment for this expression-based score was −0.04 (*P* = 0.59) (**Extended Data Fig. 3f** and **Supplementary Table 2h**). Cross-validated regressions using expression, modeled KO responses or both showed no clear improvement over a mean-effect control (**Extended Data Fig. 3g** and **Supplementary Table 2h**).

### Perturbation strength separates fate responses within shared developmental programs

We varied perturbation strength to determine whether increasing dosage disruption amplified a fate response or redirected the response toward a different destination. Across KO and OE strengths of 0.5–4 gene-expression standard deviations (s.d.), lineage-specific responses persisted, weakened or reversed sign, corresponding to stable, unstable and sign-shift classes (**Fig. 3a** and **Supplementary Table 3a**). The proportions of these classes differed across lineages and the 27 lineage-associated modules (**Fig. 3b**). Across all 31 modules, cortical neuronal precursors had the largest number of genes whose response class differed between destinations (n = 1,908), followed by diencephalic, midbrain and hindbrain radial glia (n = 1,333) and cortical radial glia (n = 838) (**Supplementary Table 3b**). Cortical precursors did not have the highest discordant fraction (22%), but discordance extended across 19 modules containing at least 20 such genes (**Extended Data Fig. 4a**).

**Fig. 3.**
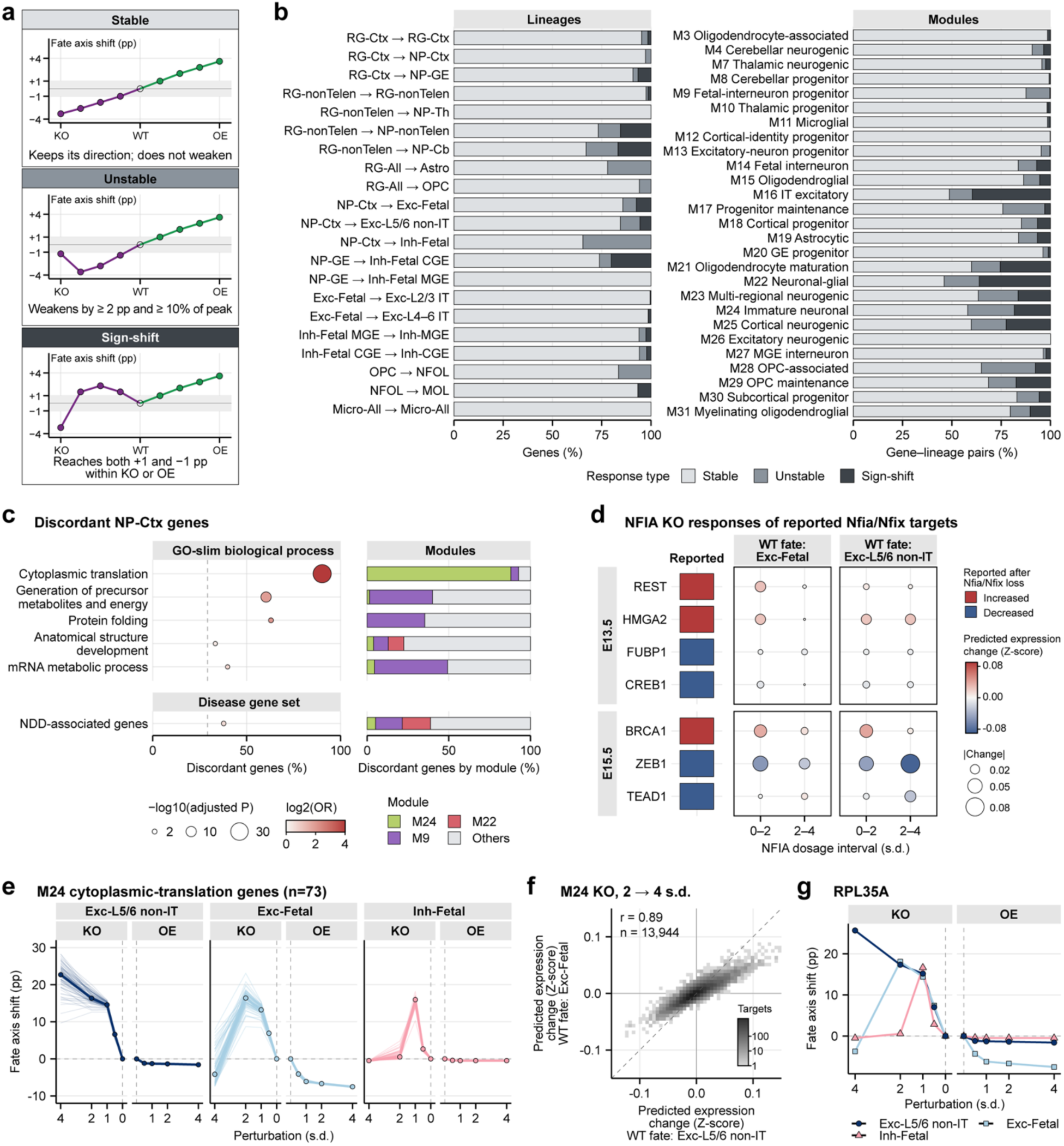
Perturbation strength separates destination-dependent fate responses. **a**, Schematic stable, unstable, and sign-shift responses across KO and OE perturbation strengths. Fate axis shift denotes the WT-relative change in the proportion of simulated cells reaching a specified destination state, expressed in percentage points (pp). **b**, Response-class proportions across lineages and 27 lineage-associated modules. **c**, Enrichment and module composition of destination-discordant cortical precursor genes. Dashed line, background discordant fraction (1,903/6,519 genes). **d**, Modeled expression changes under *NFIA* KO compared with reported changes after combined *Nfia*/*Nfix* loss. WT-fate groups comprise starting cells grouped by their matched WT destination. Columns compare WT with 2 s.d. (0–2) and 2 with 4 s.d. (2–4). Ages refer to the source mouse experiments. **e**, Fate responses to perturbation of M24 cytoplasmic-translation genes. Thin lines, individual gene means across ten replicates; thick lines, gene medians. **f**, Mean additional expression changes across the 73 KO perturbations from 2 to 4 s.d., compared between WT-fate groups. Dashed line, equal responses. Expression changes in d and f exclude the imposed input change and use model-standardized coordinates. **g**, Fate responses across KO and OE strengths for *RPL35A*.

Across cortical precursor destinations, discordant genes were most strongly enriched for cytoplasmic translation. Within the 27 lineage-associated modules, 1,903 of 6,519 genes evaluable in at least two cortical precursor destinations had discordant response classes (**Supplementary Table 3c**). Cytoplasmic-translation genes showed the strongest enrichment, with discordance in 83 of 92 genes (odds ratio [OR] = 23.34, adjusted *P* = 7.5 x 10^−33^) (**Fig. 3c**). Nine of 145 gene sets were enriched after correction, including four other biological-process categories, two neuronal developmental signatures, and two disease sets. The disease associations extended to NDD genes (116 of 307, OR = 1.50, adjusted *P* = 0.016) and ASD or developmental-disorder genes (91 of 238, OR = 1.53, adjusted *P* = 0.029) (**Supplementary Table 3d**)^30^. Most discordant cytoplasmic-translation genes belonged to M24 Immature neuronal (73 of 83), whereas M9 Fetal-interneuron progenitor contributed 35–45% of the discordant genes in the energy-metabolism, protein-folding and mRNA-metabolism sets (**Supplementary Table 3e**).

The NDD-associated regulator *NFIA* provided a link between destination discordance and a known developmental program. Combined *Nfia*/*Nfix* loss delays temporal progression in mouse cortical radial-glial progenitors and alters transcription factors associated with developmental transitions^31^. We compared these reported changes with modeled *NFIA* KO in starting cells grouped by their matched WT destination (WT-fate groups). At 2 s.d., all seven transcription factors represented in the model changed in the reported direction in both the fetal excitatory and layer 5/6 non-IT WT-fate groups (**Fig. 3d** and **Supplementary Table 3f**). *REST*, *HMGA2* and *BRCA1* increased, whereas *FUBP1*, *CREB1*, *ZEB1* and *TEAD1* decreased. Stronger KO differentially amplified these responses. From 2 to 4 s.d., the *HMGA2* response increased from 0.024 to 0.048 in the model’s standardized expression coordinates in the layer 5/6 non-IT group but remained near 0.024 in the fetal excitatory group. *ZEB1* decreased further in both groups, more strongly in the layer 5/6 non-IT group. Thus, shared directional responses to *NFIA* loss acquired different magnitudes across the two WT-fate groups as perturbation strength increased. The 73 discordant M24 cytoplasmic-translation genes showed a more pronounced separation of fate responses with increasing KO strength (**Fig. 3e** and **Supplementary Table 3g**). At 2 s.d., the median increases over WT in the percentage of simulated cells reaching layer 5/6 non-IT and fetal excitatory states were similar (+16.34 and +16.40 percentage points [pp]). At 4 s.d., the layer 5/6 non-IT shift increased to +22.70 pp, whereas the fetal excitatory shift reversed to −4.16 pp. The median fetal inhibitory response peaked at 1 s.d. (+15.96 pp) and declined thereafter. Median OE shifts remained negative, with decreases in layer 5/6 non-IT and fetal inhibitory occupancy constrained by low WT levels of 1.58% and 0.48% (**Extended Data Fig. 4b**). The same group of perturbed genes therefore promoted two excitatory destinations at intermediate KO strength but produced opposing responses at the stronger perturbation.

The divergence in fate responses was accompanied by similar predicted expression changes. The additional KO responses from 2 to 4 s.d. were strongly correlated between the two excitatory WT-fate groups across 13,944 target genes (Pearson r = 0.89) (**Fig. 3f** and **Supplementary Table 3h**). Correlations also remained high for individual perturbations (r = 0.87–0.91), although greater fate divergence was associated with lower correlation within this range (Spearman ρ = −0.80, *P* = 1.8 x 10^-17^, n = 73 perturbations) (**Extended Data Fig. 4c,d** and **Supplementary Table 3i**). *RPL35A* illustrated this combination. From 2 to 4 s.d. KO, the shift toward layer 5/6 non-IT states increased from +17.34 to +25.68 pp, whereas the shift toward fetal excitatory states fell from +18.10 to −3.74 pp (**Fig. 3g**). Predicted expression changes in the two groups nonetheless remained correlated (r = 0.88). Together, the *NFIA* and M24 examples show that shared patterns of predicted expression responses can accompany destination-dependent fate outcomes as perturbation strength increases.

### Developmental response programs connect brain-disorder risk to ASD cortical changes

Genetic risk is partly shared across brain disorders^3,4^, but genetic overlap alone does not establish whether risk genes affect the same developmental programs. We compared the module distributions and lineage responses of risk genes across neurodevelopmental, brain-malformation, white-matter, neurodegenerative and psychiatric disorders. Disease-associated genes were enriched in 13 modules (adjusted *P* < 0.05) (**Fig. 4a** and **Supplementary Table 4a**,**b**). Alzheimer’s disease (AD) genes were enriched in microglial M11^32^ (OR = 5.30, adjusted *P* = 7.46 x 10^−7^). The enrichment persisted against covariate-matched backgrounds (4.3-fold, adjusted *P* = 0.0085) (**Extended Data Fig. 5a**), consistent with genetic evidence implicating microglia^33,34^. Leukodystrophy genes were enriched in oligodendroglial M15 and cerebellar progenitor M8. The microglial–oligodendroglial contrast extended across the module distributions of nonoverlapping AD and leukodystrophy genes (total variation distance = 0.66, Holm-adjusted *P* = 0.00040) (**Extended Data Fig. 5b** and **Supplementary Table 4c**). Microcephaly genes implicated ganglionic eminence progenitor M20, whereas lissencephaly genes implicated immature neuronal M24. Across several diagnoses, neuronal–glial M22 and cortical neurogenic M25 were enriched for Parkinson’s disease, major depressive disorder, epilepsy, ASD and developmental disorder (DD) genes. ASD without intellectual disability or developmental delay (ID/DD) additionally implicated fetal interneuron M14 (adjusted *P* = 0.0035).

**Fig. 4.**
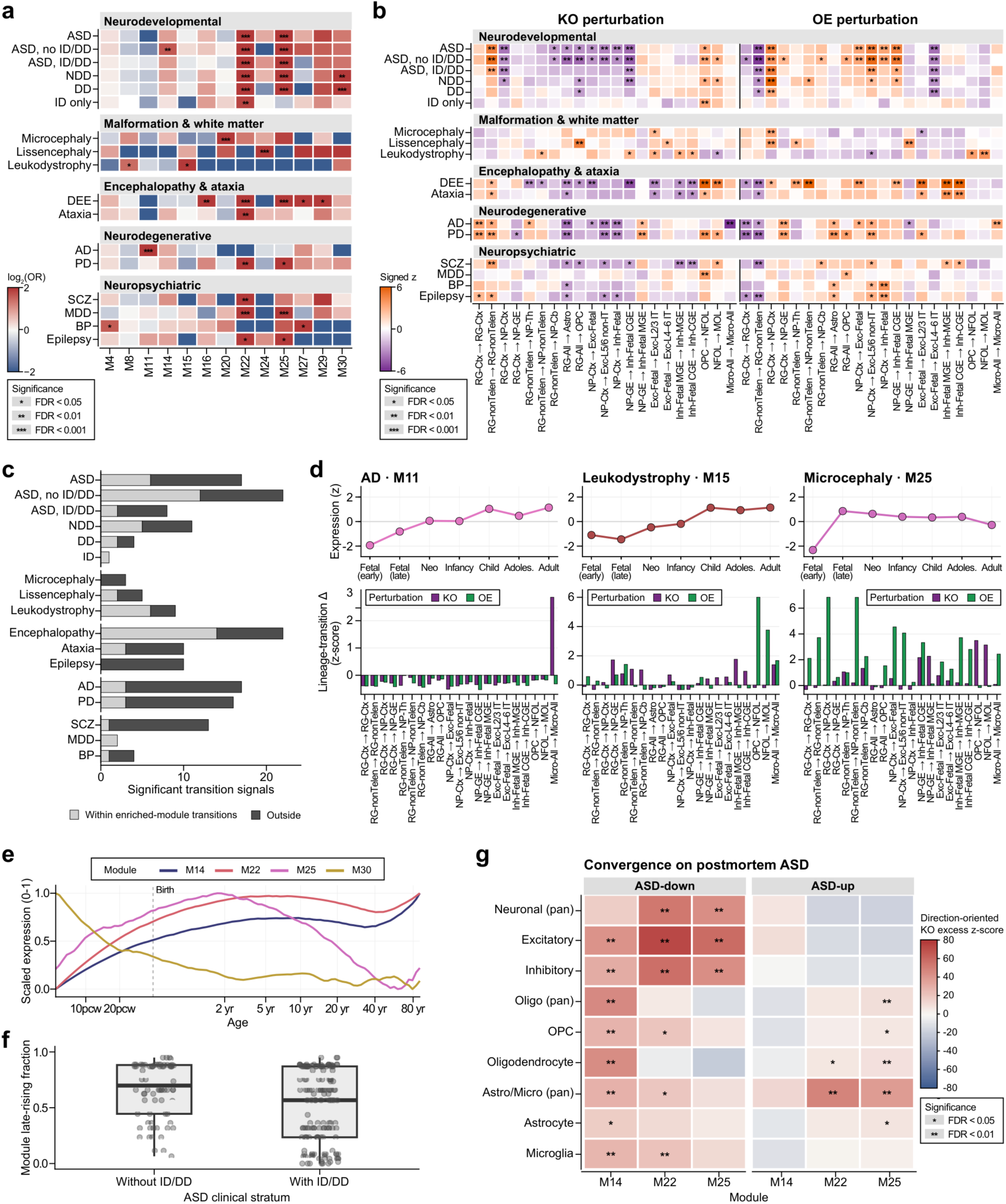
Disease-associated developmental programs and convergence with ASD cortical expression. **a**, Disease-gene enrichment across modules. **b**, Signed KO/OE transition responses relative to matched expectations. **c**, Significant responses within or outside the three principal transitions of disease-enriched modules. **d**, Developmental expression and perturbation responses of representative disease–module pairs. Upper, effect-weighted expression across developmental stages. Lower, module response magnitudes standardized against genome-wide responses. **e**, Module expression trajectories scaled to 0–1; dashed line, birth. **f**, Late-rising fractions of modules assigned to genes associated exclusively with ASD without (n = 74) or with ID/DD (n = 159). Points, genes; boxes, median and IQR; whiskers, 1.5 x IQR. **g**, KO-signature concordance with postmortem ASD expression sets for M14, M22 and M25 (10, 18 and 13 ASD genes). Colors show Stouffer z relative to the matched mean, sign-reversed for ASD-down sets so positive values indicate concordance.

We examined which developmental transitions responded to disease-gene perturbations and how closely those responses followed the enriched modules. Across 21 transitions and 17 disease sets, model KO and OE produced significant responses in 174 disease–direction–transition combinations (adjusted *P* < 0.05) (**Fig. 4b** and **Supplementary Table 4d**). Of these, 111 (63.8%) lay outside the three principal transitions of modules enriched for the corresponding disease set (**Fig. 4c** and **Supplementary Table 4d**). AD genes had opposing KO and OE responses at the microglial transition (signed z = −7.22 for KO and 3.37 for OE), consistent with their enrichment in M11 (**Fig. 4d**). OE of leukodystrophy genes increased the transition from oligodendrocyte precursors to newly formed oligodendrocytes (z = 3.72, adjusted *P* = 0.013), matching M15’s defining transition. OE of microcephaly genes increased the transition from cortical radial glia to neural progenitors (z = 3.46, adjusted *P* = 0.0086), although enrichment in M25 was not significant (adjusted *P* = 0.22). This response linked microcephaly genes to cortical neurogenesis beyond the ganglionic eminence program highlighted by M20 enrichment^35,36^.

Earlier studies associated stronger ASD than DD risk with later expression and more differentiated neurons^1,2^. We compared ASD and DD genes to determine whether this developmental distinction was reflected in the perturbation-response modules. ASD-only genes occupied later-rising modules than DD-only genes, in the direction reported previously, although the difference was small after expression and constraint adjustment (0.027, 95% CI, −0.059 to 0.112) (**Supplementary Table 4e**). A clearer separation emerged within ASD along the ID/DD axis (**Fig. 4e,f**)^30^. Genes associated exclusively with ASD without ID/DD occupied later-rising modules than those associated exclusively with ASD with ID/DD (adjusted difference = 0.136, 95% CI, 0.065– 0.207) (**Extended Data Fig. 5c,d**). Neither gene-level timing contrast was significant (regression *P* = 0.34–0.50) (**Supplementary Table 4e**). The timing distinction thus lay in module membership along the ID/DD axis within ASD.

This clinical timing distinction was also reflected in transition responses. Under KO, genes unique to the ID/DD-associated ASD stratum showed greater early-relative-to-late response magnitudes than genes unique to the no-ID/DD stratum (expression- and constraint-matched Holm-adjusted *P* = 0.0032) (**Extended Data Fig. 5e** and **Supplementary Table 4f**). Transcriptional-regulation genes showed a similar early bias relative to synaptic genes under OE (Holm-adjusted *P* = 0.038). Neither contrast significantly exceeded within-module expectations (Holm-adjusted *P* = 1.0 for the clinical strata and 0.55 for the functional classes). Across the broader ASD and NDD gene sets, signed OE drive through neurogenesis and neuronal-differentiation transitions exceeded covariate-matched expectations. This shift persisted after matching for initial perturbation size (ASD, matched AUROC = 0.57, Holm-adjusted *P* = 0.0004; NDD, matched AUROC = 0.54, Holm-adjusted *P* = 0.0024) (**Extended Data Fig. 5f** and **Supplementary Table 4g**).

Given the contribution of haploinsufficiency to ASD risk^1,2^, we focused on KO-induced expression signatures when comparing modeled responses with gene-expression changes reported in postmortem studies of the ASD cortex^15–18,37^. Signatures from ASD risk genes in M14, M22 and M25 converged with postmortem cortical changes (**Fig. 4g** and **Supplementary Table 4h**). M22 and M25 aligned with neuronal genes downregulated in ASD (Stouffer z = −82.5 and −68.7, versus matched means of −27.9 and −23.4). They also aligned with upregulated astrocyte– microglial genes (z = 75.8 and 61.4, versus matched means of 26.4 and 22.3; all adjusted *P* < 0.005), capturing the cortical neuronal-down and glial-up pattern^17,37,38^. Because neuronal downregulation and astrocyte upregulation are shared with schizophrenia and bipolar disorder^37^, we examined whether the convergence was specific to ASD. ASD-associated modules ranked highly for ASD changes, including ASD-specific contrasts (exact permutation *P* = 0.009–0.035) (**Extended Data Fig. 5g**). This ranking weakened with developmental-progression adjustment (*P* = 0.048–0.086), and the full ASD risk-gene set showed no significant excess concordance over expression- and constraint-matched genes (all adjusted *P* > 0.05) (**Supplementary Table 4h**). For genes associated with ASD without ID/DD, KO signatures showed excess concordance with ASD-specific residual expression changes in the microarray dataset^37^ (stratified Cliff’s δ = 0.105, adjusted *P* = 0.040) (**Supplementary Table 4h**).

### Single-gene responses within CNV intervals partly match gene-level dosage experiments

Recurrent CNVs alter the dosage of multiple genes, creating a need to distinguish the developmental effects of individual genes within each interval. Across the scored gene set, the 202 CNV-supported genes had a higher median bidirectional perturbation effect than the remaining 11,110 genes (median 0.320 versus 0.258, AUROC = 0.570, *P* = 3.3 x 10^−4^) (**Fig. 5a and Supplementary Table 5a**)^1^. The five highest-ranking deletion-supported genes and five highest-ranking duplication-supported genes showed distinct lineage-response profiles (**Fig. 5b** and **Supplementary Table 5b**).

**Fig. 5.**
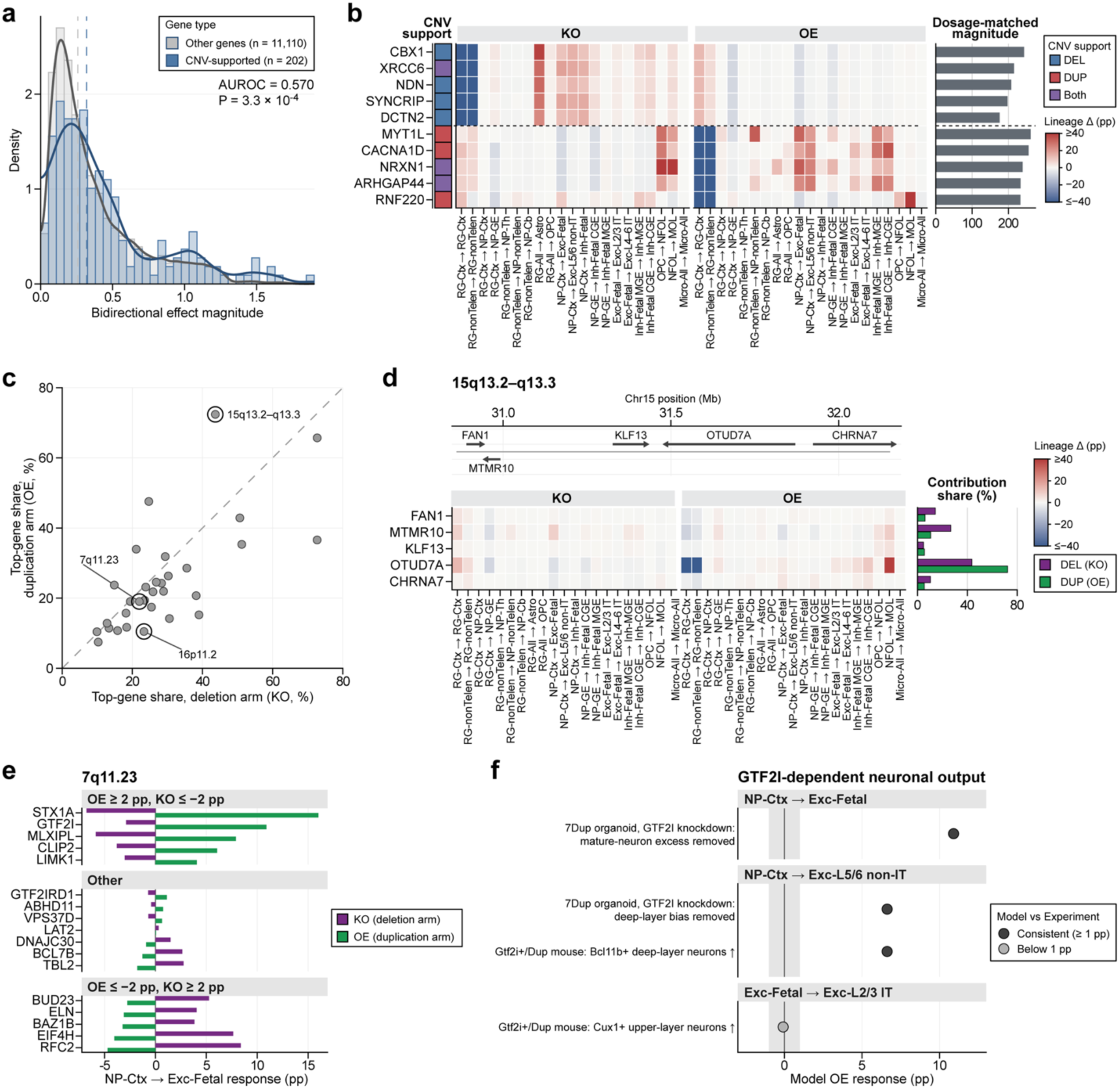
CNV intervals as dosage proxies for the perturbation map. **a**, Bidirectional perturbation effects of CNV-supported and remaining scored genes. Dashed lines, medians. **b**, Lineage responses of the five highest-ranking deletion-supported genes (above divider) and duplication-supported genes (below). **c**, Top-gene shares of within-arm response scores across 31 reciprocal CNV intervals. Rings identify intervals with gene-level experiments; dashed line, equal shares. **d**, Gene positions, lineage responses and contribution shares at 15q13.2–q13.3. **e**, Changes from WT in cortical precursor-to-fetal excitatory output for 7q11.23 genes. **f**, Modeled *GTF2I* responses corresponding to neuronal-output findings. All four findings predict increased output under OE. Gray band, absolute changes below 1 pp.

We ranked dosage-matched responses within 31 reciprocal CNV intervals and compared selected intervals with gene-level experiments (**Fig. 5c and Supplementary Table 5d**). At 15q13.3 (15q13.2–q13.3), *OTUD7A* had the largest share of the within-arm response score under deletion (43.7%) and duplication (72.4%), with the latter the highest top-gene share among the 31 duplication arms^39^ (**Fig. 5d and Supplementary Table 5e**). *OTUD7A* was also the only tested gene whose re-expression restored both dendritic spine density and dendrite outgrowth in Df(h15q13)/+ neurons^40^.

The experimental comparisons also addressed the direction of single-gene dosage effects. At 7q11.23, *GTF2I* knockdown removes the excess mature excitatory neurons in duplication organoids^41^. Modeled *GTF2I* OE increased cortical neuronal-precursor output to fetal excitatory neurons (+10.9 pp), whereas KO decreased output (−2.88 pp) (**Fig. 5e**). The responses agreed with three of four neuronal-output findings, although two deep-layer findings shared one model readout and the upper-layer response was below 1 pp (**Fig. 5f and Supplementary Table 5f**). Progenitor findings were less consistent across the full set of 15 reported findings (**Extended Data Fig. 6a**). At 16p11.2, *KCTD13* ranked 21st of 21 genes in the deletion arm and 19th of 21 in the duplication arm (**Extended Data Fig. 6b and Supplementary Table 5e**). Radial-glial retention nevertheless changed in the direction of three reported proliferation findings, including an increase under KO (+1.98 pp), providing an indirect comparison with proliferation (**Extended Data Fig. 6c** and Supplementary Table 5f)^42^.

### Rare and common risk variants are associated with different perturbation modules

Rare variants of large effect, including de novo and copy-number variants, and polygenic common variants contribute to risk in proportions that differ between disorders^3,4^. Common-variant liability is partly shared across disorders and partly disorder-specific^3,4^. Protein-truncating variants can disrupt one functional copy, whereas common regulatory variants can exert more modest effects on expression^3,43,44^. We compared de novo burden and within-family polygenic transmission in autism sequencing cohorts and gene-set association across nine psychiatric and neurological disorders to determine whether rare and common risk implicated the same developmental response programs (**Fig**. **6a**).

**Fig. 6.**
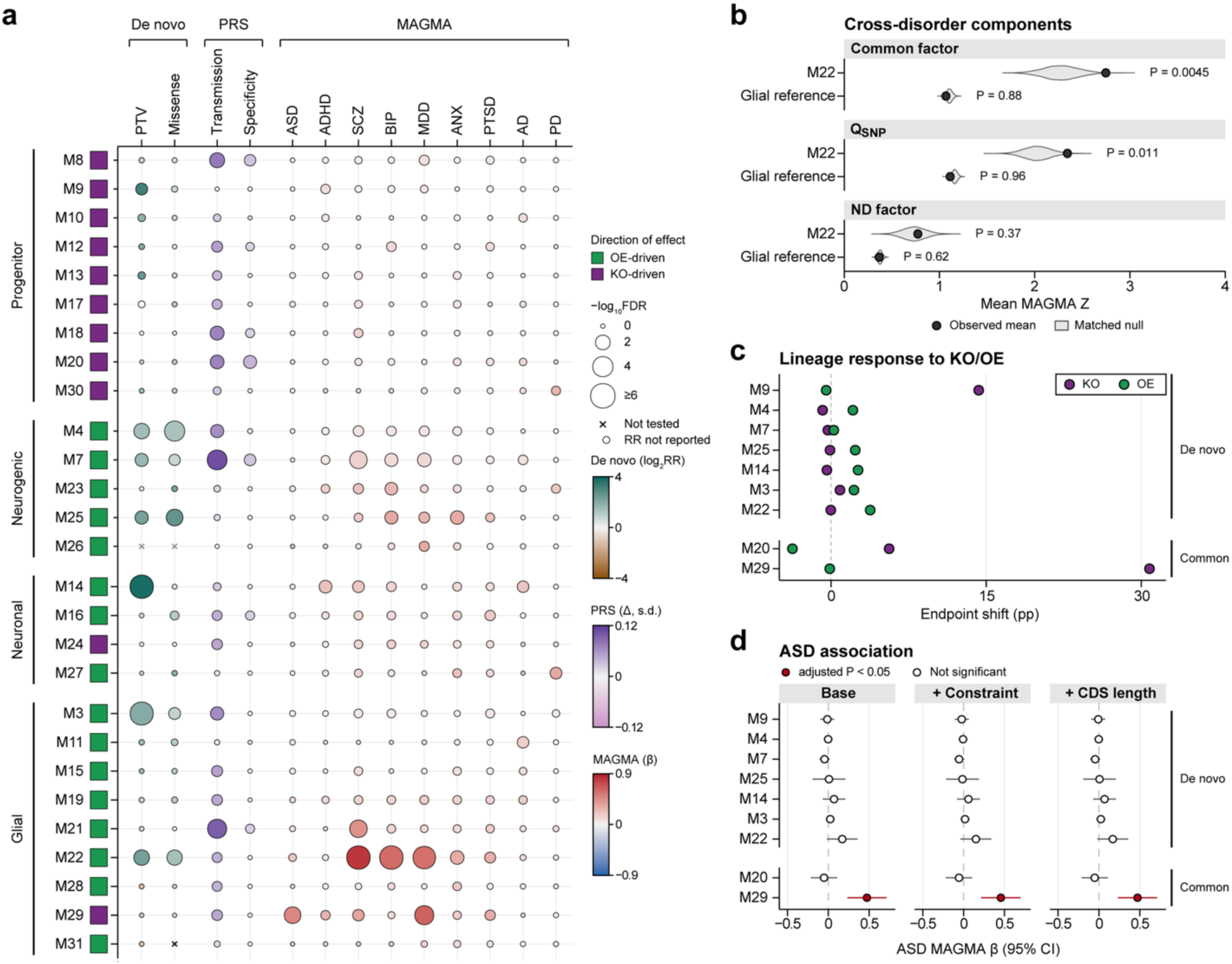
Rare and common risk variants are associated with different perturbation modules. **a**, Module associations with de novo burden (PTVs in genes in the first LOEUF decile; missense variants with MPC ≥2), ASD polygenic score transmission and common-variant risk across nine disorders. PRS Δ is in s.d. of the midparent score. **b**, Mean gene-level MAGMA Z scores for Genomic SEM components in M22 (110 genes) and a glial-reference set (1,874 genes). Violins, matched gene-set distributions; *P*, unadjusted empirical *P*. ND, ASD–ADHD factor; Q_SNP_, heterogeneity unexplained by the common factor. **c**, Mean changes from WT in the percentage of simulated cells reaching representative lineage endpoints (pp). **d**, ASD association coefficients and 95% CIs in the same 16,873 covariate-complete genes, without or with constraint or coding-length adjustment.

Damaging de novo variants were enriched in seven of the 27 modules. Across 28,147 autism probands and 11,616 unaffected siblings, protein-truncating variants (PTVs) in constrained genes (first LOEUF decile) were enriched genome-wide (relative risk [RR] = 3.33). Burden of these PTVs or missense variants (MPC ≥ 2) passed correction in seven modules, referred to below as de novo modules (**Fig. 6a** and **Supplementary Table 6a**). M14 Fetal interneuron had the strongest PTV enrichment (RR = 14.24, 95% CI, 3.49–58.07, adjusted P = 4.3 x 10^−7^) and was the only module whose lower confidence bound exceeded the genome-wide estimate. M22 Neuronal-glial was enriched for both PTVs (RR = 4.95, 95% CI, 1.79–13.73, adjusted P = 0.0031) and missense variants (MPC ≥ 2) (RR = 2.64, 95% CI, 1.47–4.73, adjusted P = 0.0039), as were M3 Oligodendrocyte-associated, M4 Cerebellar neurogenic and M25 Cortical neurogenic, whereas M7, M9 and M14 were enriched only for PTVs.

Genes hit by both damaging variant classes were enriched in M14, M22 and M25 relative to genes hit by only one class (OR = 3.58) (**Extended Data Fig. 7a**). These modules contained 15.2% of damaging de novo events in probands and 9.2% in siblings (OR = 1.76) (**Extended Data Fig. 7b**). Among missense-hit genes, synaptic and ion-channel genes had lower dosage dominance than chromatin-remodeling and transcriptional-regulatory genes (Cliff’s δ = −0.39), with the same contrast across all modeled genes (**Extended Data Fig. 7c,d**). Genes recurrently hit by damaging de novo variants also showed convergence with ASD risk-gene perturbation profiles (**Extended Data Fig. 7e** and **Supplementary Table 6a**).

Within-family transmission implicated a different distribution of common-variant ASD risk. In 6,529 affected and 3,231 unaffected offspring from SPARK, SSC and MSSNG, ASD polygenic scores restricted to module genes were overtransmitted to affected offspring in 12 of the 27 modules, including M22 and M25, and to unaffected offspring in none (**Supplementary Table 6b**). The affected–unaffected contrast passed correction in seven modules (**Fig. 6a** and **Supplementary Table 6b**). M20 GE progenitor showed the most robust excess over the matched set expectation (empirical *P* = 0.00075, adjusted *P* = 0.020), which remained below the correction threshold when its empirical P was increased by two Monte Carlo standard errors. M7 Thalamic neurogenic, which was also enriched for de novo PTVs, passed correction (empirical *P* = 0.0035, adjusted *P* = 0.047) but not after a one-standard-error increase. No significant excess was observed for M22 (empirical *P* = 0.64), M25 (*P* = 0.53) or M14 (*P* = 0.81) (**Fig. 6a** and **Supplementary Table 6b**). At the population level, gene-set associations across nine disorders identified a distinct common-variant ASD module. M29 OPC maintenance was the only module associated with ASD after correction across 27 modules (β = 0.480, adjusted *P* = 0.0013) (**Fig. 6a** and **Supplementary Table 6c**). Several de novo modules were associated with other disorders. M22 was associated with SCZ (adjusted *P* = 2.3 x 10^−11^), BIP, MDD and ANX, M7 with SCZ, BIP and MDD, M25 with BIP and ANX, and M14 with SCZ and ADHD. The associations of M22 with SCZ, BIP and MDD, of M14 with SCZ and ADHD, and of M7 with SCZ remained after joint adjustment for gene constraint against heterozygous loss of function (s_het_) and coding-sequence length (**Supplementary Table 6c**).

We next decomposed the cross-disorder signal of M22 with Genomic SEM across seven disorders, including ASD^45^. The mean MAGMA Z of the 110 eligible M22 Neuronal-glial genes exceeded matched gene-set expectations both for the common factor, which captures liability shared across the disorders, and for Q_SNP_, which captures heterogeneity not explained by the common factor (common factor, *P* = 0.0045; Q_SNP_, *P* = 0.011), whereas a glial-reference set did not (*P* = 0.88 and 0.96) (**Fig.** 6b and **Supplementary Table 6d**). M22 was not enriched for the neurodevelopmental factor defined by ASD and ADHD (*P* = 0.37) (**Fig. 6b**). Within M22, the common-factor gene-level profile was most similar to that of the internalizing factor (Spearman ρ = 0.94), and the Q_SNP_ profile to that of SCZ (ρ = 0.69).

Modules associated with the two classes of risk also responded differently to dosage (**Fig. 6c** and **Supplementary Table 6e**). In M22 and M25, overexpression increased the representative lineage transitions, whereas knockout had little effect. OE increased the RG-to-OPC endpoint by 3.8 pp for M22 and the cortical RG-to-NP endpoint by 2.3 pp for M25, whereas KO changed each by less than 0.2 pp. Six of the seven de novo modules were OE-driven (**Fig. 6a**). The exception, M9 Fetal-interneuron progenitor, was KO-driven. KO of M9 genes increased the cortical NP-to-fetal interneuron transition by 14.3 pp, whereas OE of M14 genes increased the same transition by 2.6 pp. By contrast, M20 and M29 were KO-driven. KO of M29 genes increased the OPC-to-newly formed oligodendrocyte endpoint by 30.8 pp, and M20 shifted the GE NP-to-CGE interneuron transition in both directions (KO, +5.6 pp, OE, −3.7 pp).

The M29 ASD association remained after adjustment for gene constraint and coding length. In the 16,873 genes with complete covariates, the base-model coefficient was 0.473 (adjusted *P* = 0.0015), compared with 0.456 after adjustment for s_het_ (adjusted *P* = 0.0027) and 0.472 after adjustment for coding-sequence length (adjusted *P* = 0.0016) (**Fig. 6d** and **Supplementary Table 6c**). Modules with de novo burden did not pass correction for common-variant ASD association in any model. For the four largest of these modules (M3, M4, M7 and M9, 435–1,104 genes in the covariate-complete set), the confidence intervals excluded effects as large as the M29 estimate.

## Discussion

Our perturbation map connects functionally diverse genes through shared neuronal and glial response programs, linking genetic risk across diagnoses while distinguishing programs associated with different forms of risk within ASD. The coexistence of these shared programs with dosage- and context-dependent fate outcomes suggests that genetic convergence need not entail a uniform course of neural development.

Recent experiments connect convergence among neurodevelopmental risk genes to progenitor maintenance and neural differentiation. ASD-associated transcriptional regulators share cortical binding targets, and knockdown of selected regulators in mouse cortical cultures reduces cells expressing neuronal markers while increasing cells expressing astrocytic markers^46^. In human organoids, knockdown of other ASD genes produces overlapping transcriptional responses and, for selected genes, increases excitatory-neuron proportions while reducing radial-glial proportions^47^. Lineage tracing in primary human and mouse cortical progenitors further shows that the effects of transcription-factor perturbations on neuronal production depend on developmental stage^29,48^. Although these experiments examine different genes and systems, shared gene regulation is accompanied by changes in neural cell production. Together with the relationships between dosage dominance, developmental timing and neuronal–glial fate in our map, these findings suggest that progenitor maintenance and developmental progression are points at which diverse gene functions can converge.

The developmental consequences of altered gene dosage need not scale uniformly across cell-fate transitions^27,49^. In the translation-associated program, stronger KO reversed one modeled excitatory-fate response while amplifying another, despite correlated predicted expression changes between the corresponding WT-fate groups. Experimental dissection of 7q11.23 dosage provides complementary evidence that a gene can affect developmental processes selectively. Lowering *GTF2I* expression in duplication organoids corrected accelerated excitatory-neuron production but had little effect on the imbalance in progenitor production^41^. These observations suggest that distinct components of a dosage-associated developmental phenotype can respond differently to changes in gene expression, allowing correction of one component while others remain altered.

Shared genetic liability can likewise coexist with distinct developmental routes. Common and rare variation have been linked through convergence in genomic regions and coexpression networks^9,10^. Our map places this sharing within developmental programs that also distinguish forms of risk and clinical groups within ASD. A common program may consequently connect different genetic causes without specifying a uniform clinical presentation. The correspondence of selected ASD-associated modules with postmortem cortex extends this interpretation beyond the developmental atlas. Neuronal and glial expression changes recur across psychiatric diagnoses^37,38^, and the contribution of developmental progression to our modeled–postmortem correspondence makes maturation a plausible link between diverse perturbations and similar later transcriptional states. Convergence in cortical gene expression may thus reflect developmental processes shared across genetic backgrounds, even when the earlier effects on cell fate differ. Longitudinal observations would help determine whether these shared expression patterns accompany, precede, or follow the divergence of developmental trajectories.

The perturbation map describes dosage-dependent fate responses that a fixed expression contrast cannot represent. For single-gene fate direction, expression-based scores showed similar correspondence with curated neuronal–glial regulators and a higher correlation with transcription-factor knockdown effects in primary human cortical radial glia. A fixed expression contrast assigns equal and opposite scores to loss and gain of the same gene, whereas the modeled responses show KO–OE asymmetry, changes in response class with perturbation strength and differences across lineage destinations. These responses derive from a dynamics model trained on unperturbed cells, in which each perturbation is an initial shift along a gene-associated direction. Regulatory effects that act differently in distinct lineages without a corresponding expression difference may not be captured by this representation. Cell composition also differed between the experimental and simulated systems. Inhibitory-lineage cells made up 46.6% of control cells in culture, whereas 0.48% of simulated cortical neuronal precursors reached the fetal inhibitory state under WT. Perturbations in matched developmental systems would test whether the modeled changes in lineage choice reflect the regulatory effects of individual genes.

Graded perturbations in cortical and thalamocortical organoids could distinguish changes in differentiation timing from persistent changes in lineage choice^14,50^. Pairing longitudinal expression measurements with fate tracking would test when similar molecular responses produce different effects on neuronal and glial production. Broadly expressed risk genes could be examined alongside lineage regulators, with disease-associated alleles used to connect modeled expression changes to genetic variation. Restoring dosage at defined developmental stages would then test whether normalization of expression also restores cell production, or whether an earlier change in lineage choice persists. Resolving this distinction would identify when a shared transcriptional disturbance remains reversible and when the consequences for neural cell production have already become established.

## Methods

### Human developmental brain single-cell atlas

We used the published human developmental brain single-cell atlas after updating the gene mapping to GENCODE v49/HGNC gene symbols^11^. Other processing followed the published atlas, including preprocessing, scVI batch correction, cell-type annotation, and UMAP embedding. The resulting atlas comprised 393,060 cells and 58,219 genes across developmental stages from gestational day 49 to 92 years old. The atlas log-normalized expression data, batch-corrected scVI embeddings, and UMAP coordinates were used as annotation references and for atlas-level visualization.

### Expanded 21-type cell annotation

Atlas fine clusters within the eight major cell types of the modeled cell-state space were mapped to 21 cell-type labels reflecting regional, laminar and developmental identities. The label mapping integrated the original cluster annotations, developmental and regional composition, canonical marker gene expression, and the top differentially expressed genes (**Supplementary Table 1a**). Radial glia (RG) and neuronal precursors (NP) were subdivided by regional annotation. The original neuroblast clusters were designated as NP, encompassing neuron-committed dividing intermediate progenitors and postmitotic immature neurons. Excitatory neurons were distinguished by developmental stage and laminar or projection-class annotation, and inhibitory neurons by developmental stage and medial or caudal ganglionic eminence (MGE/CGE)-associated identity.

Astrocytes and oligodendrocyte precursor cells (OPCs) retained single labels. Oligodendroglial and microglial subdivisions followed sample-age and developmental-stage composition, with marker profiles used as supporting evidence. Two clusters lacked consistent marker support. They were retained in the modeling state space without an expanded label but were not used as starting or target populations. Thus, 388,455 cells carried one of the 21 labels. The resulting labels were checked for consistency with developmental stage and regional composition and visually inspected in the atlas UMAP embedding.

Detailed annotation criteria and supporting literature are provided in **Supplementary Note 1**, and cluster mappings and evidence summaries are listed in **Supplementary Table 1a**.

### Definition of developmental lineages

We defined a lineage as a specified origin–target combination among the 21 annotated cell types, encompassing either a transition to another cell type or retention of the origin identity. The 21 lineage readouts covered progenitor retention, neurogenesis, gliogenesis, neuronal differentiation and maturation, oligodendrocyte differentiation, and microglial retention. Combinations spanning multiple differentiation stages were not defined as additional readouts.

RG-All pooled RG-Ctx and RG-nonTelen as the origin population, whereas Micro-All pooled fetal and postnatal microglia as both the origin and target populations. Target fate fractions were summed across MGE-type and CGE-type fetal inhibitory neurons for the readout starting from cortical neuronal precursors. Fetal and postnatal microglial target fractions were likewise summed for the pooled microglial retention readout. The microglial readout measured pooled identity retention rather than fetal-to-postnatal maturation.

### scDiffEq modeling state space

For scDiffEq training and perturbation analyses, we constructed a separate modeling state space. Genes detected (raw count > 0) in at least 3% of the 393,060 atlas cells were retained, yielding 14,044 genes. We then excluded endothelial cells (n = 2,095), cells annotated as Others (n = 92) and eight predicted doublets, yielding 390,865 cells. Endothelial cells were excluded to restrict the modeling space to neural and glial lineages. Counts were normalized to 10,000 per cell, log1p-transformed, scaled to unit variance with values clipped at 10, and reduced by principal-component analysis (PCA) to a 50-dimensional scaled-PCA embedding. All scDiffEq training and perturbations used this scaled-PCA embedding. Normalized developmental age, defined as the per-stage median age across the 11 developmental stages after log transformation and min-max scaling (0-1), served as the model time variable.

### scDiffEq model training

We trained scDiffEq^26^, which models single-cell dynamics with a neural stochastic differential equation (SDE) of fixed-potential form, *dX(t) = −∇Ψ(X(t)) dt + σ(X(t)) dW(t)*, where multilayer perceptrons represent the potential *Ψ* and diffusion *σ*, and *W* denotes Brownian motion. The model operated directly on the 50-dimensional scaled-PCA embedding defined above, without an additional autoencoder, and used normalized developmental age as the time variable. The drift network had hidden layers of size 512 x 512, and the diffusion network had hidden layers of size 32 x 32, both with LeakyReLU activations and a dropout rate of 0.1. The model used state-dependent diffusion driven by one-dimensional Brownian motion, formulated in *Itô’s* sense with general noise.

The model was trained on all 390,865 cells using a 90/10 train-validation split, a batch size of 2,048, and no pretraining. Training ran for 50 epochs with RMSprop, an initial learning rate of 1 x 10^-4^, and an SDE integration step of dt = 0.1. The learning rate followed a StepLR schedule with a step size of 10 epochs and a 10-fold decrease at each step. Velocity-ratio regularization was applied with a target ratio of 2.5 and an enforcement weight of 100. The final-epoch checkpoint was used for all downstream analyses.

### Potential landscape and atlas diffusion pseudotime

Potential Ψ was evaluated from the trained model in the 50-dimensional scaled-PCA space and visualized on the atlas UMAP. Representative trajectories were projected to the UMAP using distance-weighted nearest-neighbor regression. Diffusion pseudotime was computed separately from the atlas scVI embedding and was not a scDiffEq output.

Developmental ordering was examined in 275,432 RG, NP, excitatory-neuron and inhibitory-neuron cells, excluding the two unassigned clusters. We constructed a 15-nearest-neighbor graph in the scVI embedding and calculated 15 diffusion-map components using Scanpy^51^. The first diffusion component was excluded when selecting the RG root. Distances from the neuronal centroid were calculated as sums of absolute differences divided by the component standard deviations. Among RG cells at or above the 99th percentile of this distance, the cell nearest that threshold was selected as the root. Diffusion pseudotime was then calculated with Scanpy DPT^52^. Cells were ordered by DPT and divided into 80 equal-count bins. Mean marker expression was summarized in the same bins and smoothed with a seven-bin uniform filter.

### Genome-wide perturbation simulation

For each of the 14,044 genes detected in at least 3% of cells, we simulated knockout and overexpression perturbations. For each gene, a perturbation direction in the 50-dimensional scaled-PCA space was obtained from the gene’s correlation with each principal component, scaled by the per-component standard deviations and divided by the gene’s expression standard deviation. KO and OE were implemented as initial-state shifts along gene-associated directions in scaled-PCA space, corresponding to decreases and increases of two gene-expression standard deviations, respectively. The perturbed cells then evolved under the trained SDE using the Euler-Maruyama scheme for 20 steps (dt = 0.05), with the drift term *−∇Ψ* computed by automatic differentiation. Within each replicate, the wild-type, knockout and overexpression simulations of all genes shared the same Brownian increments for each starting cell. This common-random-number procedure reduced Monte Carlo noise in perturbed–WT comparisons. Terminal states were assigned by k-nearest-neighbor majority vote (k = 20) using Euclidean distance in scaled-PCA space. All 390,865 modeling cells formed the reference, excluding each simulated cell’s starting cell. The vote covered the 21 cell types and, as a separate label, the two unassigned clusters. Ties went to the type of the nearest tied neighbor. For each of 11 origin populations, 500 starting cells were sampled without replacement per replicate. Each gene was simulated in 10 independent replicates.

### Quantification of perturbation effects

For each lineage, the fate fraction was the proportion of the 500 starting cells assigned to the target state or states at the simulation endpoint. For all 21 lineages, Δ was the perturbed minus corresponding same-replicate WT arrival fraction. Positive values therefore indicate increased target-state occupancy, including increased RG retention for the two RG-retention lineages. These retention readouts do not measure proliferation or self-renewal.

Knockout and overexpression deltas across the 21 lineages were concatenated to form a 42-dimensional profile for each gene, averaged over the 10 simulation replicates. Genes with a non-zero delta in at least one profile component were retained as active genes. Lineage-specific KO– OE Pearson correlations were calculated across genes from the 10-replicate mean effects. BCa 95% confidence intervals were estimated using 2,000 bootstrap resamples of the 10 simulation replicates, with the same resamples across lineages. WT floor was defined as a mean arrival fraction ≤0.2% and ceiling as a mean fraction ≥99.8%. Boundary-limited correlations were not interpreted, but all lineages were retained for map and module construction. Pooled KO–OE correlation used the 17 non-boundary readouts and was reported descriptively because genes and readouts recurred across pairs.

### Module construction (two-tier clustering)

In tier 1, the standardized 42-dimensional perturbation profiles (21 knockout and 21 overexpression lineage-transition deltas) were used to construct a k-nearest-neighbor graph (k = 30, exact Euclidean search). Leiden community detection (resolution = 7.0, modularity objective, two iterations, edges weighted by the neighbor-graph connectivities) was run once for each of 1,600 random seeds^53^. For each seed, cluster centroids of the unstandardized profiles were computed across all 42 dimensions and z-scored across clusters. Each cluster was named by the perturbation feature (knockout or overexpression in one lineage) with the largest |z| when this value reached 1.0 and was otherwise labeled non-specific. The name kept the feature, but not the sign of the z-score, so clusters with large positive or large negative deviations on the same feature received the same name. Clusters with the same name were merged into one module, and modules with fewer than 30 genes were labeled as removed. The 1,600 seed-level partitions were combined by consensus. For each gene pair, we computed the fraction of seeds in which both genes carried the same label. Non-specific genes were counted as one group and genes in removed modules as another. Genes were clustered by average linkage on one minus this fraction, and the tree was cut at 0.5. Each consensus cluster took the label that member genes carried most often across seeds. Consensus clusters with the same feature name were merged, and modules with fewer than 30 genes were removed. In tier 2, genes in the consensus non-specific set were partitioned by K-means clustering (k = 4, seed 42, 20 initializations, on profiles standardized within this set). This procedure yielded 31 perturbation-response modules (M1-M31, numbered by size including the four tier-2 modules), and all 14,044 genes were assigned.

### Module annotation and visualization

Module focus was quantified using the focus z-score, defined as the maximum absolute z-score of the mean 42-dimensional module profile, with each dimension standardized across the 31 modules. Tier-1 modules were classified as focused (focus z-score ≥ 2) or intermediate (focus z-score < 2), and the four tier-2 modules as distributed.

Module identities integrated lineage-specific perturbation effects, cell-type expression patterns, and Gene Ontology (GO) biological-process and MSigDB Hallmark enrichment. For each non-diffuse module, the representative transition maximized |zOE − zKO|, where KO and OE module means were standardized separately across the 31 modules for each transition. A positive contrast defined OE-driven, and a negative contrast defined KO-driven. The four diffuse modules received no direction label. This module-level label did not require every gene or transition to share the same direction. Functional enrichment used the 14,044 perturbed genes as the background, with Benjamini–Hochberg adjustment within each module and gene-set collection. Loss-of-function constraint was annotated using gnomAD v4.1 pLI scores^54^, available for 12,021 of 14,044 genes (85.6%). Annotation rules and enrichment procedures are detailed in **Supplementary Note 3**, and module-level annotations are provided in **Supplementary Table 1d**. For the gene-profile UMAP, the 42 axes were standardized across genes. UMAP used Euclidean distance, 30 neighbors, and a minimum-distance parameter of 0.3. Module centroids were unweighted mean perturbation profiles, with each axis standardized across modules. Module similarity was summarized by Ward clustering of these centroids using Euclidean distance. Module-expression summaries averaged gene-wise min–max-scaled cell-type means.

### Gene functional-class annotation

Each gene was assigned to a single functional class, which was used throughout the study as an annotation on the perturbation-derived fate and dosage axes. Gene Ontology (GO) annotations from both biological process and molecular function categories (GOALL) were retrieved using org.Hs.eg.db. A gene was assigned to a class when the gene carried at least one curated term belonging to that class. Genes matching multiple classes were assigned according to a fixed priority hierarchy, ordered from most specific to most general: synaptic > ion channel > chromatin remodeling > transcriptional regulation > RNA processing > ubiquitin/proteostasis > cytoskeleton > cell adhesion > neurogenesis > signaling > cell cycle > immune/inflammation. This priority rule assigned each gene to one class. Remaining genes matching noncoding-RNA symbol patterns (LINC, MIR, SNORD, SNORA, SCARNA, RNU, RN7S, -AS and -DT) or uncharacterized-gene patterns, such as LOC followed by digits, or AC or AL followed by six digits, were grouped under ncRNA. Remaining genes without these patterns were classified as Other. This procedure yielded 12 functional classes, along with the ncRNA and Other categories.

### Module-clustering robustness

The primary module assignment used profiles averaged over 10 simulation replicates, Leiden resolution 7 and consensus across 1,600 seeds. Clustering-seed stability, simulation sensitivity and individual-module recovery are described in **Supplementary Note 2**.

### Developmental trajectory modeling and developmentally dynamic genes

Raw counts from the atlas used for model training were normalized per cell to the median library size across atlas cells and log1p-transformed. Pseudobulk expression was calculated by averaging atlas log-normalized expression within each sample–cluster group containing at least 30 cells. Excluding the two unassigned clusters, RG SOX2 NEBL-1 and Exc SLC17A SNCB, left 983 summaries, with 14,044 modeled genes represented in the reference. Lifespan expression trajectories were modeled for each gene within five maturation cell-type groups: Excitatory (fetal and mature excitatory), Inhibitory (fetal and mature inhibitory), Oligodendroglial (OPC and oligodendrocyte), Astrocytic, and Microglial. Radial glia and neuronal precursors were excluded from these maturation-group trajectories. For each gene and group, expression was fitted against log10 age in days using a generalized additive model (GAM). The model used a cubic B-spline basis with eight quantile-placed knots, an intercept, and a second-difference roughness penalty (λ = 6). Fitted trajectories were evaluated on a 100-point log-age grid. Each pseudobulk summary contributed equally to fitting and developmental-stage comparisons. Knots were placed at age quantiles across all retained pseudobulk summaries.

A gene was classified as developmentally dynamic (devDEG) within a group if (i) the gene was expressed (maximum stage mean > 0.1), (ii) expression differed across the five collapsed developmental stages (Fetal [1st–3rd trimester], Neonatal, Childhood [1–12 years], Adolescence [12–20 years], and Adult [≥ 20 years]) as assessed by a Kruskal–Wallis test (Benjamini–Hochberg FDR < 0.05), and (iii) the dynamic range (maximum − minimum of stage means) exceeded 0.05. Benjamini–Hochberg correction was applied separately to expressed genes within each maturation group.

### Trajectory-class classification and module developmental timing

Each developmentally dynamic trajectory was min–max scaled to 0–1 and classified individually into one of four broad expression patterns described by Herring et al.^55^. We assigned these patterns using an explicit shape rule: transiently rising (an interior maximum exceeding both ends by > 0.15 without a qualifying interior minimum), transiently falling (the converse pattern), rising (end ≥ start), or falling (end < start). Endpoint values determined the rising or falling assignment when neither or both transient criteria were met. Module developmental timing was the percentage of the module’s developmentally dynamic gene–cell-group trajectories classified as rising or transiently rising. Trajectories were pooled across the five maturation groups, so a gene dynamic in multiple groups contributed one trajectory per group; genes without a qualifying trajectory did not enter the denominator.

Six example modules were selected from the 27 lineage-associated modules to represent all four trajectory classes. For each class, modules were ranked by the proportion of their dynamic trajectories assigned to that class. The two highest-ranking modules were shown for rising and falling patterns and the highest-ranking module for each transient pattern. Curves showed within-class medians and interquartile ranges. Class means were shown for all 27 lineage-associated modules, omitting classes that accounted for less than 5% of a module’s trajectories.

### Knockout and overexpression dosage geometry

Module dosage dominance was log_2_[(mean_t mean_g |ΔKO_g,t|/σ_t + 10^-9^)/(mean_t mean_g |ΔOE_g,t|/σ_t + 10^-9^)], using the 21 lineage readouts. Here g indexes module genes, and σ_t was the shared per-lineage WT noise scale: 2√2 times the greater of the sample standard deviation over 30 WT simulations with fixed starting cells and different Brownian increments and the one-cell resolution floor (1/500)/√12. Positive values indicated larger KO response magnitudes and negative values larger OE response magnitudes. This module statistic was a ratio of mean scaled magnitudes, not the mean of gene-level dosage ratios. The association between late-rising trajectory fraction and module dosage dominance was assessed across 27 lineage-associated modules using two-sided Pearson and Spearman tests. The plotted line was an ordinary least-squares (OLS) fit, with a pointwise 95% confidence interval for the fitted mean. The profiles were averaged across ten simulation runs. The four diffuse modules were excluded from the 31-module map for module-level analyses.

### Progenitor-versus-mature module expression dynamics

Because developmental trajectories excluded radial glia and neuronal precursors, we separately summarized module expression along a progenitor-to-mature axis. For each module, mean expression on the pseudobulk reference was computed for progenitor and immature cellular compartments (radial glia, neuronal precursors, and fetal excitatory and inhibitory neurons) and for a mature compartment (excitatory and inhibitory neurons, oligodendrocyte precursor cells, oligodendrocytes, astrocytes, and microglia). The progenitor-to-mature log_2_ expression ratio was computed as log_2_((progenitor mean + 0.01) / (mature mean + 0.01)). This expression ratio was correlated with perturbation-derived dosage dominance across the 27 lineage-associated modules using Pearson correlation to test whether module dosage dominance was associated with the expression of their constituent genes along the progenitor-mature axis. Within each compartment, expression was averaged over all pseudobulk samples and module genes available in the reference after excluding the two unassigned clusters. Pseudobulk samples were weighted equally within each compartment; the gene-level expression ratio instead weighted cell types equally within each group.

### Expression-adjusted functional-class comparisons

Gene-level dosage dominance was log2[(sum_t |ΔKO_g,t|/σ_t + 10^-3^)/(sum_t |ΔOE_g,t|/σ_t + 10^-3^)], where t indexes the 21 lineage readouts and the established per-lineage scale σ_t was shared between KO and OE. Ordinary least-squares models compared functional-class coefficients before adjustment and after inclusion of the progenitor/immature-to-mature expression ratio and mean expression. The ratio was log_2_[(progenitor/immature mean + 0.01)/(mature mean + 0.01)], based on the unweighted means of the 10 progenitor/immature and 11 mature cell-type expression means. The expression-adjusted model included 13 functional-class indicators, the expression ratio, and mean atlas expression, and unclassified genes (Other; 4,746 genes) served as the functional-class reference. Coefficients (β) and ordinary OLS 95% confidence intervals were reported for all 14,044 genes. Genes were the analysis unit. Both models included an intercept. Two-sided t-tests of the class coefficients were corrected jointly across the 26 class coefficients from the unadjusted and adjusted models using Benjamini–Hochberg. Residual dosage dominance in **Extended Data Fig.** 3c was obtained from an expression-only model containing both covariates and no class indicators.

The progenitor/immature group comprised two radial-glial, five neuronal-precursor, fetal excitatory, and MGE- and CGE-derived fetal inhibitory cell types. The mature group comprised three excitatory, two inhibitory, OPC, two oligodendrocyte, astrocyte and two microglial cell types. Mean atlas expression was the average log1p-normalized expression across atlas cells after excluding the two unassigned clusters, RG SOX2 NEBL-1 and Exc SLC17A SNCB, and was entered into the models without further transformation. The association between progenitor-to-mature expression ratio and dosage dominance was assessed using a two-sided Spearman test across all 14,044 genes. For visualization, genes were divided into ten approximately equal-sized groups ranked by expression ratio; ties could be split between groups. Points showed the median expression ratio and median dosage dominance in each group, with the interquartile range of dosage dominance. These summaries did not enter the correlation test.

### KO- and OE-specific neuronal–glial fate changes

KO and OE fate changes were calculated separately relative to WT from 500 radial-glial start cells per run, sampled from pooled cortical and non-telencephalic origins. Summing counts across ten runs gave 5,000 start cells per gene and condition. We summed simulated endpoint counts across astrocyte and OPC states (A), and separately across neuronal-precursor states (N). The latter comprised cortical, ganglionic eminence, thalamic, midbrain–hindbrain and cerebellar neuronal-precursor states.

Gliogenic share was A/(A + N), calculated after pooling counts across runs, and the change from the pooled WT share was expressed in percentage points. Shares with A + N = 0 were undefined; all 14,044 genes had finite pooled shares in the primary analysis. Gliogenic shift was defined as log2[(A + 0.5)/(N + 0.5)] in the perturbed condition minus the same quantity in WT. Positive values indicated a shift toward gliogenic states and negative values toward neuronal-precursor states; the absolute value measured shift magnitude. A pseudocount of 1 was used in a sensitivity analysis. Radial-glial retention, unresolved endpoints, and other destinations did not enter the A + N denominator and were summarized separately. Destination-count summaries used the median across genes within each dosage-dominance decile, based on ten-run pooled counts.

### Fate–dosage dominance associations

We used two-sided Spearman tests to relate gene dosage dominance to gliogenic share change and to absolute gliogenic shift, analyzing KO and OE separately. Gene-level 95% confidence intervals were the 2.5th and 97.5th percentiles of 2,000 bootstrap estimates obtained by sampling genes with replacement, and the same sampled indices were used for KO and OE. Module shifts were medians across member genes, with percentile intervals from 2,000 within-module gene bootstrap samples. A module’s shift direction was assigned when the bootstrap interval excluded zero. Module-level share-change tests used means across genes in each of the 27 modules,

Spearman correlations, and bootstrap resampling of modules. *P* values for absolute-shift correlations were unadjusted. Benjamini–Hochberg correction covered the four share-change tests (gene/module x KO/OE); paired bootstrap estimates of ρOE − ρKO were used to compare the correlations. Resampling used NumPy’s default random-number generator with seed 20261110.

To separate within-module associations from differences between modules, we restricted analysis to the 8,490 genes in the 27 lineage-associated modules. Dominance and share change were ranked across these genes with average ranks for ties, and each gene’s module-specific mean rank was subtracted. Pearson correlation between the centered ranks quantified the within-module association. Confidence intervals used 2,000 bootstrap samples drawn within each module, with ranks recalculated in every sample. KO and OE were analyzed separately. Variability across simulation replicates and sensitivity to lineage selection are described in **Supplementary Note 4**.

### Gene-set and regulator comparisons of fate direction

Five GO biological-process sets were defined using org.Hs.eg.db 3.20.0, including descendant terms: positive and negative regulation of gliogenesis (GO:0014015 and GO:0014014), positive and negative regulation of neuron differentiation (GO:0045666 and GO:0045665), and Notch signaling (GO:0007219). Within the 14,044 modeled genes, each set was compared with all remaining genes separately under KO and OE using two-sided Mann–Whitney U tests. AUROC was calculated as U/(n_set x n_background), with values >0.5 indicating higher gliogenic shifts in the set. Benjamini–Hochberg correction covered the ten set-by-condition comparisons. The same tests were applied after excluding M30 and M10 from both groups and to WT-relative changes in neuronal-precursor endpoint counts pooled across ten simulation replicates (**Extended Data Fig.** 3e), and these additional comparisons reported unadjusted *P* values.

For an expression-only reference, dE was the unweighted mean log1p-normalized expression across astrocyte and OPC states minus that across the five neuronal-precursor types. Scores of −dE for KO and dE for OE were evaluated using the same gene-set comparisons. An astrocyte-only reference assessed sensitivity to glial weighting. Each reference used a separate ten-test Benjamini–Hochberg correction family.

Published loss- and gain-of-function effects on neuronal versus glial output were curated from abstracts and relevant full-text results without restriction on brain region. This selected literature set was not a systematic review. Unambiguous loss-of-function directions were compared with modeled KO shifts for 25 regulators. Reported gain-of-function directions were compared with OE shifts. Modeled shifts with absolute magnitude <0.5 were classified as small; those with absolute magnitude ≥0.5 were classified as matching or opposite to the reported direction. Concordance was descriptive, and jointly perturbed genes shared a study identifier. A separate comparison applied the same raw-sign rule to modeled shifts and expression-reference scores, without magnitude thresholds; zero was not counted as a match. Sources, experimental systems, phenotypes, single-gene, combined-perturbation or indirect evidence, and comparison results are provided in **Supplementary Table 2g**.

For the primary-cell CRISPRi comparison, we used day-7 cells from three 2D libraries in GSE284197^29^. We retained cells carrying non-targeting sgRNAs or guides against a single transcription factor, excluding technical, microglial and vascular cells and cells carrying only inactive guides (18 guides with minimum cell-class log2 fold change > −0.4). Inhibitory (IPC_IN, IN) and excitatory (IPC_EN, EN_ImN, EN) lineage counts were compared with non-targeting controls using Mantel–Haenszel log odds ratios across nine donor-by-library strata. Experimental effects were tested using Cochran–Mantel–Haenszel tests with Benjamini–Hochberg correction across 44 factors per readout. Modeled effects were KO-minus-WT changes in the natural log ratio of fetal inhibitory to combined fetal excitatory and Exc-L5/6 non-IT endpoint counts from cortical neuronal precursors, pooled across ten simulation replicates with 0.5 added to each count.

The expression-only KO reference was atlas mean expression in fetal excitatory minus fetal inhibitory neurons. Comparisons used the 43 transcription factors represented in the perturbation map. Spearman correlations used 10,000 factor-label permutations (one-sided) and 10,000 factor-bootstrap resamples for percentile 95% confidence intervals. Partial Spearman correlation was calculated between model and experimental residual ranks after adjustment for reference ranks. For the partial-correlation test, model scores were permuted across factors and residuals recalculated in each of 10,000 permutations (one-sided). Directional concordance was summarized from matching signs across all 43 factors and separately among factors with experimental BH-adjusted *P* < 0.05. No magnitude threshold was applied, and zero-valued scores were not counted as matches.

We compared linear predictors of experimental knockdown effects using leave-one-transcription-factor-out cross-validation across the 43 factors. OLS models with an intercept used atlas mean expression in fetal excitatory and fetal inhibitory neurons, the modeled KO shift, or all three predictors. Expression means were unweighted averages across the corresponding pseudobulk groups. Each factor was predicted using models fitted to the other 42, with their mean experimental effect as a control. Predictions were evaluated by mean squared error (MSE). Relative MSE was the ratio of each model’s MSE to the control MSE. The added contribution of modeled KO shifts was evaluated as MSE(expression regression) − MSE(combined regression). Percentile 95% confidence intervals were estimated from 2,000 factor-level bootstrap resamples, repeating cross-validation with multiplicity weights and excluding all copies of the evaluated factor from training. Experimental effect estimates were held fixed.

### Strength-dependent lineage response classification

We used the mean fate shifts across the 10 simulation replicates at 0.5, 1, 2 and 4 gene-expression s.d. in both KO and OE directions (arms). The ±2 s.d. module assignments were held fixed across strengths. Fate shifts were expressed as percentage-point differences in endpoint target-state occupancy relative to matched WT simulations. An arm was evaluable if at least two strengths had an absolute effect of at least 1 percentage point. A sign-shift response required an evaluable arm with effects of at least +1 and at most −1 percentage point. To detect unstable responses, each arm was oriented by the sign of the largest absolute effect in that arm; a decrease between any lower and higher strength of at least the larger of 2 percentage points or 10% of that arm’s maximum absolute effect defined an unstable response. Remaining evaluable curves were classified as stable. Curves with the same dominant sign in both evaluable KO and OE arms formed a separate category. Sign-shift took precedence over this category, which in turn took precedence over unstable responses. Curves with a maximum absolute effect below 1 percentage point or insufficient evaluable points were retained separately. Module- and lineage-level composition summaries were restricted to the 27 lineage-associated modules. Percentages were calculated among curves classified as stable, unstable or sign-shift, excluding same-sign, small-effect and insufficiently evaluable curves. Lineage-level summaries counted genes, whereas module-level summaries counted gene–lineage pairs.

### Destination discordance and functional enrichment

For each origin with at least two destinations, genes were evaluable if at least two destination curves had stable, unstable or sign-shift response types. A gene was discordant if at least two evaluable destinations had different response types. Each gene was counted once per origin. Module breadth was the number of modules containing at least a given number of discordant genes. Origin-level counts used all 31 module assignments. The cortical-precursor enrichment background comprised evaluable genes in the 27 lineage-associated modules, excluding the four distributed modules, after gene-symbol harmonization. We compared each gene set with the remaining genes in this background using a one-sided Fisher’s exact test. Gene Ontology biological-process annotations were mapped to broader categories in the generic GO-slim subset using subtype and part–whole relationships. Only non-obsolete categories were retained. Current symbols and uniquely mapped previous or alias symbols were resolved using HGNC. Gene sets with at least five evaluable members and at least five evaluable nonmembers were tested. Benjamini–Hochberg correction was applied jointly across the 145 tested GO-slim, transcription-factor, developmental-signature, perturbation-screen and disease sets. Transcription-factor sets comprised DNA-binding-domain families and motif-linked transcription factors from Trevino et al. ^56^ and inferred cortical-lineage regulators from Velmeshev et al. ^57^. Developmental-signature sets were lineage developmental genes from the same study, in which the Ex2 branch gives rise to L5 and L6 excitatory neurons. Perturbation-screen sets comprised the 244 regulators screened by Ziller et al. ^58^ and subsets defined by their effects in that screen. The NDD set comprised the 497 genes with NDD TADA FDR < 0.001 in Satterstrom et al. ^30^, and other disease sets followed the definitions used for module disease enrichment. M24 follow-up genes were the intersection of M24 membership, cytoplasmic-translation annotation and destination discordance.

### WT-fixed downstream expression responses

Starting cells were grouped by their destination in the matched WT simulation (WT-fate groups). Group membership was held fixed across perturbation directions and strengths. Within each group and simulation replicate, we calculated the difference between the mean perturbed and WT endpoints in the 50-dimensional modeling space. We projected this difference through the corresponding PCA loadings to reconstruct gene-expression changes. The projected initial-state displacement was then subtracted to obtain the input-subtracted response. These responses were expressed in the model’s standardized gene-expression coordinates. The WT-to-2 s.d. contrast was the input-subtracted response at 2 s.d. The 2-to-4 s.d. contrast was the difference between the responses at 4 and 2 s.d. within each replicate. WT-empty groups were treated as missing. Estimates were averaged across the 10 paired simulation replicates.

### Comparison of *NFIA* downstream responses with experimental NFI loss

We used the eight transcription factors highlighted in the developmental-stage-specific *Nfia*/*Nfix* loss-of-function comparisons reported by Zhang et al. ^31^: Rest, Hmga2, Fubp1 and Creb1 at E13.5, and Brca1, Foxm1, Zeb1 and Tead1 at E15.5. Mouse symbols were matched to uniquely mapped human orthologs and harmonized to the modeled target genes. FOXM1 was absent from the modeled target set, leaving seven factors. Published changes in the combined *Nfia*/*Nfix* knockout provided the reference sign for the modeled *NFIA* KO response. We compared these signs with the mean input-subtracted response at 2 s.d. in the fetal excitatory and layer 5/6 non-IT WT-fate groups, then computed the paired additional changes from 2 to 4 s.d. The embryonic-stage labels identify the source experimental comparisons; the two modeled WT-fate groups were not assigned to those embryonic ages. Directional agreement was reported descriptively for the prespecified factors.

### Translation-module fate and downstream comparisons

The same 73 M24 genes were used for the fate curves and the downstream comparison. Fate curves retained all measured KO and OE strengths from 0.5 to 4 s.d.; group summaries were medians across genes. For the downstream comparison, we reconstructed the input-subtracted KO responses in the two excitatory WT-fate groups and calculated each target’s 2-to-4 s.d. change. All 100 genes in the original translation-perturbation input set were excluded from the downstream target universe. Responses were first averaged across model replicates and then equally across the 73 perturbation genes, giving one mean response per target and destination. Agreement between the two groups was summarized as the Pearson correlation across targets, for the perturbation-averaged responses and separately for each perturbation. For each KO perturbation, fate divergence was the additional percentage-point change from 2 to 4 s.d. toward layer 5/6 non-IT excitatory states minus the corresponding change toward fetal excitatory states. The association between fate divergence and the target-level Pearson correlation was assessed using a two-sided Spearman correlation test across the 73 perturbations. No perturbation subgroups were defined, and the comparisons were summarized descriptively.

### Brain-disease risk gene sets

The primary ASD set comprised 290 genes from Satterstrom 2026 overall TADA (FDR < 0.001) or the curated EAGLE list^30,59^. Corresponding clinical-stratum TADA analyses (FDR < 0.05) or EAGLE yielded 376 ID/DD and 271 no-ID/DD genes, kept distinct from the primary set. The NDD set comprised 497 genes (478 mapped); the DD set comprised 309 genes with DD TADA FDR < 0.001 in Fu et al.^1^ (303 mapped). Current, previous, and unambiguous HGNC aliases matched symbols to the 14,044 modeled genes.

AD genes were predicted effectors from Uffelmann et al.^32^, Supplementary Data 2: 190 unique genes at 127 loci (157 mapped), excluding generic HLA, MS4A and IGH labels. Schizophrenia genes had combined SCHEMA case–control *P* < 0.01 in Singh et al. ^60^. GRCh37 Ensembl identifiers were converted with GENCODE 19 before HGNC matching (244 genes; 209 mapped). Parkinson’s disease, major depressive disorder, bipolar disorder, and epilepsy lists were from the GWAS Catalog, Meng et al. ^61^, Palmer et al. ^62^, and Epi25^63^, respectively. ClinVar pathogenic or likely pathogenic records were deduplicated by variant identifier, excluding conflicting interpretations (February 2026 variant summary; 18 February VCF)^64^. Mondo descendants and OMIM equivalences (6 October 2026 release) defined intellectual disability (MONDO:0001071), leukodystrophy (0019046), developmental and epileptic encephalopathy (0100620) and hereditary ataxia (0100309)^65^. Microcephaly (MONDO:0001149) and lissencephaly (0018838) were defined as separate gene sets. Multigene records were assigned only to the gene named in the HGVS description. Intellectual disability genes overlapping any of the three ASD sets were removed.

### Module enrichment and disease comparisons

Disease–module enrichment used one-sided Fisher’s exact tests against remaining modeled genes, including those in distributed modules. Benjamini–Hochberg correction covered 15 sets x 27 lineage-associated modules, and separately 27 modules each for microcephaly and lissencephaly. Enrichment required OR > 1 and q < 0.05; ASD-enriched modules were significant for at least one of the three ASD sets. Covariate-matched enrichment used 1,000 disease-label permutations within expression, pLI and input-displacement quintiles, with upper-tail plus-one *P* values and Benjamini–Hochberg correction across 34 associations.

After excluding shared genes, disease pairs were compared by total variation distance across 31 modules using 10,000 label permutations within expression–pLI strata containing both groups. Holm correction covered four pairs (**Supplementary Table 4c**).

### Disease-associated lineage responses and enrichment coverage

Per-gene signed responses were ΔT/ΣT|ΔT| and magnitude responses were |ΔT|/ΣT|ΔT|. Normalization was performed separately for KO and OE. Zero-total-effect profiles remained zero. Disease means were compared with 5,000 equal-sized null sets sampled with replacement within expression–pLI tertiles, separating missing pLI. Response z-scores were observed minus null means, divided by the null standard deviation. Empirical tests were upper tailed for magnitude and two-sided about the null mean for signed responses. Benjamini–Hochberg correction covered 17 sets x 21 transitions x two directions per endpoint.

Module mean |Δ| was standardized against the genome-wide mean and standard deviation per transition and direction. The larger KO or OE z-score ranked transitions within each module. A significant disease response was covered if the corresponding transition ranked among the top three of any disease-enriched modules, irrespective of direction. A random-placement reference independently assigned covered status with probability equal to each disease’s covered-transition count divided by 21 (10,000 draws; two-sided empirical test).

For representative disease–module pairs, each module’s defining transition and direction maximized the absolute genome-standardized module z-score. Disease transitions were ranked by their smaller KO or OE normal-approximation *P* value. We selected the lowest-q enriched module whose defining transition ranked among the top three; otherwise, the highest-OR module defined by the leading transition. Significance used the empirical disease-response tests above. Module expression in relevant cell types was weighted by each gene’s absolute defining-transition effect, averaged and z-scored across seven stages: early fetal (first–second trimester), late fetal (third trimester), neonatal, infancy (1–6 yr), childhood (6–12 yr), adolescence (12–20 yr) and adult (≥20 yr). Complementary curves used equal gene weights, donor-first stage averages or Herring data alone.

### Developmental timing comparisons among neurodevelopmental risk-gene sets

Module curves summarized previously inferred gene expression trajectories as pointwise medians on the common 100-point log-age grid, rescaled to each module’s range. Timing was the fraction of developmentally dynamic gene–cell-group trajectories classified as rising or transiently rising, calculated within the assigned lineage-associated module or individual gene. Comparisons used nonoverlapping ASD and DD genes and exclusive ASD clinical-stratum genes, excluding missing trajectory estimates.

Genes were the comparison unit. Unadjusted contrasts were summarized by mean differences and Cliff’s delta, with 95% percentile CIs for Cliff’s delta from 2,000 independent within-group gene bootstrap samples. Regression included all genes with timing estimates, group indicators, and natural cubic splines for expression percentile and pLI (5 and 3 degrees of freedom). Missing pLI was median imputed with a missingness indicator; 95% CIs used HC3 standard errors. The four regression contrasts are reported with unadjusted two-sided *P* values (**Supplementary Table 4e**).

### Functional and clinical transition bias

Per-gene magnitudes were |Δ| divided by the per-lineage WT noise scale σ_t_ defined above, separately for KO and OE. Within primary ASD genes, Satterstrom GO group 3 defined transcriptional regulation and groups 15 and 21 synapse formation and organization. Excluding overlaps left 57 transcriptional and 43 synaptic genes. Clinical comparisons used 213 ID/DD-only and 108 no-ID/DD-only genes without restriction to primary ASD.

The early–late index subtracted the mean magnitude across nine neuronal differentiation and maturation transitions from that across nine progenitor retention, neurogenesis and gliogenesis transitions. Transcriptional-minus-synaptic and ID/DD-minus-no-ID/DD contrasts used one-sided tests with 10,000 label permutations within expression ventiles crossed with pLI quintiles (separating missing pLI), or within modules alone. Holm correction covered two contrasts x two directions per control family. Transition-specific contrasts used 2,000 expression–pLI permutations and two-sided tests, with Benjamini–Hochberg correction across 21 transitions per contrast and direction.

### Rare risk-gene dosage geometry

KO and OE deltas were z-standardized separately along each lineage axis across all 14,044 genes, using population standard deviations. Bidirectional magnitude was the geometric mean of 21-dimensional KO and OE Euclidean lengths. Neurogenic OE drive averaged signed OE z across ten neurogenesis and neuronal-differentiation transitions. Initial perturbation size was the Euclidean norm of the input displacement in PC space. ASD and NDD were compared with 10,000 expression–pLI-matched draws, with or without matching for this size. Input matching crossed 30 expression, five pLI, and five displacement bins, excluding genes without matched support.

Nearest-neighbor controls used ten nonmembers per gene by Mahalanobis distance on these covariates, matching missing-pLI status. Matched AUROC ranges were drawn percentiles (2.5– 97.5%); Holm correction covered two sets x two metrics per control family.

### Convergence of perturbation signatures with postmortem ASD cortex

KO and OE signatures pooled 500 endpoint cells per starting type (11 types, 20 steps), with paired WT starting cells and Brownian paths. Mean perturbation–WT differences in 50-dimensional principal component space were projected into gene expression using PCA loadings, then averaged across ten simulations with equal starting-type weights. The target background comprised modeled genes represented in postmortem sets after applying a gene-symbol filter for housekeeping, ribosomal and mitochondrial genes. The filter excluded symbols beginning with RPL, RPS, MRPL, MRPS, MT-, TUBA, TUBB, ACTB, ACTG, EEF, EIF, HSP, FTL, FTH1, TMSB, GAPDH, MALAT1, NEAT1, PTMA, VIM, UBC, UBB, FAU, NACA or TPT1, and the symbols *MTRNR2L8* and *MTRNR2L12* The same filter was used for the cortical specificity analyses. Postmortem ASD gene sets came from Parikshak et al.^15^, both Gandal 2018 resources^16,37^, Velmeshev et al.^17^ and Wamsley et al.^18^. Parikshak sets included FDR < 0.05 differentially expressed genes and modules M19 (upregulated), M16 and M10 (downregulated). Gandal sets included full CD11 and CD4 memberships (upregulated), CD1 and CD13 (downregulated), and isoform-study genes with ASD FDR < 0.05. Velmeshev sets used q < 0.05, or FDR < 0.05 for bulk-deconvolved glia. Wamsley sets required |log2 fold change| > 0.25, FDR < 0.05, and ≥30 genes; oligodendrocyte and OPC sets came only from this study. Differentially expressed genes were separated by direction. After spreadsheet-symbol restoration and HGNC matching, unions by cell class and direction combined 72 sets into 18 groups.

Competitive scores for perturbed gene g and postmortem set S were z(g,S) = (mean_S γ_g − mean_U γ_g)/(SD_U γ_g/√n_S), where γ_g is the signature, U the target background, SD_U the population standard deviation of the target background, and n_S the retained set size. Stouffer’s method, Σz/√n, combined scores across primary ASD genes in M14 (10 genes), M22 (18) and M25 (13). Comparators were 5,000 sets of non-ASD genes sampled without replacement within expression, pLI and signature-norm quintiles. Empirical tails followed postmortem direction: upper for upregulated sets, lower for downregulated sets. Benjamini–Hochberg correction covered 216 individual-set and 54 grouped comparisons separately within each perturbation direction.

### Cross-disorder specificity of cortical expression concordance

For Gandal cortical contrasts^16,37^, each KO signature was standardized across atlas targets before averaging across all genes per module. Spearman correlations compared module signatures with disease-associated changes across shared targets, excluding housekeeping, ribosomal, and mitochondrial genes. ASD-specific residuals were controlled for schizophrenia, bipolar disorder, major depressive disorder, and alcohol-abuse/dependence in microarray data, or schizophrenia and bipolar disorder in RNA-seq. Mean concordance of the three ASD-associated modules was ranked among all 2,925 triplets from 27 modules using upper-tail tests. Progression adjustment regressed concordance on dosage dominance and late-rising fraction in the other 24 modules for each triplet.

Per-gene concordance compared risk genes with nonmembers using 10,000 one-sided label permutations within expression–pLI tertiles, separating missing pLI. Benjamini–Hochberg correction covered 132 tests. Stratum-specific Cliff’s delta values were weighted by risk-gene counts across 11 cortical contrasts.

### CNV-supported gene definition and dosage-response comparisons

CNV evidence was obtained from Fu et al.^1^. Genes with positive log10 Bayes factors for deletion, duplication, or both were classified as deletion-supported, duplication-supported, or bidirectionally supported, respectively (missing values were set to 0). The scored gene universe comprised 11,312 genes shared by the perturbation map and the gene-level CNV evidence table, of which 202 were CNV-supported. Gene-level bidirectional effect magnitude was sqrt[(Σ_t |ΔKO_t|)(Σ_t |ΔOE_t|)], summed over all 21 lineage readouts of the ten-replicate mean profiles. The 202 CNV-supported genes were compared with the remaining 11,110 scored genes using a one-sided Mann–Whitney U test for larger magnitudes in CNV-supported genes. AUROC was U divided by the product of group sizes. For the dosage-matched gene heatmap, deletion-supported genes were ranked by Σ_t |ΔKO_t| and duplication-supported genes by Σ_t |ΔOE_t|, with ties broken by gene name. The five highest-ranked genes from each arm were selected, with bidirectionally supported genes eligible for both rankings. Selection stability was assessed by applying the same rule to each of the ten single-replicate profiles (**Supplementary Table 5c**).

### CNV-arm dosage-matched contribution

Recurrent genomic-disorder CNV regions and their deletion and duplication arms were taken from Fu et al. ^1^, and genes were assigned to arms by GENCODE v49 coordinates. All 79 arms contained perturbed genes, giving 1,426 gene–arm pairs (**Supplementary Table 5g**). Deletion arms were evaluated using KO responses and duplication arms using OE responses. For each gene, the dosage-matched score was the maximum absolute response across the 21 lineage readouts. The contribution share was this score divided by the sum of scores across all modeled genes in the same arm. This normalized score summarized the relative response magnitudes of individually perturbed genes within each arm. Reciprocal regions were pairs of deletion and duplication arms, each containing at least two genes, whose gene sets had a Jaccard index ≥0.5 (31 pairs). For each pair, we recorded the top-contributing gene, the contribution share and the margin over the second-ranked gene in each arm (**Supplementary Table 5d**).

### Comparison with gene-level CNV dosage experiments

A targeted, non-systematic literature review identified gene-level dosage experiments for three of the 31 reciprocal CNV regions: 15q13.2–q13.3, with gene re-expression in Df(h15q13)/+ cortical neurons^40^; 7q11.23, with Williams–Beuren syndrome and 7q11.23 duplication cortical organoids with *GTF2I* knockdown, and *Gtf2i*+/− and *Gtf2i*+/Dup mice^41^; and 16p11.2, with *kctd13* knockdown and *KCTD13* overexpression in zebrafish, and *Kctd13* knockdown in mouse cortex^42^. For 15q13.2–q13.3, rescue of dendritic spine density and dendrite outgrowth by each tested gene was recorded as reported. For 7q11.23, the 17 region genes in the map were grouped by responses from cortical neuronal precursors to fetal excitatory neurons into OE ≥2 pp and KO ≤−2 pp, the reverse, or neither. Primary model comparisons used the 10-replicate mean responses at ±2 gene-expression s.d.

Each reported finding of Lopez-Tobon et al. ^41^ (15 findings) and Golzio et al. ^42^ (5 findings) was matched to a model quantity, a lineage readout and an expected sign. Loss of gene or region dosage was compared with KO, gain with OE, and deletion-versus-duplication contrasts with KO – OE. Phenotypes removed by *GTF2I* knockdown in duplication organoids were compared with OE, with the sign of the duplication phenotype. A phenotype unchanged by knockdown was expected to show no OE response. Progenitor proliferation or abundance was matched to cortical radial-glial retention as an indirect proxy, excitatory neuron number or maturation to transitions from cortical neuronal precursors to fetal excitatory neurons, deep-layer neurons to transitions from cortical neuronal precursors to layer 5/6 non-IT excitatory neurons and upper-layer neurons to transitions from fetal excitatory neurons to layer 2/3 IT excitatory neurons. Intermediate progenitors had no matching readout. A finding was consistent when the model response had the expected sign with absolute value ≥1 pp, inconsistent when the sign was opposite with absolute value ≥1 pp, and below threshold otherwise. When no change was expected, an absolute response <1 pp was consistent. Thresholds of 0.5 and 2 pp were used in sensitivity analyses. Directional concordance was descriptive. Findings mapped to the same model response were not treated as independent tests. Sources, systems, phenotypes and model values are listed in **Supplementary Table 5f**.

### Ethics statement and description of the ASD sequencing consortium cohort

This study was conducted in accordance with the ethical principles outlined in the Declaration of Helsinki. All procedures involving human genomic data were approved by the Institutional Review Board of Seoul National University Bundang Hospital (IRB No. B-1703-388-303 and B-2108-700-107), Soon Chun Hyang University Hospital Bucheon (SCHBC; IRB No. SCHBC 2018-04-020 and SCHBC 2022-04-016), and Seoul Child Hospital (IRB No. P01-201908-BM-02 and P01-202111–21-003). Informed consent was obtained from all participants or their legal guardians. No financial or material compensation was provided for this study. The respective IRBs approved the collection, storage, and analysis of biospecimens and genetic data. All data were handled in accordance with the relevant data protection and privacy regulations.

We integrated whole-exome (WES) and whole-genome (WGS) sequencing data from four ASD family cohorts, each composed of probands, parents, and unaffected siblings. The WES data comprised a Korean autism cohort (252 individuals: 73 probands, 124 parents, 55 unaffected siblings)^66^ and SPARK v2 (54,900 individuals: 17,482 probands, 30,544 parents, 6,874 unaffected siblings)^67^. The WGS data comprised the Korean cohort (3,823 individuals: 1,160 probands, 2,286 parents, 377 siblings)^66^, SPARK v1.1 (12,396 individuals: 3,499 probands, 6,708 parents, 2,189 siblings)^67^, the Simons Simplex Collection (SSC; v05-12-2019; 9,082 individuals: 2,380 probands, 4,764 parents, 1,938 siblings)^68^ and MSSNG db7 (9,463 individuals: 3,336 probands, 5,716 parents, 411 siblings)^69^. SPARK and SSC data were obtained through SFARI Base, and MSSNG data through the Autism Speaks MSSNG resource. After sample-level quality control, the integrated family-based resource comprised 89,916 samples from 25,071 families. The original de novo analysis resource comprised 39,774 sequenced probands and siblings with parent–child data sufficient for de novo variant identification. After excluding 10 SSC offspring following consent withdrawal and one SPARK offspring sequenced only in a later SPARK release, and correcting diagnostic labels for 47 SPARK whole-genome-sequenced and 176 SPARK exome-sequenced offspring, the de novo analyses included 39,763 offspring: 28,147 ASD probands and 11,616 unaffected siblings used as controls.

### Case-control burden enrichment

Variants annotated as escaping nonsense-mediated decay were excluded before all de novo variant analyses. For each gene set and variant class, ASD probands were compared with unaffected siblings on a per-carrier basis. Gene sets included genome-wide genes, ASD risk genes, complementary non-ASD genes, and each fate module. Carriers had at least one DNV of the tested class in the tested gene set. Carrier and noncarrier counts formed a 2 x 2 table. Relative risk and Wald 95% confidence intervals were reported when both groups had at least one carrier. We obtained two-sided Fisher’s exact *P* values from raw carrier counts. BH correction was applied jointly across the 259 gene-set-by-variant-class comparisons with at least one carrier in either group. The 11 comparisons with no carriers in either group were not tested. Protein-truncating variants were stratified by gene constraint using gnomAD LOEUF deciles, and damaging missense variants were stratified by MPC score. The Damaging category combined PTVs in the first LOEUF decile and missense variants with MPC ≥ 2. No additional population allele-count filter was applied to the burden analysis.

In parallel, we reported paternal-age-adjusted mean DNV rates per individual. Genome-wide DNV count was regressed on paternal age separately for WGS and WES, and residuals were recentered to the observed mean. Each individual’s rate contribution was weighted by the adjusted-to-observed count ratio. Individuals with missing paternal age retained a weight of one. Carrier counts used for relative-risk estimation and Fisher’s exact tests were left unweighted. Enrichments were computed in the combined cohort.

### Fate-axis annotation for carrier comparisons

For the carrier phenotype comparisons, genes were assigned a fate-axis score F = (g − n)/(|g| + |n| + 0.3), where g was the mean OE response across glial transitions and n was the mean across neuronal transitions. OE responses were standardized across genes separately for each transition. The glial group comprised RG → OPC, RG → Astro, OPC → NFOL and NFOL → MOL; the neuronal group comprised the 14 neurogenic and neuronal-maturation transitions. The two radial-glial retention transitions and the microglial transition were excluded. Genes with positive scores were classified as gliogenic and genes with negative scores as neurogenic. This OE-based annotation was distinct from the condition-specific neuronal–glial shifts used for the dosage-dominance analyses.

### Within-case gene and variant analyses

Gene-level and phenotype analyses used PTVs in the first LOEUF decile and missense variants with MPC ≥ 2 in ASD probands, excluding variants with an All of Us allele count greater than five. Variants with missing allele counts were retained. Damaging DNV genes were classified as PTV-only, missense-only, or shared. Shared genes carried at least one damaging PTV and at least one damaging missense variant among ASD probands. PTV-only genes carried damaging PTVs but no damaging missense variants, whereas missense-only genes carried damaging missense variants but no damaging PTVs among ASD probands. Enrichment of established ASD risk genes, loss-intolerant genes, and ASD-enriched fate-module assignment was tested in shared-hit genes against the other two classes combined using one-sided Fisher’s exact tests. Missing annotations were counted as not meeting the criterion for these tests. Loss intolerance was defined as pLI > 0.9.

Among model-represented genes hit by damaging missense variants (MPC ≥ 2) in ASD probands, continuous dosage dominance was compared between synaptic and ion-channel genes and chromatin-remodeling and transcriptional-regulatory genes using a two-sided Mann–Whitney U test and Cliff’s δ. The same comparison was made across all model genes in these functional classes. Specificity to damaging-missense genes was tested by regressing percentile-ranked dosage dominance on functional class, damaging-missense gene membership, and their interaction (**Extended Data Fig. 7d**).

### Perturbation-profile convergence with ASD risk genes

For each gene, the perturbation profile was represented by the concatenation of knockout and overexpression effects across the retained lineage transitions. Fate-module convergence to ASD risk genes was defined as the mean cosine similarity between a gene’s perturbation profile and those of the curated ASD risk gene set, computed using a leave-one-out procedure for genes in the risk set. Co-expression convergence was computed analogously from cell-type mean expression profiles in the primary developmental atlas.

To test whether fate-module convergence carried information beyond gene expression and evolutionary constraint, we fitted logistic regression models to predict membership in each target gene set using pLI, co-expression convergence, and fate-module convergence. Target gene sets were curated ASD risk genes and genes recurrently hit by damaging de novo variants. Missing pLI values were replaced by the median, and the three predictors were standardized to zero mean and unit variance before fitting. Odds ratios therefore represent a one-s.d. increase in the corresponding convergence score. The added contribution of fate-module convergence was evaluated with a likelihood-ratio test comparing models with and without the fate-module convergence term, conditional on pLI and co-expression convergence. For the two convergence tests in Supplementary Table 6a, recurrently hit genes had at least two damaging de novo events. Their ASD reference excluded recurrently hit genes, leaving 126 reference genes. Empirical significance of the added likelihood ratio was assessed with 10,000 label permutations within 125 strata defined by quintiles of pLI, mean expression and perturbation input magnitude. Reference sets and convergence scores were recomputed for each permutation before refitting the models. Empirical *P* values were (exceedances + 1)/(10,000 + 1) and were BH-adjusted across the two convergence tests.

For the case–sibling comparison, damaging de novo variants observed in ASD probands, and unaffected siblings were mapped to their carrier genes and fate modules. The fraction of pooled damaging de novo events in ASD-enriched fate modules was compared between probands and siblings using a two-sided Fisher’s exact test (**Extended Data Fig. 7b**).

### Phenotype association

Among ASD probands, carriers of the stringent damaging DNV set were compared with non-carriers, with 12 carrier groups defined by ASD-risk-gene status, variant-type gene group, fate module, and fate-axis annotation. As a specificity control, nine carrier groups were also compared with all other damaging DNV carriers. Eighteen clinical and developmental measures were analyzed, spanning autism symptoms, social communication, restricted and repetitive behavior, cognition, adaptive function, and motor coordination. Each measure was z-score normalized across ASD probands with an observed score. For each carrier group and phenotype, a sex-adjusted linear model (*z-score ∼ carrier + sex*) was fitted, and the standardized carrier coefficient was used as the effect size. Groups with ten or fewer carriers, a single sex, or no phenotypic variance were not tested. Benjamini–Hochberg correction was applied separately across the 209 tested group–phenotype comparisons against non-carriers and the 155 comparisons against other damaging-DNV carriers. Group definitions, carrier counts, comparator counts and phenotype missingness are reported in **Supplementary Table 6f**.

Overall, autism symptom severity was measured using the Autism Diagnostic Observation Schedule-2 (ADOS-2) calibrated severity score^70,71^ and the current and lifetime SCQ score^72^. Social-communication traits were measured using the Social Responsiveness scale (SRS) ^73,74^, the ADOS-2 Social Affect (SA), and domains A (social interaction) and B (communication) of the Autism Diagnostic Interview-Revised (ADI-R)^75^. Restricted and repetitive behavior was measured using the ADOS-2 Restricted and Repetitive Behavior (RRB), ADI-R domain C, and the Repetitive Behavior Scale-Revised (RBS-R)^76^. Cognitive and adaptive measures included the full-scale IQ (FSIQ)^77,78^, nonverbal IQ, the Vineland Adaptive Behavior Scales (VABS)^79^ scores across communication, daily living, socialization, and motor skills, and the Developmental Coordination Disorder Questionnaire (DCDQ)^80^. SPARK scores came from the 25 June 2026 release. We used official ADOS-2 calibrated severity scores from the highest module recorded, core descriptive IQ values, and Vineland-3 standard scores from the assessment at the highest recorded age. For the modified online SRS-2, we used the valid observed T score at the highest recorded assessment age across the school-age, dependent-adult other-report and adult self-report forms. At tied ages, we prioritized the dependent-adult other-report, school-age and adult self-report forms, in that order.

### Psychiatric and neurological GWAS datasets

Summary statistics for psychiatric disorders were obtained from published Psychiatric Genomics Consortium and related large-scale GWAS releases. The six-disorder panel for matched-module projection, the cross-disorder combined statistic and S-LDSC comprised schizophrenia (SCZ)^81^, bipolar disorder (BIP), major depressive disorder (MDD), attention-deficit/hyperactivity disorder (ADHD)^82^, anxiety disorder (ANX), and post-traumatic stress disorder (PTSD). The Genomic SEM analyses added autism spectrum disorder (ASD; iPSYCH) to these six disorders^5^. The competitive gene-set MAGMA analyses used ASD, the six psychiatric disorders, Alzheimer’s disease (AD) ^32^ and Parkinson’s disease (PD) ^83^. For BIP, we used the European-ancestry release excluding 23andMe participants^84^. For MDD, we used the European-ancestry subset of the trans-ancestry GWAS, excluding 23andMe participants^85^. For ANX, we used the publicly released European-ancestry summary statistics excluding the UTAH cohort^86^. For PTSD, we used the primary European-ancestry Freeze 3 analysis, which combines diagnostic and symptom-based phenotypes^87^. For AD, we used the publicly released European-ancestry summary statistics of the GWAS^32^, which exclude 23andMe data, with the SNP-specific effective sample sizes supplied by the authors and the native GRCh37 coordinates. The multi-ancestry release was analyzed as a sensitivity analysis with the same European LD reference (**Supplementary Note 5**).

### Family-based PRS transmission

ASD polygenic scores were computed from existing iPSYCH-v2 PRS-CS weights restricted to deduplicated SNP unions within gene bodies ±100 kb of each module’s genes. In 6,529 affected and 3,231 unaffected offspring of European ancestry from SPARK, SSC and MSSNG, transmission was the offspring-minus-midparent score standardized by the midparent-score SD^88^. Cohort estimates with family-clustered CR1 standard errors were combined by inverse-variance fixed-effect meta-analysis, separately for affected and unaffected offspring and for their contrast. Meta-analysis *P* values were obtained from two-sided z-tests and were Benjamini–Hochberg-adjusted across the 27 modules separately for affected offspring, unaffected offspring and their contrast.

Each module was compared with 4,000 matched gene sets retained from four sampling chains. Matching constrained gene length, SNP count, mean and summed LD scores, Grove-and iPSYCH-based genetic variance measures (sums of 2f(1 − f)β^2^, where f is allele frequency and β is the SNP effect), SNP proportions in four minor-allele-frequency bins (0.01–0.05, 0.05–0.10, 0.10–0.20 and 0.20–0.50), median brain expression, and the posterior mean of s_het_, together with chromosome and SNP-union size. Gene length, SNP count, the two variance measures and median brain expression were log1p-transformed. Each chain was initialized by mixed-integer linear programming and sampled using a tempered heat-bath algorithm. Candidate eligibility, matching tolerances and sampling intervals are described in **Supplementary Note 5**. For each retained set, the three cohort estimates were meta-analyzed as for the observed module, and the upper-tail empirical *P* value was (exceedances + 1)/(4,000 + 1). Empirical P values were Benjamini–Hochberg-adjusted across the 27 modules separately for affected and unaffected offspring. The Monte Carlo standard error of each empirical P was √(P(1 − P)/n_eff), where n_eff is the effective sample size of the exceedance indicator across the four chains. Gene constraint was represented by the GeneBayes posterior mean of s_het_, the selection coefficient against heterozygous loss-of-function variants^89^.

### Gene-level and module-level GWAS mapping

Per-SNP z-scores and effective sample sizes were reconstructed from raw data or VCF summary-statistic releases. Gene-level association statistics were computed with MAGMA using hg19 gene coordinates and the 1000 Genomes European reference panel^90,91^. The matched-module projection and the Genomic SEM component analyses used gene-body variant-to-gene mapping, whereas the nine-disorder gene-set analyses mapped SNPs to protein-coding genes with a ±10-kb window. MAGMA competitive gene-set tests were run separately for perturbation-module gene sets using the same gene-level association outputs, with MAGMA’s positive one-sided competitive test and MAGMA’s internal gene covariates. *P* values were corrected by the Benjamini–Hochberg procedure across 27 modules within each disorder, and across all 243 disorder–module tests as a separate correction. Genes whose bodies overlapped the MHC region (GRCh37 chr6:25,000,000–34,000,000; 284 genes in the gene-location file) were excluded at the gene-set stage; results including MHC genes are reported as a sensitivity analysis (**Supplementary Table 6c**).

For each of the nine disorders, we fitted the base MAGMA competitive model and three additional models conditioned on standardized log s_het_, standardized log coding-sequence length, or both covariates. The four models for each disorder used the same covariate-complete genes outside the MHC region, including 16,873 genes for ASD. *P* values were BH-adjusted across 27 modules separately for each disorder and model (**Supplementary Table 6c**).

For the gene-property-matched module projection, the background comprised modeled genes with MAGMA gene-level results, excluding genes whose start positions lay within the MHC interval defined above. Genes with BH-adjusted *P* < 0.05 within this background were prioritized. If fewer than 30 genes met this criterion, genes at or above the 99th percentile of MAGMA Z-statistics were used. For each disorder and module, the observed statistic was the number of prioritized genes assigned to that module. This count was compared with 3,000 matched null samples. Each prioritized gene was replaced by a random draw, with replacement, from the background genes in the same joint stratum of gene length, MAGMA SNP count, mean developmental expression and pLI. Each covariate was divided into deciles after breaking ties by input order, and missing pLI values were replaced by the median. The sampling pools included the prioritized genes. Matched-null z-scores were calculated as the standardized deviation of the observed module count from the matched-null distribution^90,92^.

S-LDSC analyses were restricted to the six-disorder panel. Module annotations were derived from SNPs within 100 kb of genes assigned to each module and tested in a baseline-LD-conditioned model^93–95^. Module-level two-sided S-LDSC enrichment *P* values and coefficient z-scores were reported. Concordance between matched-resampling and S-LDSC support was evaluated by comparing disorder-by-module matched-null z-scores with baseline-LD-conditioned S-LDSC coefficient z-scores.

### Cross-disorder and Genomic SEM analyses

Cross-disorder module enrichment was summarized across the six-disorder panel. For each module, disorder-level matched-null z-scores were combined using a correlation-adjusted Stouffer statistic, with the Pearson correlation matrix of gene-level MAGMA z-scores across 10,940 non-MHC model genes shared by the six disorders, yielding a one-sided *P* value for recurrent positive enrichment. These *P* values were Bonferroni-corrected across 27 modules (Supplementary Table 6g). Genomic SEM was used to separate shared psychiatric liability from disorder-specific heterogeneity^45^. Multivariable LDSC across seven disorders (SCZ, BIP, MDD, ADHD, ANX, PTSD and ASD) provided the genetic covariance matrix. Effective sample sizes were used, with the LDSC sample-prevalence parameter set to 0.5. Population prevalences were set to 1% for SCZ and BIP, 15% for MDD, 5% for ADHD, 10% for ANX, and 8% for PTSD. For ASD, we used the Danish population prevalence of 1.22% reported by Grove et al.^5^. The SCZ effective sample size was 146,348. LD scores were the standard 1000 Genomes European LD scores.

For the all-gene shared-liability and heterogeneity analysis, we fitted a common-factor Genomic SEM model to 1,018,159 HapMap3 SNPs. This model yielded two genome-wide association outputs: a common-factor association statistic that captures shared liability across the seven disorders, and a Q_SNP_ statistic that identifies SNPs with residual heterogeneity not explained by the common factor^45,96^. Both sets of SNP-level *P* values were converted to MAGMA gene-level association scores using a fixed effective sample size of 602,792 for Genomic SEM-derived inputs^90^.

Separately, we fitted a three-factor Genomic SEM model based on the factor structure described by Grotzinger et al. ^97^. SCZ and BIP loaded on a schizophrenia–bipolar factor, ASD and ADHD on a neurodevelopmental factor, and MDD, ANX and PTSD on an internalizing factor. Correlations between factors were estimated freely. In a sensitivity analysis, the ASD and ADHD loadings on the neurodevelopmental factor were constrained to be equal. Model-fit statistics are reported in **Supplementary Table 6d**. Factor-level association statistics for the three factors were converted to MAGMA gene-level profiles using the same Genomic SEM-to-MAGMA workflow.

All-gene Genomic SEM module projection was then performed by applying the MAGMA-to-module framework described above to the common-factor, Q_SNP_, and factor-level gene-level statistics. For each module and Genomic SEM component, the observed statistic was the mean MAGMA Z-statistic across module genes. The background comprised modeled genes with the corresponding MAGMA statistics, excluding genes whose bodies overlapped the MHC interval. We generated 10,000 matched null samples by drawing one gene with replacement for each set member from the same joint decile stratum of gene length, MAGMA SNP count, mean developmental expression, and pLI. Missing covariate values were replaced by their medians, and decile ties were broken by input order. Sampling pools included the evaluated set members^90,92^. Upper-tail empirical *P* values were (exceedances + 1)/(10,000 + 1).

We applied this matched-null comparison separately to M22 and to a glial-reference set comprising genes in M3, M11, M15, M19, M21, M28, M29 and M31. These modules were defined by glial-lineage perturbation and expression in oligodendrocyte precursor cells, oligodendrocytes, astrocytes or microglia. Spearman rank correlations across M22 genes compared the common-factor and Q_SNP_ profiles with the three factors and individual disorders. These correlations were descriptive (**Supplementary Table 6d**).

## Code availability

The code used for the analyses, figures and tables of this study will be made publicly available at https://github.com/joonan-lab/human-brain-geometry upon publication. The scDiffEq framework is available at https://github.com/scDiffEq/scDiffEq.

## Data availability

The data generated in this study are deposited on Zenodo (https://doi.org/10.5281/zenodo.24341520) and will be made publicly available upon publication. The record contains the developing human brain atlas with gene annotation updated to GENCODE v49 and HGNC, the trained scDiffEq model, the genome-wide knockout and overexpression perturbation map of 14,044 genes across 21 developmental lineages, the lineage responses at four perturbation strengths, and the simulation inputs needed to reproduce the results. The human developmental brain single-cell atlas (BTS atlas) dataset is available on Zenodo (https://doi.org/10.5281/zenodo.10939707)^11^. Genome-wide perturbation results, module assignments, and disease gene mapping are provided in Supplementary Tables.

## Acknowledgements

We acknowledge the contributions of all individuals and families participating in the Korean autism cohort, SSC, SPARK, and MSSNG, as well as the clinical sites and staff members involved. We also acknowledge access to genetic and phenotypic data from SSC and SPARK on SFARI Base, as well as from MSSNG on Autism Speaks. This study was supported by the National Research Foundation (NRF) of Korea (RS-2024-00439474 to H.J.Y. and J.-Y.A., RS-2025-00553304 to J.-Y.A.), Korea University Research Grant (#K2420391 to J.-Y.A.), the Korea-US Collaborative Research Fund (KUCRF), funded by the Ministry of Science and ICT and Ministry of Health & Welfare, Republic of Korea (RS-2024-00467876 to J.-Y.A.), the NRF grant funded by the Korea government (MSIT and MOE) (RS-2025-16652968 to J.-Y.A.), and the Basic Science Research Program through the NRF funded by the Ministry of Education (RS-2025-25438031 to I.G.K.). The Korea Bio Data Station (K-BDS) supported this work with computing resources, including technical support.

## Author contributions

I.G.K. and D.-M.S. contributed equally to this work. **Conceptualization**, I.G.K., D.-M.S. and J.-Y.A.; **Methodology**, I.G.K., D.-M.S., S.-W.K. and J.-Y.A.; **Software**, I.G.K., D.-M.S. and J.-Y.A.; **Validation**, I.G.K., D.-M.S., C.C., and J.-Y.A.; **Formal analysis**, I.G.K., D.-M.S., B.S.K. and J.-Y.A.; **Investigation**, I.G.K., D.-M.S., B.S.K. and J.-Y.A.; **Resources**, I.G.K., D.-M.S. and J.-Y.A.; **Sample collection**, J.H.S., J.W.L., G.B. and H.J.Y.; **Data curation**, I.G.K., D.-M.S., S.J.C., S.-W.K., H.L., J.H.S., J.W.L., G.B., H.J.Y. and J.-Y.A.; **Writing -original draft**, I.G.K., D.-M.S., B.S.K., S.J.C. and J.-Y.A.; **Writing -review and editing**, all authors; **Visualization**, I.G.K., D.-M.S., B.S.K., S.-W.K. and J.-Y.A.; **Supervision**, H.J.Y. and J.-Y.A.; **Project administration**, H.J.Y. and J.-Y.A.; **Funding acquisition**, H.J.Y. and J.-Y.A.

## Additional Information

**Supplementary Information** is available for this paper.

---

**Extended Data Fig. 1.**
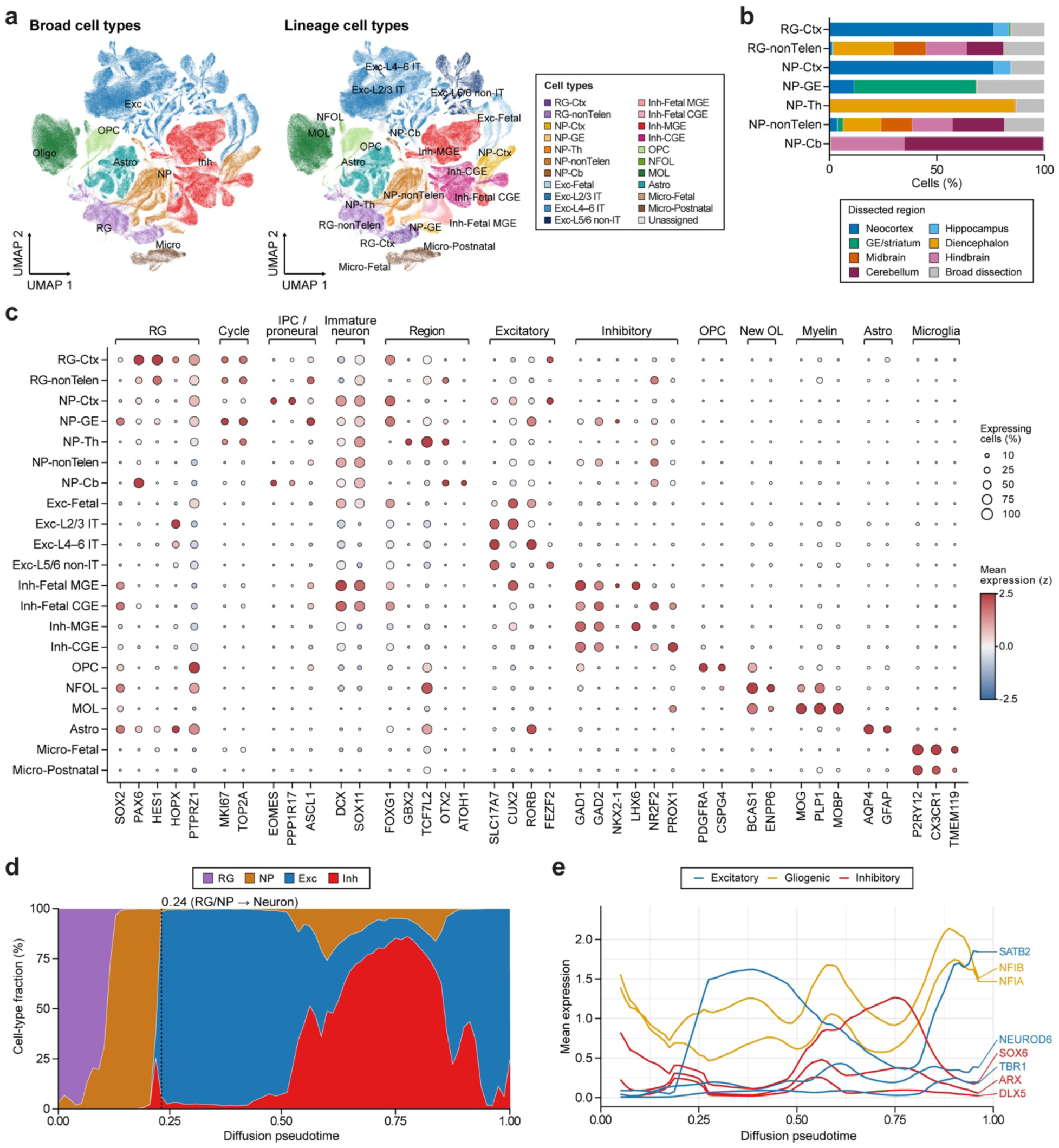
Refined cell-type annotation and developmental ordering. **a**, Comparison of broad and refined cell-type annotations. **b**, Regional composition of progenitor populations. **c**, Marker expression across annotated cell types (388,455 cells). Expression is standardized across cell types for each gene. **d**, Cell composition along rank-normalized diffusion pseudotime. The dashed line marks the first bin containing fewer than 50% radial glia and neuronal precursors combined. **e**, Smoothed marker expression along the same ordering. Diffusion pseudotime differs from the learned potential. Cell-type abbreviations follow Fig. 1.

**Extended Data Fig. 2.**
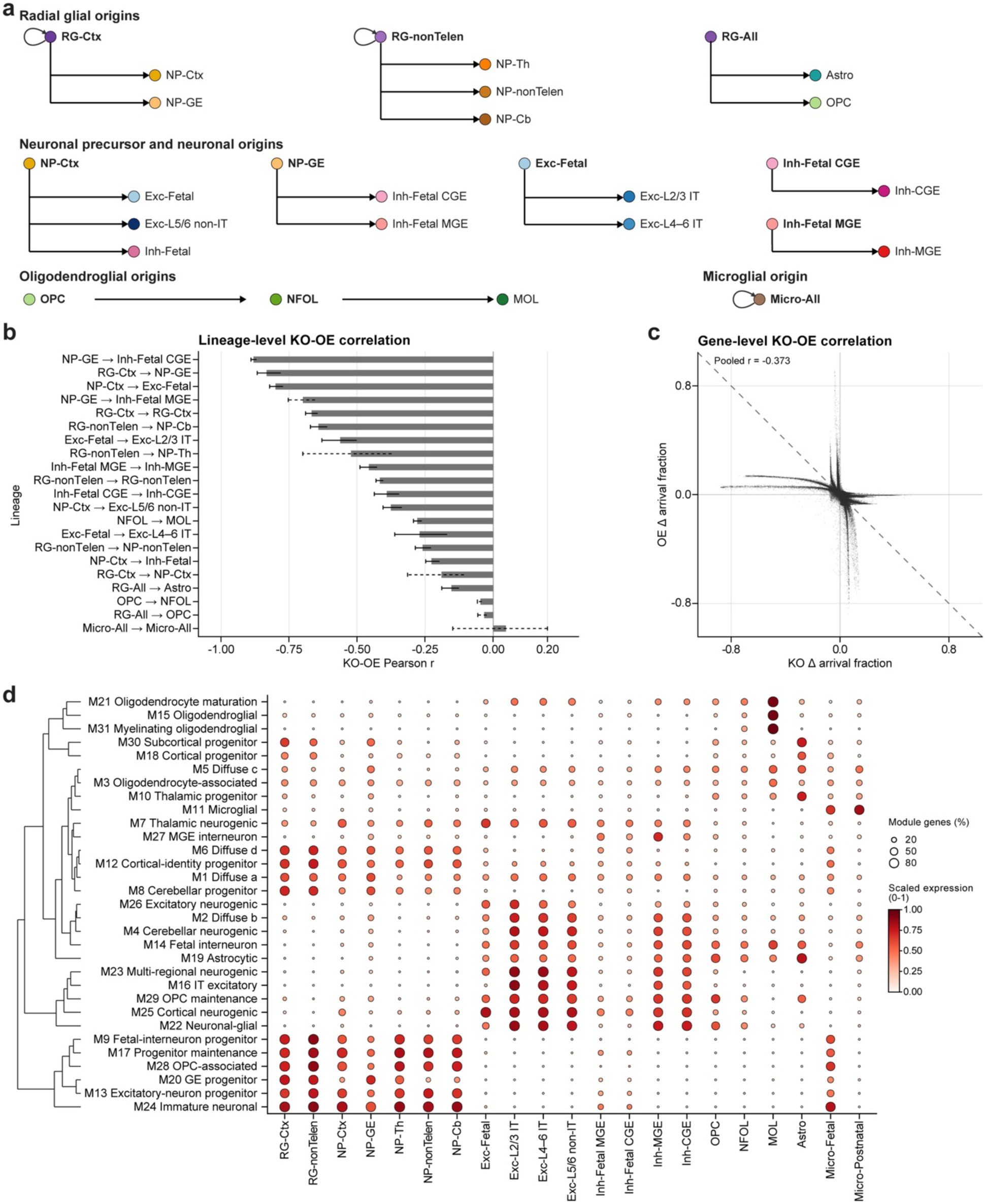
Defined lineages and KO–OE relationships. **a**, Starting and endpoint populations defining the simulated lineages, including identity retention. RG-All pools radial-glial types, Micro-All pools fetal and postnatal microglia, and Inh-Fetal pools fetal MGE/CGE types. **b**, KO–OE correlations across 14,044 genes for each lineage. Bars, 95% bootstrap CIs across simulation replicates. Dashed intervals flag unstable endpoints or WT floor/ceiling effects. **c**, Paired mean effects across the 17 non-boundary lineages, with each point representing a gene– lineage pair. Dashed line, equal and opposite effects. **d**, Module expression by cell type. Color, mean gene-scaled expression; dot area, proportion of module genes with scaled expression >0.5.

**Extended Data Fig. 3.**
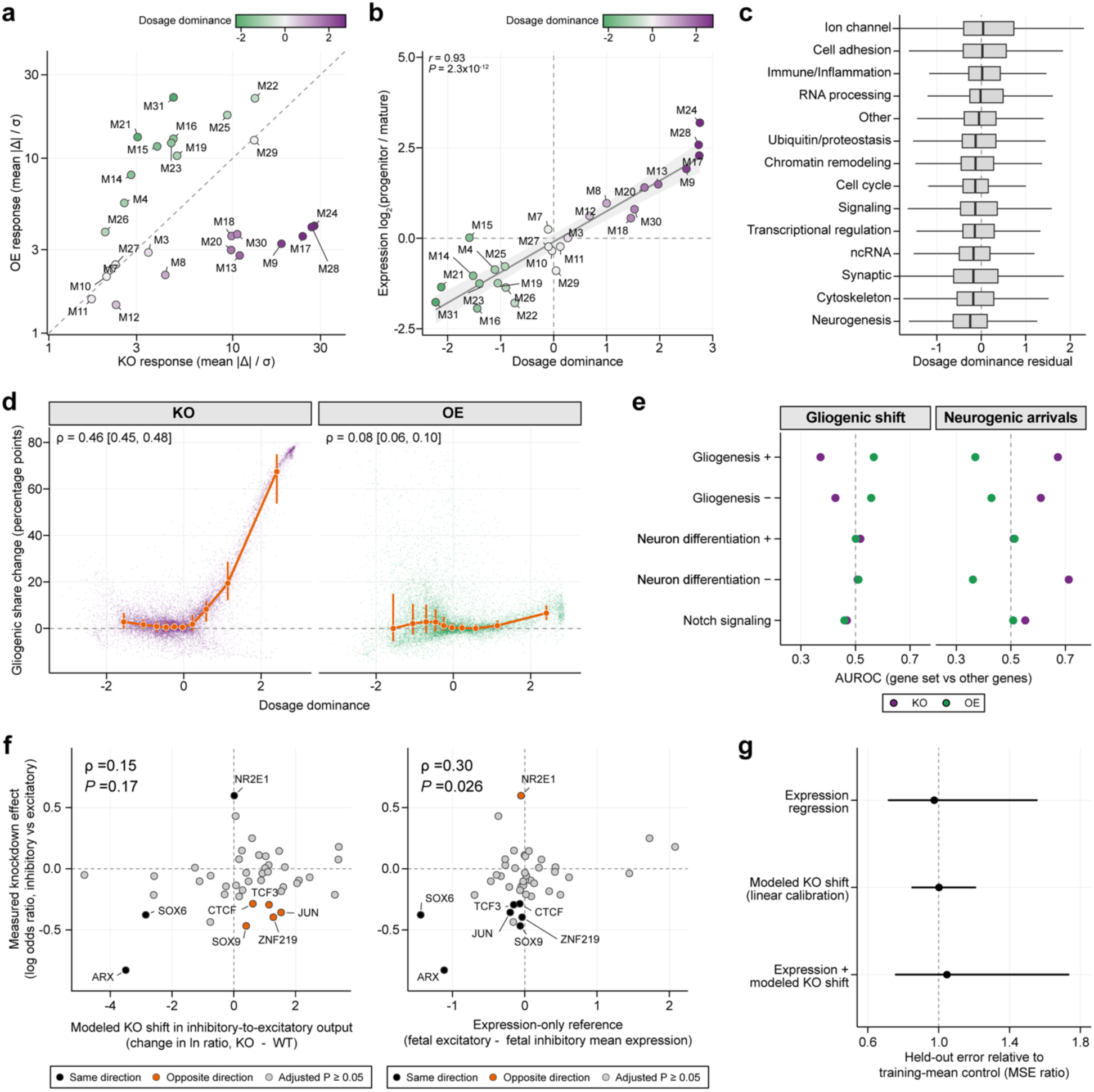
Developmental expression, dosage responses and comparison with experimental fate effects. **a**, Comparison of noise-scaled KO and OE response magnitudes across 27 modules. The dashed line indicates equal magnitudes. **b**, Relationship between the progenitor-to-mature expression ratio and dosage dominance across 27 modules. Line and shading indicate a linear fit and 95% confidence band. Two-sided Pearson test. **c**, Dosage dominance after adjustment for expression ratio and mean expression, grouped by functional class across 14,044 genes. Boxes show the median and interquartile range (IQR), with whiskers extending to the most extreme values within 1.5 × IQR. **d**, Relationship between dosage dominance and WT-relative changes in gliogenic share under KO and OE across 14,044 genes. Orange points and bars show decile medians and IQRs. Brackets give 95% bootstrap confidence intervals for Spearman correlations. **e**, GO-set comparisons after excluding M30/M10 genes (left) and for WT-relative changes in neuronal-precursor endpoint counts using all genes (right). AUROC > 0.5 indicates higher responses in the set than in other genes. **f**, Measured knockdown effects for 43 transcription factors compared with modeled KO shifts (left) and an expression-only reference (right). Directional matches and mismatches are highlighted for factors with experimental Benjamini–Hochberg-adjusted *P* < 0.05. Spearman correlations use one-sided permutation tests. **g**, Prediction error of regressions using expression, modeled KO shifts or both, evaluated by leave-one-factor-out cross-validation across 43 factors. Points and bars show mean squared error relative to a training-mean control and 95% confidence intervals from 2,000 factor-level bootstrap resamples.

**Extended Data Fig. 4.**
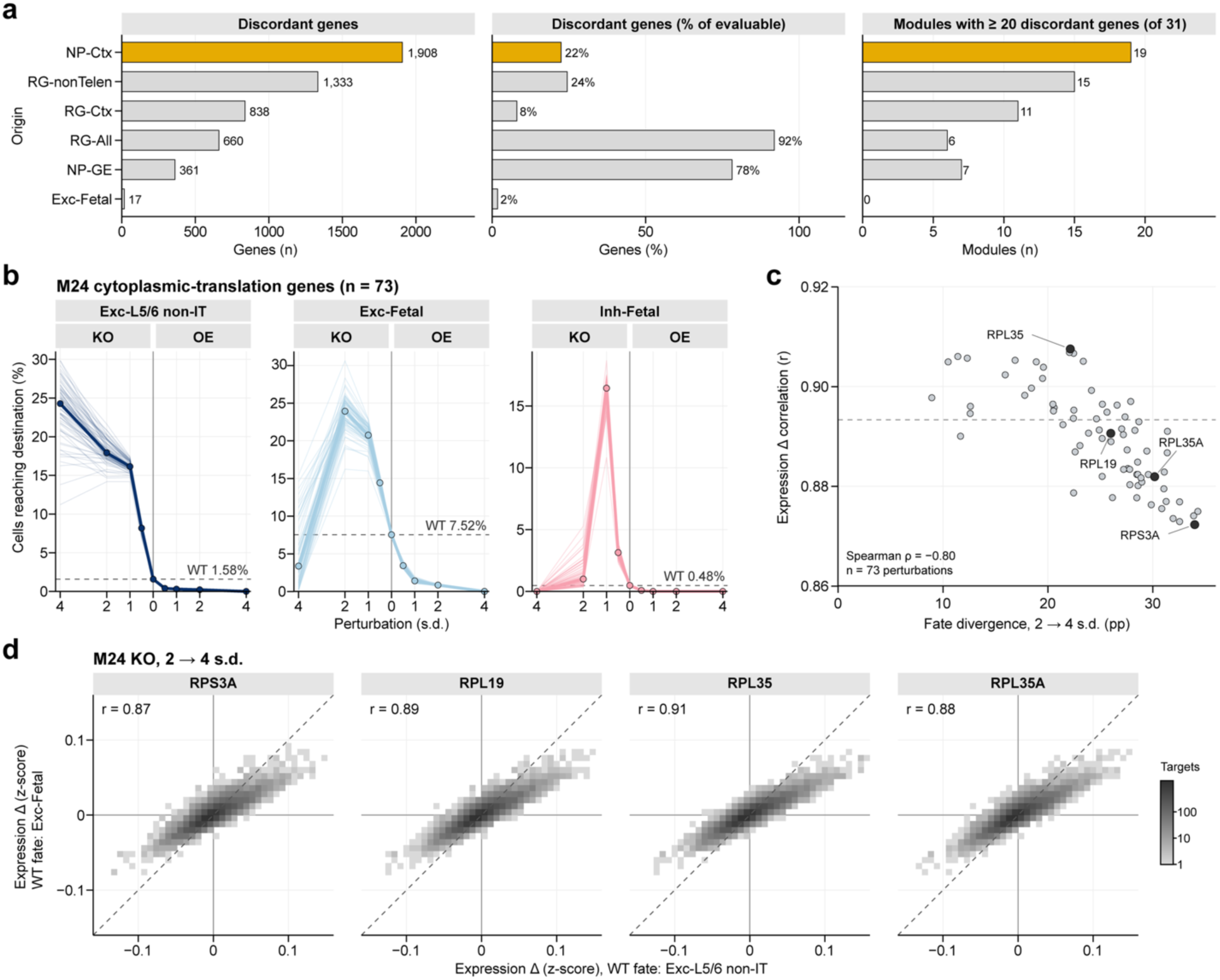
Destination discordance and downstream responses of M24 genes. **a**, Numbers and fractions of discordant genes, and numbers of modules containing at least 20 such genes, by origin. Discordance denotes different response classes in at least two destinations. **b**, Percentage of simulated cells reaching each endpoint after perturbation of M24 genes. Thin lines, individual gene means; thick lines, medians; dashed lines, WT values. **c**, Additional fate divergence from 2 to 4 s.d. KO versus expression-response correlation for each M24 perturbation. Fate divergence is the difference between additional shifts toward layer 5/6 non-IT and fetal excitatory states, in pp. Dashed line, correlation of the perturbation-averaged responses. **d**, Target-gene changes for perturbations with the lowest, median and highest correlations in **c**, and *RPL35A*. Expression changes are input-subtracted in the two WT-fate groups. Dashed lines, equal responses.

**Extended Data Fig. 5.**
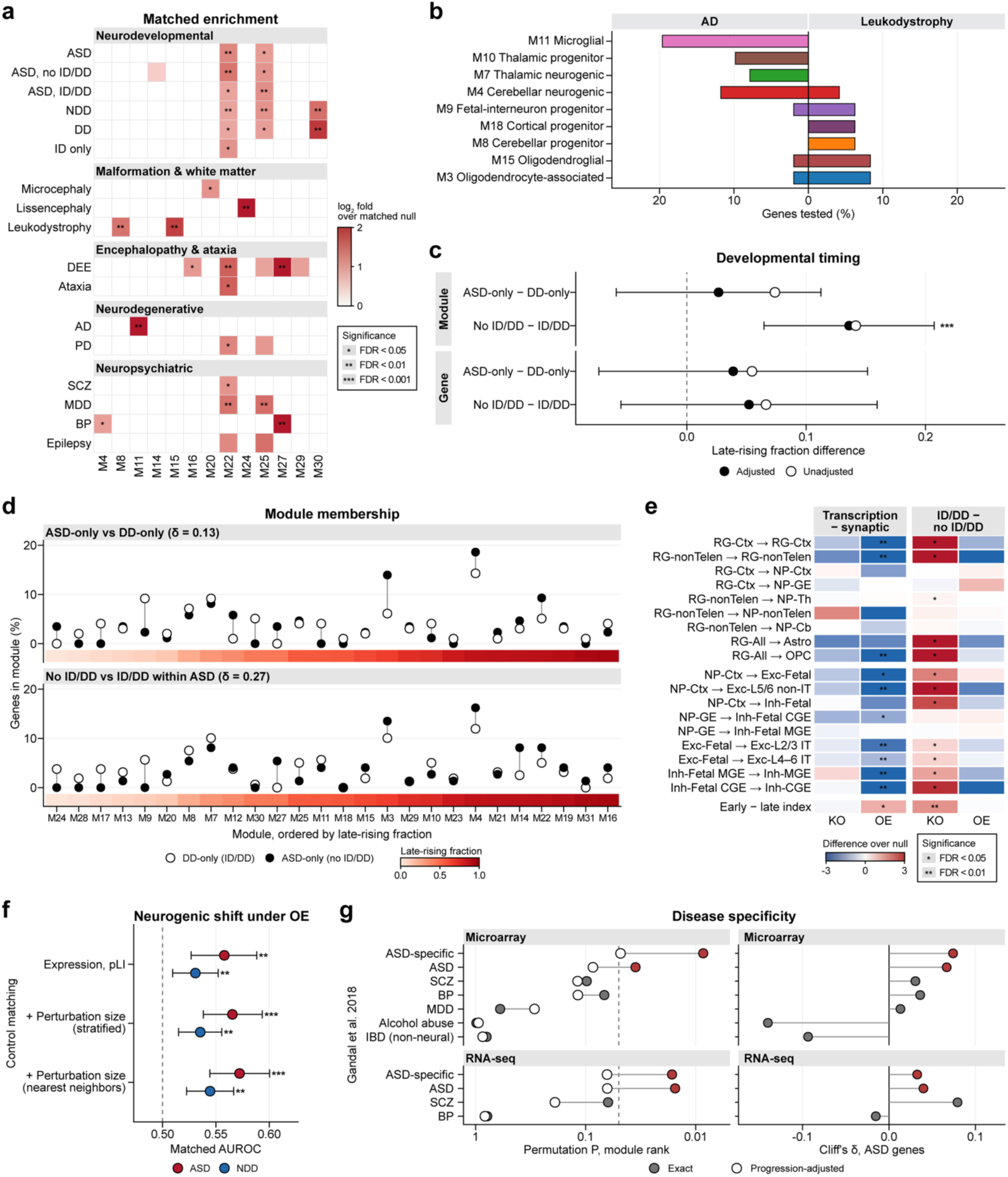
Disease-program comparisons and developmental and cortical contexts of ASD risk. **a**, Covariate-matched enrichment of disease–module associations. Gray, untested. **b**, Module distributions of nonoverlapping AD and leukodystrophy genes (n = 51 and 48). Modules containing ≥3 genes in either group are displayed. **c**, Late-rising fraction differences between exclusive disease or clinical groups, at module and gene levels. Bars, 95% CIs for expression- and constraint-adjusted differences. Group sizes for module/gene timing are 86/98 and 106/116 for ASD-only/DD-only, and 74/159 and 90/190 for no-ID/DD/ID/DD. Asterisks indicate nominal *P values from the covariate-adjusted models*. **d**, Module membership of groups in **c**, ordered by module timing. δ, unadjusted Cliff’s delta. **e**, Functional-class and clinical-stratum contrasts in response magnitude beyond matched expectations. Final row, early-minus-late index. **f**, Neurogenic OE drive against matched backgrounds. Points, mean AUROC; intervals, 2.5th– 97.5th percentiles across matched backgrounds. **g**, Concordance between modeled KO expression signatures and postmortem cortical changes. Left, ranking of the three ASD-associated modules before and after developmental-progression adjustment; dashed line, *P* = 0.05. Right, expression- and constraint-stratified Cliff’s delta for all 290 ASD genes. ASD-specific contrasts remove shared psychiatric-disease changes. BP, bipolar disorder; IBD, inflammatory bowel disease.

**Extended Data Fig. 6.**
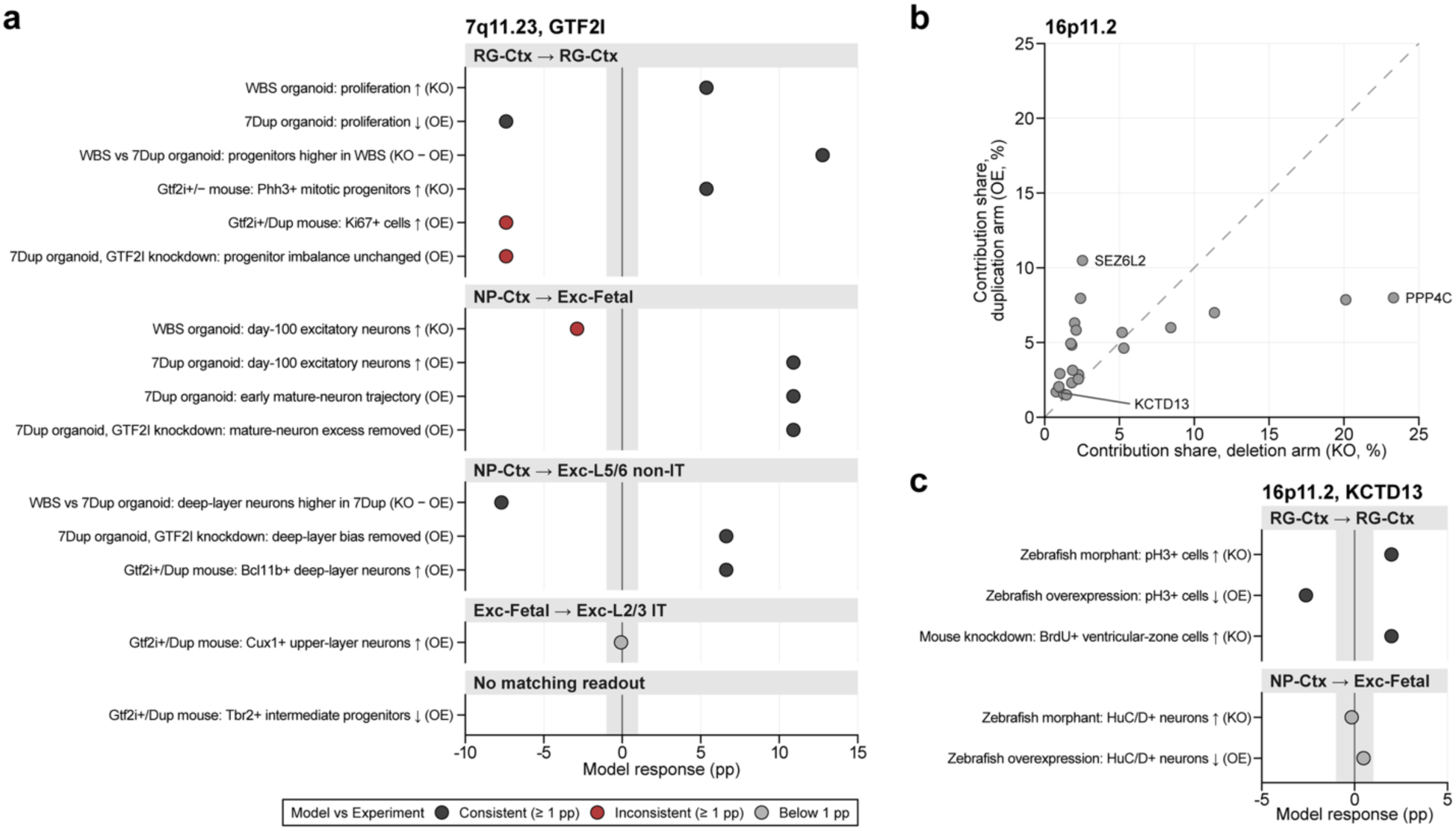
CNV-related dosage experiments compared with modeled responses. **a**, Fifteen reported 7q11.23 findings and corresponding model responses. The KO − OE contrast represents deletion-versus-duplication comparisons. Tbr2-positive intermediate progenitors lack a matching model readout. **b**, Within-arm response-score shares for 16p11.2 genes; dashed line, equal shares. **c**, Comparison of reported *KCTD13* phenotypes with corresponding model responses. In a and **c**, points are WT-relative changes in endpoint percentage (pp), and gray bands mark absolute changes below 1 pp. Radial-glial retention provides an indirect comparison with proliferation; multiple findings can share one model response.

**Extended Data Fig. 7.**
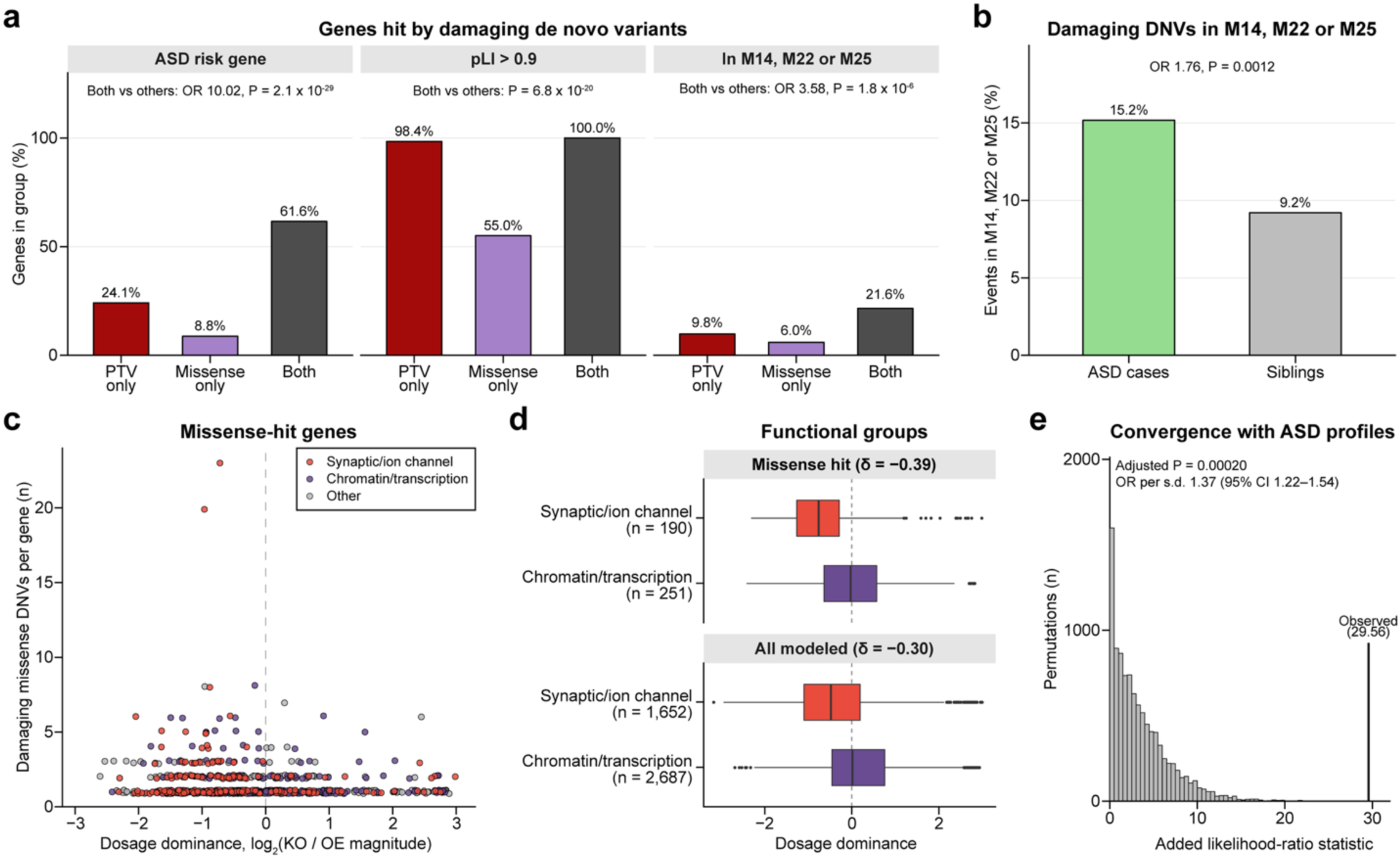
De novo variant-associated genes and perturbation profiles. **a**, Properties of genes hit by damaging PTVs only (n = 316), missense variants only (n = 705) or both (n = 125). ASD-risk fractions use the annotated subsets (311, 638 and 125 genes). **b**, Pooled de novo events in M14, M22 or M25 among cases (323/2,129) and siblings (38/413). The unit is the variant event. **c**, Relationship between dosage dominance and damaging missense variant counts per gene. **d**, Differences in dosage dominance between functional classes among missense-hit genes and among all modeled genes, including the missense-hit genes. Boxes, median and IQR; whiskers, 1.5 x IQR. δ, gene-level Cliff’s delta. **e**, Convergence of 376 recurrently hit genes with 126 nonoverlapping ASD reference genes. Histogram, permutation distribution; line, observed statistic. The OR per s.d. is adjusted for constraint and co-expression.

## References

1 Fu, J. M. et al. Rare coding variation provides insight into the genetic architecture and phenotypic context of autism. Nat Genet 54, 1320–1331 (2022). 10.1038/s41588-022-01104-0

2 Satterstrom, F. et al. Large-Scale Exome Sequencing Study Implicates Both Developmental and Functional Changes in the Neurobiology of Autism. Cell 180, 568–584.e23 (2020). 10.1016/j.cell.2019.12.036

3 Consortium, B. et al. Analysis of shared heritability in common disorders of the brain. Science 360 (2018). 10.1126/science.aap8757

4 Cross-Disorder Group of the Psychiatric Genomics Consortium. Genomic Relationships, Novel Loci, and Pleiotropic Mechanisms across Eight Psychiatric Disorders. Cell 179, 1469–1482 e11 (2019). 10.1016/j.cell.2019.11.020

5 Grove, J. et al. Identification of common genetic risk variants for autism spectrum disorder. Nat Genet 51, 431–444 (2019). 10.1038/s41588-019-0344-8

6 Gaugler, T. et al. Most genetic risk for autism resides with common variation. Nat Genet 46, 881–5 (2014). 10.1038/ng.3039

7 Niemi, M. E. K. et al. Common genetic variants contribute to risk of rare severe neurodevelopmental disorders. Nature 562, 268–271 (2018). 10.1038/s41586-018-0566-4

8 Antaki, D. et al. A phenotypic spectrum of autism is attributable to the combined effects of rare variants, polygenic risk and sex. Nat Genet 54, 1284–1292 (2022). 10.1038/s41588-022-01064-5

9 Weiner, D. J. et al. Statistical and functional convergence of common and rare genetic influences on autism at chromosome 16p. Nat Genet 54, 1630–1639 (2022). 10.1038/s41588-022-01203-y

10 Abe, H. et al. Convergent coexpression reveals shared biological mechanisms underlying common and rare variant risk in six neuropsychiatric disorders. Mol Psychiatry 31, 4787–4798 (2026). 10.1038/s41380-026-03571-x

11 Kim, S. et al. An integrative single-cell atlas for exploring the cellular and temporal specificity of genes related to neurological disorders during human brain development. Exp Mol Med 56, 2271–2282 (2024). 10.1038/s12276-024-01328-6

12 Jin, X. et al. In vivo Perturb-Seq reveals neuronal and glial abnormalities associated with autism risk genes. Science 370 (2020). 10.1126/science.aaz6063

13 Fernandez Garcia, M., et al. Transcriptomic and phenotypic convergence of neurodevelopmental disorder risk genes in vitro and in vivo. Nat Neurosci 29, 1079–1094 (2026). 10.1038/s41593-026-02247-7

14 Gordon, A. et al. Developmental convergence and divergence in human stem cell models of autism. Nature 651, 707–719 (2026). 10.1038/s41586-025-10047-5

15 Parikshak, N. N. et al. Genome-wide changes in lncRNA, splicing, and regional gene expression patterns in autism. Nature 540, 423–427 (2016). 10.1038/nature20612

16 Gandal, M. J. et al. Transcriptome-wide isoform-level dysregulation in ASD, schizophrenia, and bipolar disorder. Science 362 (2018). 10.1126/science.aat8127

17 Velmeshev, D. et al. Single-cell genomics identifies cell type-specific molecular changes in autism. Science 364, 685–689 (2019). 10.1126/science.aav8130

18 Wamsley, B. et al. Molecular cascades and cell type-specific signatures in ASD revealed by single-cell genomics. Science 384, eadh2602 (2024). 10.1126/science.adh2602

19 Kawamura, A. et al. Duplication of the autism-related gene Chd8 leads to behavioral hyperactivity and neurodevelopmental defects in mice. Nat Commun 16, 4641 (2025). 10.1038/s41467-025-59853-5

20 Collins, R. L. et al. A cross-disorder dosage sensitivity map of the human genome. Cell 185, 3041–3055 e25 (2022). 10.1016/j.cell.2022.06.036

21 Shi, T. et al. Genome-scale functional mapping of the mammalian whole brain with in vivo Perturb-seq. bioRxiv (2026). 10.64898/2026.03.16.711480

22 Theodoris, C. V. et al. Transfer learning enables predictions in network biology. Nature 618, 616–624 (2023). 10.1038/s41586-023-06139-9

23 Kamimoto, K. et al. Dissecting cell identity via network inference and in silico gene perturbation. Nature 614, 742–751 (2023). 10.1038/s41586-022-05688-9

24 Wang, H., Li, Q. & Zhu, Y. An in silico transcription factor perturbation simulator uncovers diverse genetic architectures of brain disorders. Genome Medicine (2026). 10.1186/s13073-026-01752-5

25 Koh, I. G. et al. AI-driven framework modeling perturbation in brain organoids reveals candidate genes for autism. bioRxiv, 2026.08.22.746387 (2026). 10.64898/2026.08.22.746387

26 Vinyard, M. E. et al. Learning cell dynamics with neural differential equations. Nat Mach Intell 7, 1969–1984 (2025). 10.1038/s42256-025-01150-3

27 Tchieu, J. et al. NFIA is a gliogenic switch enabling rapid derivation of functional human astrocytes from pluripotent stem cells. Nat Biotechnol 37, 267–275 (2019). 10.1038/s41587-019-0035-0

28 Leung, R. F. et al. The DLX/Notch axis is necessary for spatiotemporal regulation of neural cell fate. Nat Commun 17 (2026). 10.1038/s41467-026-76838-0

29 Ding, J. W. et al. Dissecting gene regulatory networks governing human cortical cell fate. Nature 651, 732–742 (2026). 10.1038/s41586-025-09997-7

30 Satterstrom, F. K. et al. Rare variation illuminates the distinct and pleiotropic genetic architecture of autism across neuropsychiatric traits. medRxiv, 2026.08.24.26360398 (2026). 10.64898/2026.08.24.26360398

31 Zhang, Q. et al. Temporal regulation of progenitor lineage progression and output by NFIs underlying human neocortical malformation. Neuron (2026). 10.1016/j.neuron.2026.08.021

32 Uffelmann, E. et al. Alzheimer’s disease genetics converges on microglial and neuronal dysregulation. Nature (2026). 10.1038/s41586-026-11206-y

33 Nott, A. et al. Brain cell type-specific enhancer-promoter interactome maps and disease-risk association. Science 366, 1134–1139 (2019). 10.1126/science.aay0793

34 Bellenguez, C. et al. New insights into the genetic etiology of Alzheimer’s disease and related dementias. Nat Genet 54, 412–436 (2022). 10.1038/s41588-022-01024-z

35 Jayaraman, D., Bae, B. I. & Walsh, C. A. The Genetics of Primary Microcephaly. Annu Rev Genomics Hum Genet 19, 177–200 (2018). 10.1146/annurev-genom-083117-021441

36 Thornton, G. K. & Woods, C. G. Primary microcephaly: do all roads lead to Rome? Trends Genet 25, 501–10 (2009). 10.1016/j.tig.2009.09.011

37 Gandal, M. J. et al. Shared molecular neuropathology across major psychiatric disorders parallels polygenic overlap. Science 359, 693–697 (2018). 10.1126/science.aad6469

38 Gandal, M. J. et al. Broad transcriptomic dysregulation occurs across the cerebral cortex in ASD. Nature 611, 532–539 (2022). 10.1038/s41586-022-05377-7

39 Yin, J. et al. Otud7a Knockout Mice Recapitulate Many Neurological Features of 15q13.3 Microdeletion Syndrome. Am J Hum Genet 102, 296–308 (2018). 10.1016/j.ajhg.2018.01.005

40 Uddin, M. et al. OTUD7A Regulates Neurodevelopmental Phenotypes in the 15q13.3 Microdeletion Syndrome. Am J Hum Genet 102, 278–295 (2018). 10.1016/j.ajhg.2018.01.006

41 Lopez-Tobon, A. et al. GTF2I dosage regulates neuronal differentiation and social behavior in 7q11.23 neurodevelopmental disorders. Sci Adv 9, eadh2726 (2023). 10.1126/sciadv.adh2726

42 Golzio, C. et al. KCTD13 is a major driver of mirrored neuroanatomical phenotypes of the 16p11.2 copy number variant. Nature 485, 363–7 (2012). 10.1038/nature11091

43 Consortium, G. T. The GTEx Consortium atlas of genetic regulatory effects across human tissues. Science 369, 1318–1330 (2020). 10.1126/science.aaz1776

44 Mohammadi, P., Castel, S. E., Brown, A. A. & Lappalainen, T. Quantifying the regulatory effect size of cis-acting genetic variation using allelic fold change. Genome Res 27, 1872–1884 (2017). 10.1101/gr.216747.116

45 Grotzinger, A. et al. Genomic structural equation modelling provides insights into the multivariate genetic architecture of complex traits. Nat Hum Behav 3, 513–525 (2019). 10.1038/s41562-019-0566-x

46 Fazel Darbandi, S., et al. Five autism-associated transcriptional regulators target shared loci proximal to brain-expressed genes. Cell Rep 43, 114329 (2024). 10.1016/j.celrep.2024.114329

47 Andersen, R. E. et al. Transcriptomic convergence and the female protective effect in autism. Nature (2026). 10.1038/s41586-026-11236-6

48 Bright, A. R. et al. Temporal control of progenitor competence shapes maturation in GABAergic neuron development in mice. Nat Neurosci 28, 1663–1675 (2025). 10.1038/s41593-025-01999-y

49 Hurley, S. et al. Distinct, dosage-sensitive requirements for the autism-associated factor CHD8 during cortical development. Mol Autism 12, 16 (2021). 10.1186/s13229-020-00409-3

50 Shin, D. et al. Thalamocortical organoids enable in vitro modeling of 22q11.2 microdeletion associated with neuropsychiatric disorders. Cell Stem Cell 31, 421–432 e8 (2024). 10.1016/j.stem.2024.01.010

51 Wolf, F. A., Angerer, P. & Theis, F. J. SCANPY: large-scale single-cell gene expression data analysis. Genome Biol 19, 15 (2018). 10.1186/s13059-017-1382-0

52 Haghverdi, L., Buttner, M., Wolf, F. A., Buettner, F. & Theis, F. J. Diffusion pseudotime robustly reconstructs lineage branching. Nat Methods 13, 845–8 (2016). 10.1038/nmeth.3971

53 Traag, V., Waltman, L. & van Eck, N. From Louvain to Leiden: guaranteeing well-connected communities. Sci Rep 9, 5233 (2019). 10.1038/s41598-019-41695-z

54 Chen, S. et al. A genomic mutational constraint map using variation in 76,156 human genomes. Nature 625, 92–100 (2024). 10.1038/s41586-023-06045-0

55 Herring, C. A. et al. Human prefrontal cortex gene regulatory dynamics from gestation to adulthood at single-cell resolution. Cell 185, 4428–4447 e28 (2022). 10.1016/j.cell.2022.09.039

56 Trevino, A. E. et al. Chromatin accessibility dynamics in a model of human forebrain development. Science 367 (2020). 10.1126/science.aay1645

57 Velmeshev, D. et al. Single-cell analysis of prenatal and postnatal human cortical development. Science 382, eadf0834 (2023). 10.1126/science.adf0834

58 Ziller, M. J. et al. Dissecting neural differentiation regulatory networks through epigenetic footprinting. Nature 518, 355–359 (2015). 10.1038/nature13990

59 Schaaf, C. P. et al. A framework for an evidence-based gene list relevant to autism spectrum disorder. Nat Rev Genet 21, 367–376 (2020). 10.1038/s41576-020-0231-2

60 Singh, T. et al. Rare coding variants in ten genes confer substantial risk for schizophrenia. Nature 604, 509–516 (2022). 10.1038/s41586-022-04556-w

61 Meng, X. et al. Multi-ancestry genome-wide association study of major depression aids locus discovery, fine mapping, gene prioritization and causal inference. Nat Genet 56, 222–233 (2024). 10.1038/s41588-023-01596-4

62 Palmer, D. S. et al. Exome sequencing in bipolar disorder identifies AKAP11 as a risk gene shared with schizophrenia. Nat Genet 54, 541–547 (2022). 10.1038/s41588-022-01034-x

63 Epi, C. Exome sequencing of 20,979 individuals with epilepsy reveals shared and distinct ultra-rare genetic risk across disorder subtypes. Nat Neurosci 27, 1864–1879 (2024). 10.1038/s41593-024-01747-8

64 Landrum, M. J. et al. ClinVar: improving access to variant interpretations and supporting evidence. Nucleic Acids Res 46, D1062–D1067 (2018). 10.1093/nar/gkx1153

65 Vasilevsky, N. A. et al. Mondo: integrating disease terminology across communities. Genetics 232 (2026). 10.1093/genetics/iyaf215

66 Kim, S. W. et al. Whole genome sequencing analysis identifies sex differences of familial pattern contributing to phenotypic diversity in autism. Genome Med 16, 114 (2024). 10.1186/s13073-024-01385-6

67 SPARK Consortium. SPARK: A US Cohort of 50,000 Families to Accelerate Autism Research. Neuron 97, 488–493 (2018). 10.1016/j.neuron.2018.01.015

68 Fischbach, G. D. & Lord, C. The Simons Simplex Collection: a resource for identification of autism genetic risk factors. Neuron 68, 192–5 (2010). 10.1016/j.neuron.2010.10.006

69 Trost, B. et al. Genomic architecture of autism from comprehensive whole-genome sequence annotation. Cell 185, 4409–4427 e18 (2022). 10.1016/j.cell.2022.10.009

70 Yoo, H., et al. Korean autism diagnostic observation schedule-2 (K-ADOS-2). Seoul: Hakjisa (2018).

71 Gotham, K., Pickles, A. & Lord, C. Standardizing ADOS scores for a measure of severity in autism spectrum disorders. J Autism Dev Disord 39, 693–705 (2009). 10.1007/s10803-008-0674-3

72 Kim, J.-H. et al. A Validation Study of the Korean Version of Social Communication Questionnaire. J Korean Acad Child Adolesc Psychiatry 26, 197–208 (2015). 10.5765/jkacap.2015.26.3.197

73 Constantino, J. N. et al. Validation of a brief quantitative measure of autistic traits: comparison of the social responsiveness scale with the autism diagnostic interview-revised. J Autism Dev Disord 33, 427–33 (2003). 10.1023/a:1025014929212

74 Cheon, K. A. et al. The social responsiveness scale in relation to DSM IV and DSM5 ASD in Korean children. Autism Res 9, 970–80 (2016). 10.1002/aur.1671

75 Rutter, M., LeCouteur, A. & Lord, C. Autism diagnostic interview-revised. American Journal on Mental Retardation (2003).

76 Lam, K. S. & Aman, M. G. The Repetitive Behavior Scale-Revised: independent validation in individuals with autism spectrum disorders. J Autism Dev Disord 37, 855–66 (2007). 10.1007/s10803-006-0213-z

77 Wechsler, D. Wechsler Intelligence Scale for Children–Fourth Edition (WISC-IV). (Psychological Corporation, 2003).

78 Wechsler, D. WPPSI-III: Administration and Scoring Manual. (Psychological Corporation, 2002).

79 Sparrow, S. S., Cicchetti, D. V. & Balla, D. A. Vineland Adaptive Behavior Scales–Second Edition (VABS-II). (Pearson Assessments, 2005).

80 Wilson, B. N. et al. Psychometric properties of the revised Developmental Coordination Disorder Questionnaire. Phys Occup Ther Pediatr 29, 182–202 (2009). 10.1080/01942630902784761

81 Trubetskoy, V. et al. Mapping genomic loci implicates genes and synaptic biology in schizophrenia. Nature 604, 502–508 (2022). 10.1038/s41586-022-04434-5

82 Demontis, D. et al. Genome-wide analyses of ADHD identify 27 risk loci, refine the genetic architecture and implicate several cognitive domains. Nat Genet 55, 198–208 (2023). 10.1038/s41588-022-01285-8

83 Nalls, M. A. et al. Identification of novel risk loci, causal insights, and heritable risk for Parkinson’s disease: a meta-analysis of genome-wide association studies. Lancet Neurol 18, 1091–1102 (2019). 10.1016/S1474-4422(19)30320-5

84 O’Connell, K. S. et al. Genomics yields biological and phenotypic insights into bipolar disorder. Nature 639, 968–975 (2025). 10.1038/s41586-024-08468-9

85 Major Depressive Disorder Working Group of the Psychiatric Genomics Consortium. Trans-ancestry genome-wide study of depression identifies 697 associations implicating cell types and pharmacotherapies. Cell 188, 640–652 e9 (2025). 10.1016/j.cell.2024.12.002

86 Strom, N. I. et al. Genome-wide association study of major anxiety disorders in 122,341 European-ancestry cases identifies 58 loci and highlights GABAergic signaling. Nat Genet 58, 275–288 (2026). 10.1038/s41588-025-02485-8

87 Nievergelt, C. M. et al. Genome-wide association analyses identify 95 risk loci and provide insights into the neurobiology of post-traumatic stress disorder. Nat Genet 56, 792–808 (2024). 10.1038/s41588-024-01707-9

88 Weiner, D. J. et al. Polygenic transmission disequilibrium confirms that common and rare variation act additively to create risk for autism spectrum disorders. Nat Genet 49, 978–985 (2017). 10.1038/ng.3863

89 Zeng, T., Spence, J. P., Mostafavi, H. & Pritchard, J. K. Bayesian estimation of gene constraint from an evolutionary model with gene features. Nat Genet 56, 1632–1643 (2024). 10.1038/s41588-024-01820-9

90 de Leeuw, C., Mooij, J., Heskes, T. & Posthuma, D. MAGMA: generalized gene-set analysis of GWAS data. PLoS Comput Biol 11, e1004219 (2015). 10.1371/journal.pcbi.1004219

91 Consortium, G. P. et al. A global reference for human genetic variation. Nature 526, 68–74 (2015). 10.1038/nature15393

92 Lek, M. et al. Analysis of protein-coding genetic variation in 60,706 humans. Nature 536, 285–91 (2016). 10.1038/nature19057

93 Bulik-Sullivan, B. K. et al. LD Score regression distinguishes confounding from polygenicity in genome-wide association studies. Nat Genet 47, 291–5 (2015). 10.1038/ng.3211

94 Finucane, H. et al. Partitioning heritability by functional annotation using genome-wide association summary statistics. Nat Genet 47, 1228–35 (2015). 10.1038/ng.3404

95 Gazal, S. et al. Linkage disequilibrium-dependent architecture of human complex traits shows action of negative selection. Nat Genet 49, 1421–1427 (2017). 10.1038/ng.3954

96 International HapMap, C., et al. Integrating common and rare genetic variation in diverse human populations. Nature 467, 52–8 (2010). 10.1038/nature09298

97 Grotzinger, A. D. et al. Genetic architecture of 11 major psychiatric disorders at biobehavioral, functional genomic and molecular genetic levels of analysis. Nat Genet 54, 548–559 (2022). 10.1038/s41588-022-01057-4

