## Supplementary Information for "Developmental timing of gene expression predicts dosage dominance, which tracks neuronal–glial fate in the developing human brain"

**Extended Data Figures**

**Extended Data Fig. 1.** Refined cell-type annotation and developmental ordering.

**Extended Data Fig. 2.** Defined lineages and KO–OE relationships.

**Extended Data Fig. 3.** Developmental expression, dosage responses and comparison with experimental fate effects.

**Extended Data Fig. 4.** Destination discordance and downstream responses of M24 genes.

**Extended Data Fig. 5.** Disease-program comparisons and developmental and cortical contexts of ASD risk.

**Extended Data Fig. 6.** CNV-related dosage experiments compared with modeled responses.

**Extended Data Fig. 7.** De novo variant-associated genes and perturbation profiles.

**Supplementary Notes**

**Supplementary Note 1**. Supporting evidence for cell-type annotation and lineage specificity

**Supplementary Note 2.** Robustness of perturbation map and module clustering

**Supplementary Note 3.** Biological annotation of perturbation-response modules

**Supplementary Note 4**. Simulation variability and sensitivity analyses

**Supplementary Note 5**. Supporting analyses of rare and common risk across perturbation modules

**Supplementary Figures**

**Supplementary Fig. 1.** Module-clustering stability and sensitivity to simulation variation.

**Supplementary Fig. 2.** Developmental expression trajectories and their composition across 27 lineage-associated modules.

**Supplementary Tables**

**Supplementary Table 1.** Genome-wide in silico perturbation profiles and fate-regulatory module annotations.

**Supplementary Table 2**. Developmental timing, expression position and neurogenic-gliogenic allocation in relation to dosage dominance

**Supplementary Table 3.** Strength-dependent fate responses, destination discordance and the *NFIA* and M24 examples

**Supplementary Table 4.** Disease gene sets, developmental response programs and ASD analyses.

**Supplementary Table 5.** CNV intervals as dosage proxies for the knockout and overexpression perturbation map

**Supplementary Table 6.** Rare and common genetic risk across perturbation modules

**Supplementary References
Extended Data Figures**

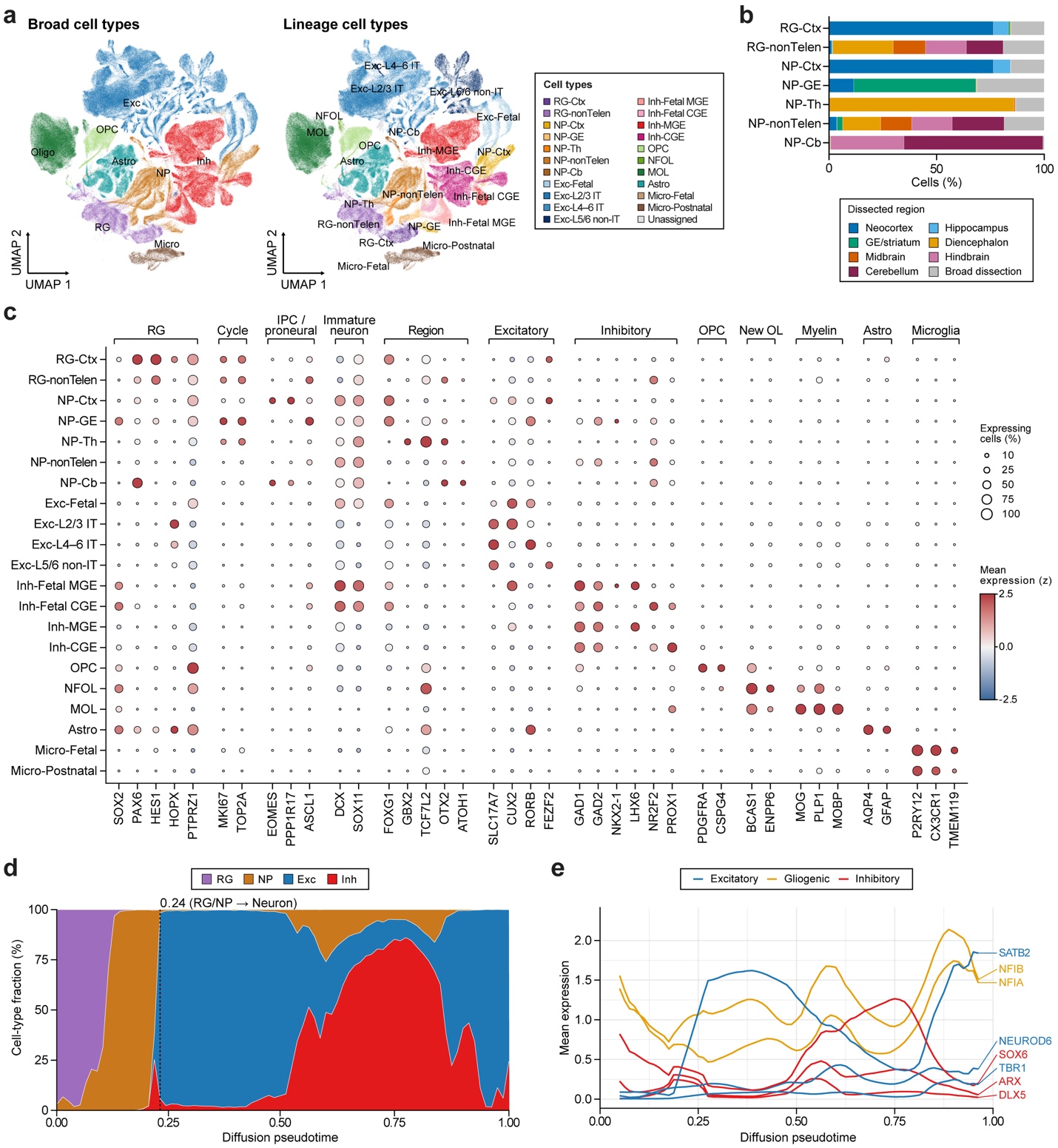

**Extended Data Fig. 1. Refined cell-type annotation and developmental ordering.** **a**, Comparison of broad and refined cell-type annotations. **b**, Regional composition of progenitor populations. **c**, Marker expression across annotated cell types (388,455 cells). Expression is standardized across cell types for each gene. **d**, Cell composition along rank-normalized diffusion pseudotime. The dashed line marks the first bin containing fewer than 50% radial glia and neuronal precursors combined. **e**, Smoothed marker expression along the same ordering. Diffusion pseudotime differs from the learned potential. Cell-type abbreviations follow **Fig. 1**.

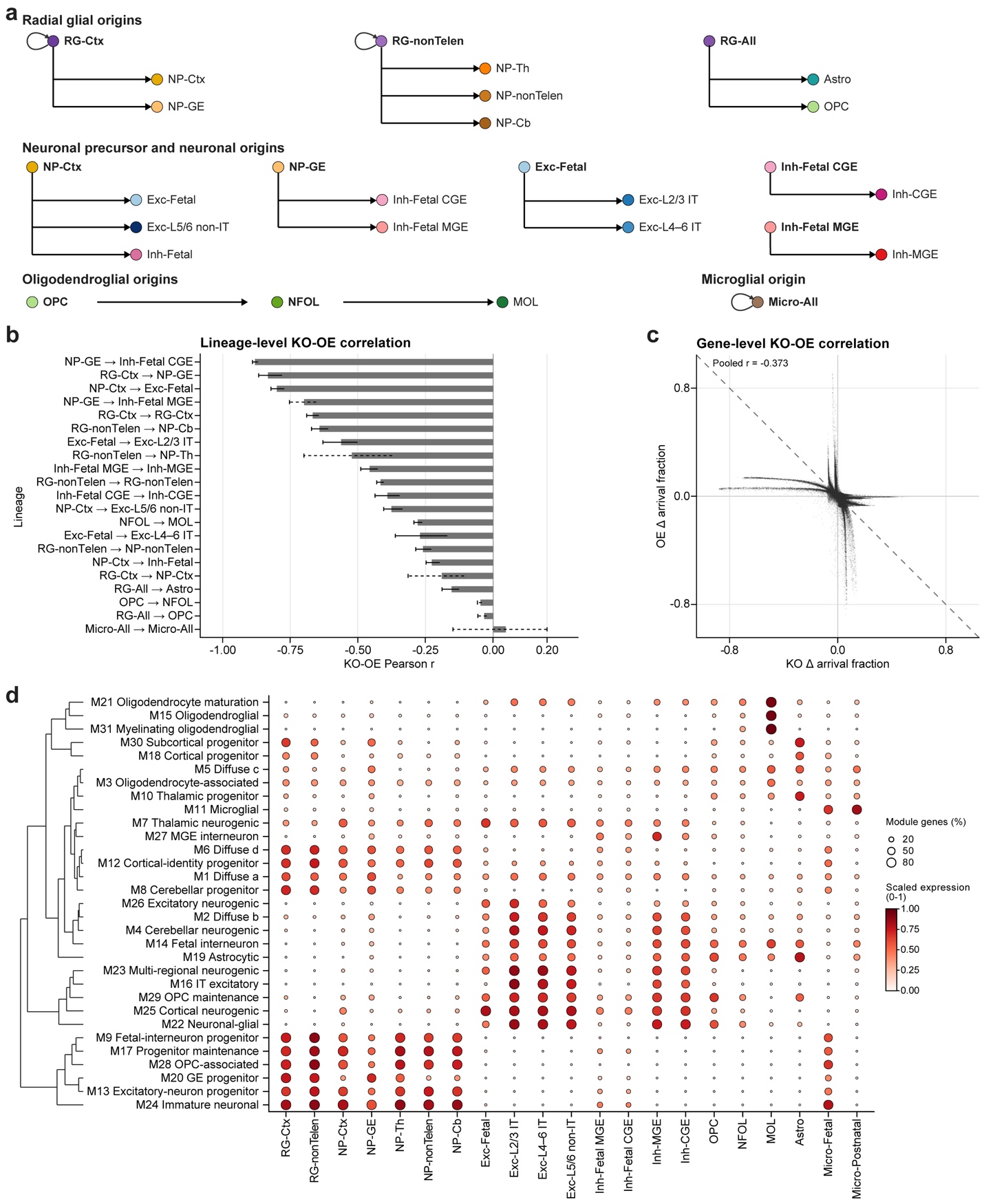

**Extended Data Fig. 2. Defined lineages and KO–OE relationships.** **a**, Starting and endpoint populations defining the simulated lineages, including identity retention. RG-All pools radial-glial types, Micro-All pools fetal and postnatal microglia, and Inh-Fetal pools fetal MGE/CGE types. **b**, KO–OE correlations across 14,044 genes for each lineage. Bars, 95% bootstrap CIs across simulation replicates. Dashed intervals flag unstable endpoints or WT floor/ceiling effects. **c**, Paired mean effects across the 17 non-boundary lineages, with each point representing a gene–lineage pair. Dashed line, equal and opposite effects. **d**, Module expression by cell type. Color, mean gene-scaled expression; dot area, proportion of module genes with scaled expression >0.5.

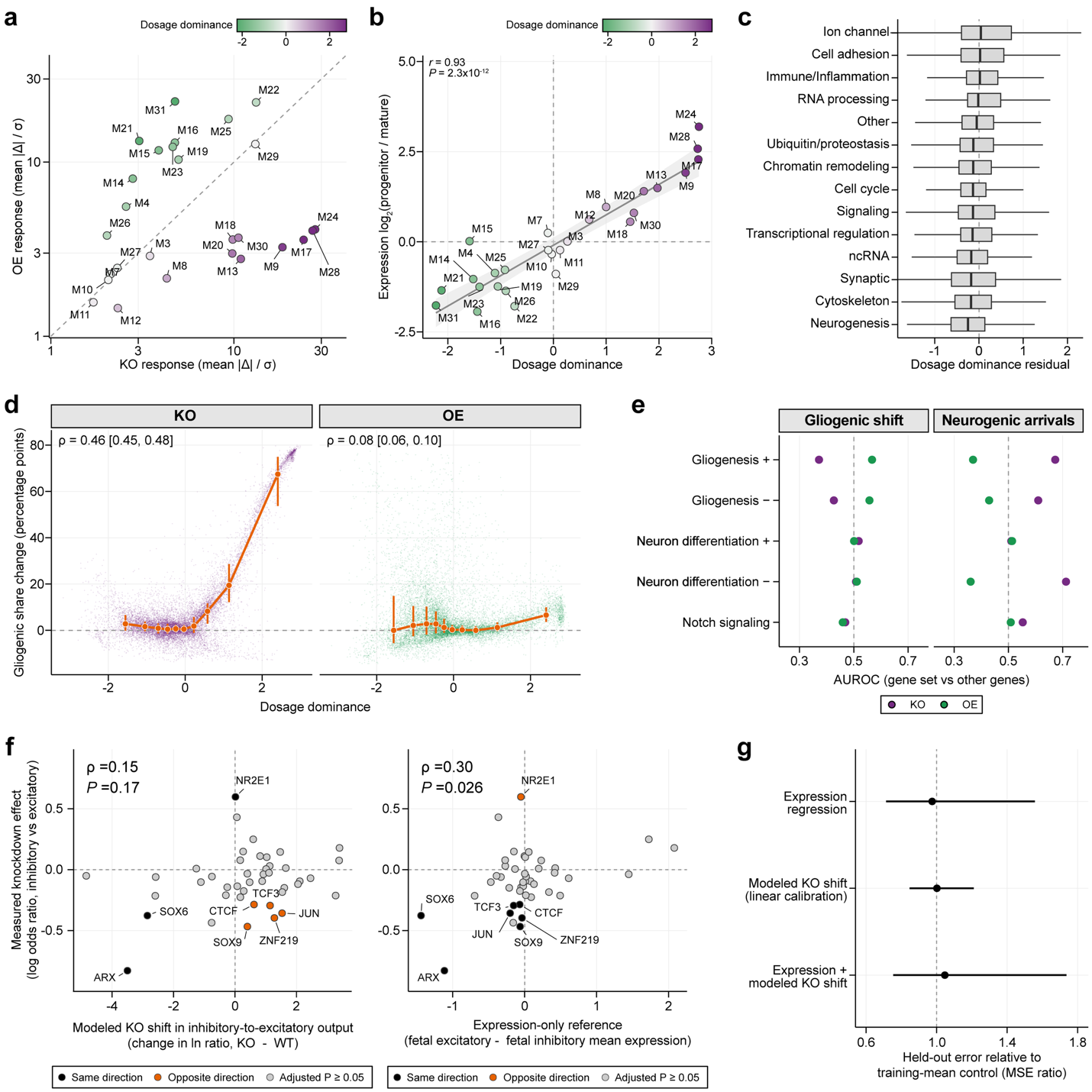

**Extended Data Fig. 3. Developmental expression, dosage responses and comparison with experimental fate effects.** **a**, Comparison of noise-scaled KO and OE response magnitudes across 27 modules. The dashed line indicates equal magnitudes. **b**, Relationship between the progenitor-to-mature expression ratio and dosage dominance across 27 modules. Line and shading indicate a linear fit and 95% confidence band. Two-sided Pearson test. **c**, Dosage dominance after adjustment for expression ratio and mean expression, grouped by functional class across 14,044 genes. Boxes show the median and interquartile range (IQR), with whiskers extending to the most extreme values within 1.5 × IQR. **d**, Relationship between dosage dominance and WT-relative changes in gliogenic share under KO and OE across 14,044 genes. Orange points and bars show decile medians and IQRs. Brackets give 95% bootstrap confidence intervals for Spearman correlations. **e**, GO-set comparisons after excluding M30/M10 genes (left) and for WT-relative changes in neuronal-precursor endpoint counts using all genes (right). AUROC > 0.5 indicates higher responses in the set than in other genes. **f**, Measured knockdown effects for 43 transcription factors compared with modeled KO shifts (left) and an expression-only reference (right). Directional matches and mismatches are highlighted for factors with experimental Benjamini–Hochberg-adjusted *P* < 0.05. Spearman correlations use one-sided permutation tests. **g**, Prediction error of regressions using expression, modeled KO shifts or both, evaluated by leave-one-factor-out cross-validation across 43 factors. Points and bars show mean squared error relative to a training-mean control and 95% confidence intervals from 2,000 factor-level bootstrap resamples.

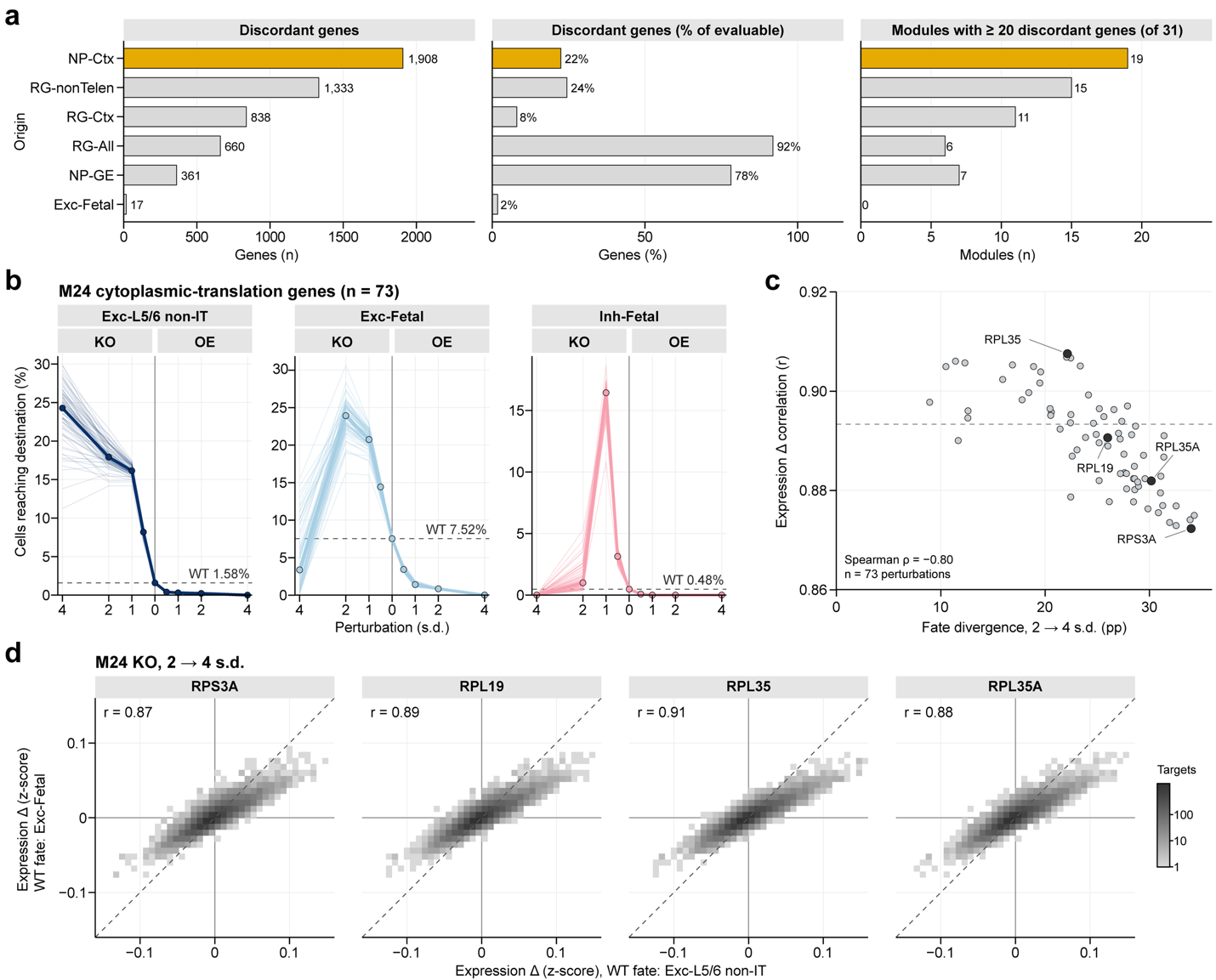

**Extended Data Fig. 4. Destination discordance and downstream responses of M24 genes.** **a**, Numbers and fractions of discordant genes, and numbers of modules containing at least 20 such genes, by origin. Discordance denotes different response classes in at least two destinations. **b**, Percentage of simulated cells reaching each endpoint after perturbation of M24 genes. Thin lines, individual gene means; thick lines, medians; dashed lines, WT values. **c**, Additional fate divergence from 2 to 4 s.d. KO versus expression-response correlation for each M24 perturbation. Fate divergence is the difference between additional shifts toward layer 5/6 non-IT and fetal excitatory states, in pp. Dashed line, correlation of the perturbation-averaged responses. **d**, Target-gene changes for perturbations with the lowest, median and highest correlations in **c**, and *RPL35A*. Expression changes are input-subtracted in the two WT-fate groups. Dashed lines, equal responses.

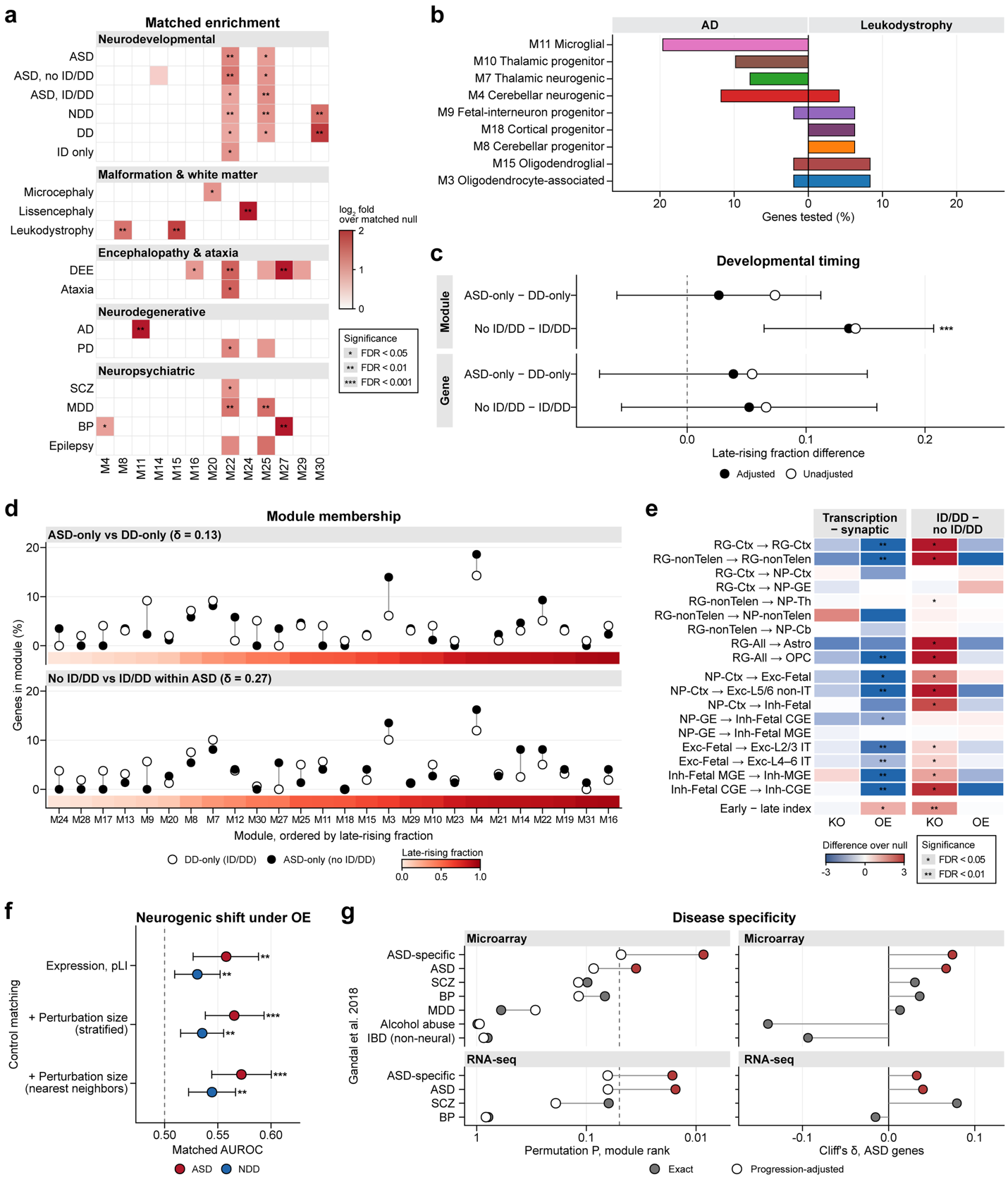

**Extended Data Fig. 5. Disease-program comparisons and developmental and cortical contexts of ASD risk.** **a**, Covariate-matched enrichment of disease–module associations. Gray, untested. **b**, Module distributions of nonoverlapping AD and leukodystrophy genes (n = 51 and 48). Modules containing ≥3 genes in either group are displayed. **c**, Late-rising fraction differences between exclusive disease or clinical groups, at module and gene levels. Bars, 95% CIs for expression- and constraint-adjusted differences. Group sizes for module/gene timing are 86/98 and 106/116 for ASD-only/DD-only, and 74/159 and 90/190 for no-ID/DD/ID/DD. Asterisks indicate nominal *P values from the covariate-adjusted models*. **d**, Module membership of groups in **c**, ordered by module timing. δ, unadjusted Cliff's delta. **e**, Functional-class and clinical-stratum contrasts in response magnitude beyond matched expectations. Final row, early-minus-late index. **f**, Neurogenic OE drive against matched backgrounds. Points, mean AUROC; intervals, 2.5th–97.5th percentiles across matched backgrounds. **g**, Concordance between modeled KO expression signatures and postmortem cortical changes. Left, ranking of the three ASD-associated modules before and after developmental-progression adjustment; dashed line, *P* = 0.05. Right, expression- and constraint-stratified Cliff's delta for all 290 ASD genes. ASD-specific contrasts remove shared psychiatric-disease changes. BP, bipolar disorder; IBD, inflammatory bowel disease.

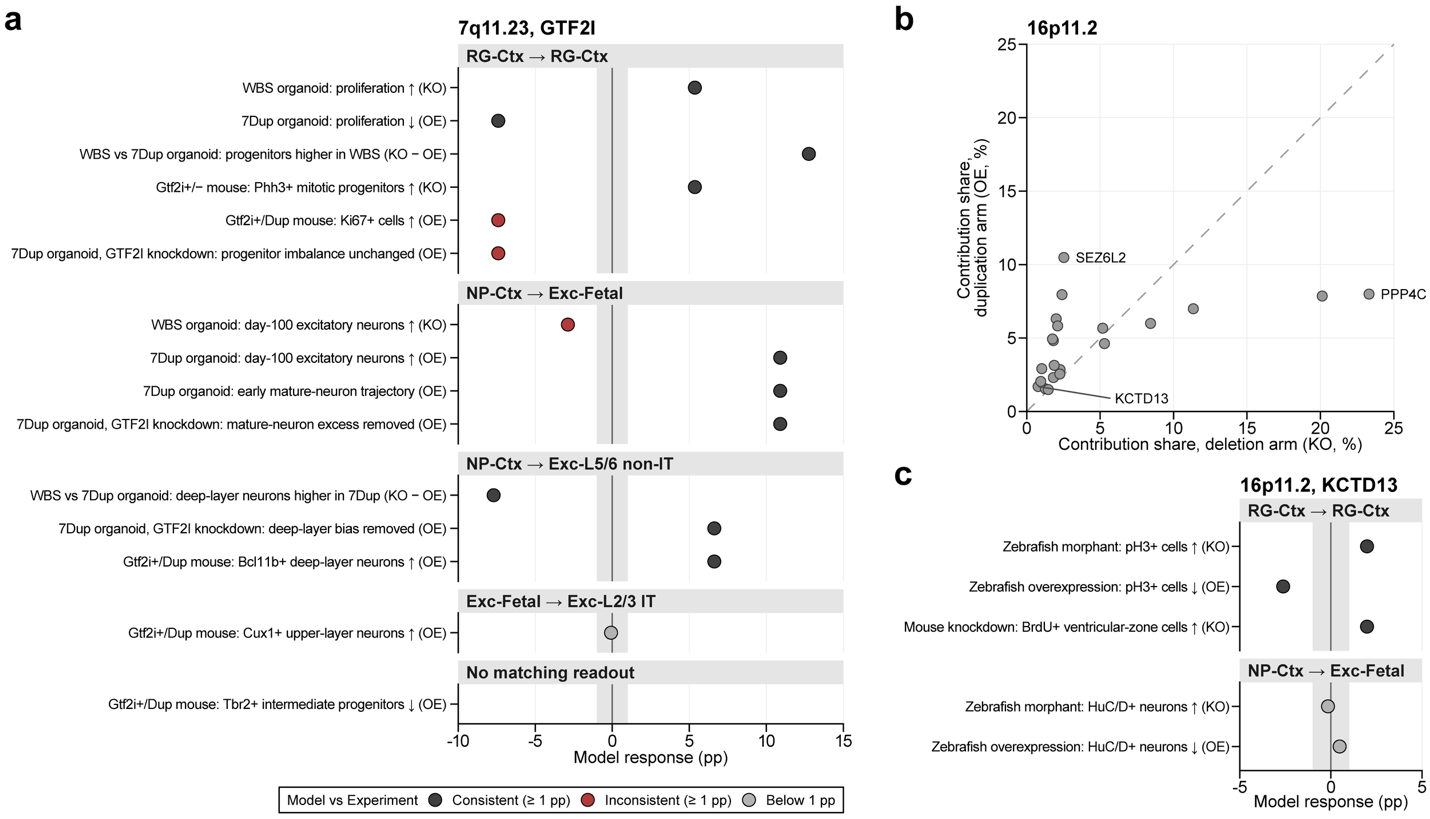

**Extended Data Fig. 6. CNV-related dosage experiments compared with modeled responses.** **a**, Fifteen reported 7q11.23 findings and corresponding model responses. The KO − OE contrast represents deletion-versus-duplication comparisons. Tbr2-positive intermediate progenitors lack a matching model readout. **b**, Within-arm response-score shares for 16p11.2 genes; dashed line, equal shares. **c**, Comparison of reported *KCTD13* phenotypes with corresponding model responses. In a and **c**, points are WT-relative changes in endpoint percentage (pp), and gray bands mark absolute changes below 1 pp. Radial-glial retention provides an indirect comparison with proliferation; multiple findings can share one model response.

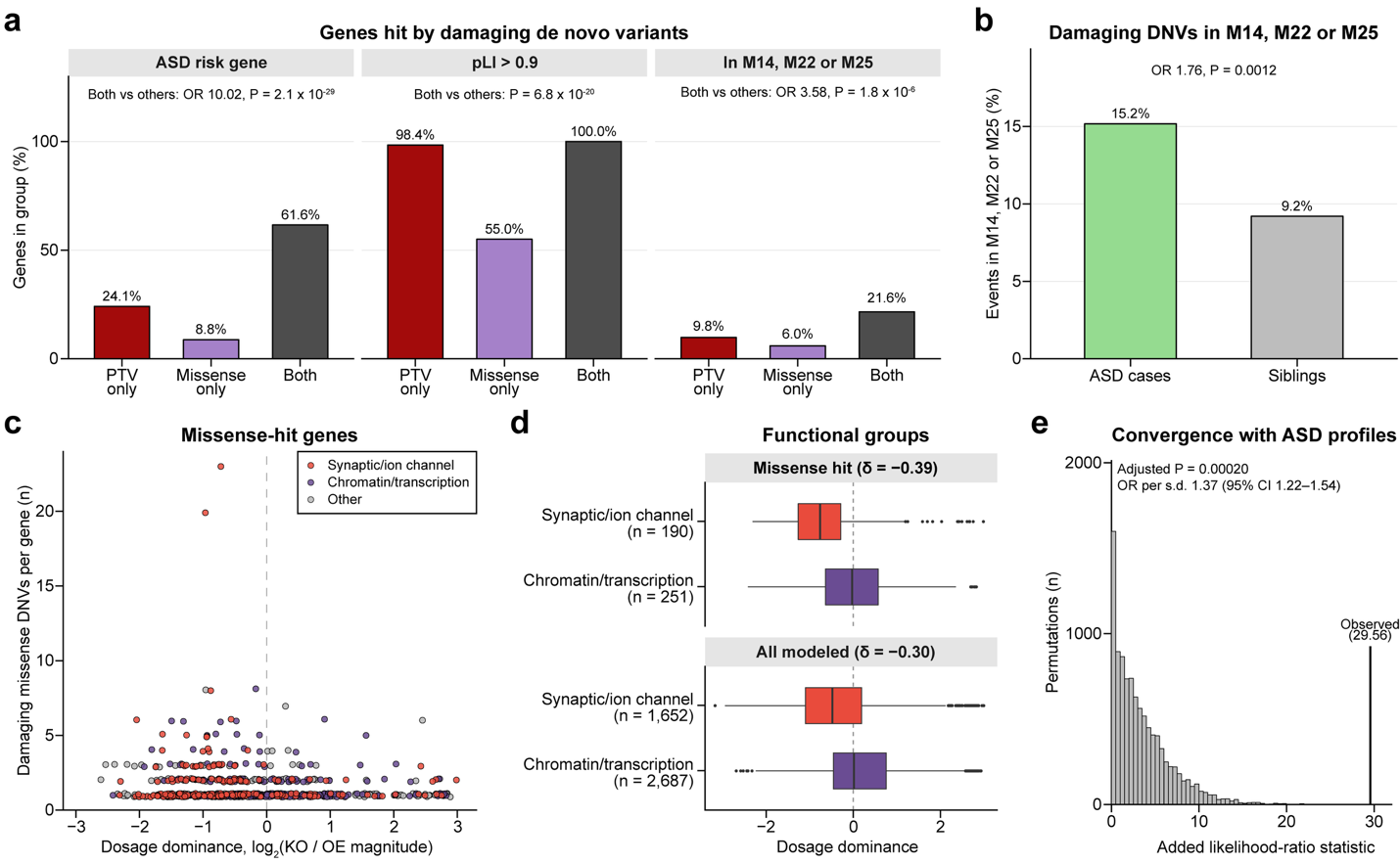

**Extended Data Fig. 7. De novo variant-associated genes and perturbation profiles.** **a**, Properties of genes hit by damaging PTVs only (n = 316), missense variants only (n = 705) or both (n = 125). ASD-risk fractions use the annotated subsets (311, 638 and 125 genes). **b**, Pooled de novo events in M14, M22 or M25 among cases (323/2,129) and siblings (38/413). The unit is the variant event. **c**, Relationship between dosage dominance and damaging missense variant counts per gene. **d**, Differences in dosage dominance between functional classes among missense-hit genes and among all modeled genes, including the missense-hit genes. Boxes, median and IQR; whiskers, 1.5 x IQR. δ, gene-level Cliff's delta. **e**, Convergence of 376 recurrently hit genes with 126 nonoverlapping ASD reference genes. Histogram, permutation distribution; line, observed statistic. The OR per s.d. is adjusted for constraint and co-expression.

**Supplementary Notes**

**Supplementary Note 1. Supporting evidence for cell-type annotation and lineage specificity**

This note details the regional, developmental and marker evidence used to assign the 21 cell-type labels. The overall mapping procedure and treatment of unassigned clusters are described in Methods. Cluster mappings and supporting measurements are provided in **Supplementary Table 1a**. Responses toward microglial states from neural progenitor origins are examined below.

*Radial glia and neuronal precursors*

Radial glia (RG) were annotated using the combined criteria described in Methods, with *HES1* expression providing supporting evidence^1^. The five clusters annotated as neuroblasts in the original atlas were relabeled as neuronal precursors (NP). These comprised neuron-committed cells, including dividing neuronal intermediate progenitors and postmitotic immature neurons. This designation was supported by expression of intermediate-progenitor or proneural markers (*EOMES*, *PPP1R17*, *NEUROG2* or *ASCL1*) or immature-neuron profiles (*SOX11* with *DCX* or *STMN2* in non-cycling cells) (**Extended Data Fig. 1c**)^2-4^.

Regional composition and *FOXG1* expression guided the assignment of the two RG clusters to cortical RG (RG-Ctx) from the neocortex and hippocampus, and to RG of the diencephalon, midbrain and hindbrain (RG-nonTelen). The five NP clusters were assigned to NP of the cortex (NP-Ctx), ganglionic eminence (NP-GE), thalamus (NP-Th), cerebellum (NP-Cb) and the diencephalon, midbrain and hindbrain (NP-nonTelen) using atlas cluster identities, regional composition, and region- and lineage-associated markers, including *TCF7L2* for NP-Th, *ATOH1* and *ZIC1* for NP-Cb and DLX-family markers for NP-GE. Regional marker interpretation was informed by human developmental studies of the early brain, thalamus and cerebellum^3,5,6^. Regional identity was not inferred from any single marker.

*Excitatory and inhibitory neurons*

Excitatory neurons were classified as fetal excitatory neurons (Exc-Fetal) or as one of three mature classes: layer 2/3 intratelencephalic (IT; Exc-L2/3 IT), layer 4-6 IT (Exc-L4-6 IT), and layer 5/6 non-intratelencephalic (Exc-L5/6 non-IT). This classification combined developmental, laminar, and projection-class annotations with marker expression, including *CUX1* and *CUX2* for L2/3 IT, RORB for L4-6 IT, and *FEZF2* and *BCL11B* for L5/6 non-IT neurons^7^.

Inhibitory neurons were grouped by developmental stage and by medial or caudal ganglionic eminence (MGE or CGE) origin into MGE- and CGE-derived interneurons (Inh-MGE, Inh-CGE) and fetal MGE- and CGE-type inhibitory neurons (Inh-Fetal MGE, Inh-Fetal CGE). Origin-associated annotations were considered alongside marker profiles, including *LHX6*, *SST* and *PVALB* for MGE-derived interneurons and *NR2F2*, *VIP* and *ADARB2* for CGE-derived interneurons. Fetal inhibitory neuron clusters consisted almost entirely of cells from fetal samples and expressed *DLX5*. Within these clusters, *LHX6* expression supported MGE-type identity, whereas NR2F2 expression supported CGE-type identity. MGE/CGE-associated marker programs were interpreted in the context of published interneuron development^4^. LAMP5 was not treated as an exclusive marker of either origin.

*Glial populations*

Astrocytes (Astro) and oligodendrocyte precursor cells (OPCs) were retained as single labels. Oligodendrocytes were assigned to newly formed (NFOL) or mature (MOL) labels, and microglia to fetal (Micro-Fetal) or postnatal (Micro-Postnatal) labels, based on the age and developmental-stage proportions of the contributing samples. Marker profiles were inspected as supporting evidence, without changing these existing assignments. Astrocyte and OPC profiles included AQP4/GFAP and PDGFRA/CSPG4, respectively^7,8^. *BCAS1* and *ENPP6* supported an early oligodendrocyte differentiation program^9,10^. *MOG*, *PLP1* and *MOBP* provided complementary myelin-associated readouts^11^. These state-associated genes were not assumed to define exclusive NFOL/MOL boundaries. *P2RY12*, *CX3CR1* and *TMEM119* supported microglial identity^12^. Fetal and postnatal microglial labels were based on sample age, not on these markers.

*Responses toward microglial states*

We examined microglial endpoints from RG-Ctx, RG-nonTelen, pooled RG-All, NP-Ctx and NP-GE origins, combining fetal and postnatal microglial states. Endpoint assignment and the WT noise threshold followed the procedures used for the 21 defined lineages (Methods). Across ten simulation replicates of 500 starting cells per origin, no WT cells reached microglial states. Among 14,044 genes, no KO responses exceeded the threshold, whereas 0.10–0.44% of OE responses did.

By comparison, the median responsive-gene fraction across the 21 defined lineages was 56.2% under KO and 52.6% under OE. Even the three defined lineages averaging fewer than one WT endpoint cell per replicate had higher response rates. Among 205 starting–endpoint pairs outside the defined lineages, 77 reached the minimum response rate among defined lineages under KO or OE. These pairs are not all biologically impossible transitions, so this comparison does not estimate a global false-positive rate.

**Supplementary Note 2. Robustness of perturbation map and module clustering**

We assessed two sources of variability in module assignment: clustering randomness on a fixed perturbation map and changes in the map caused by simulation sampling. The first measures the reproducibility of the clustering procedure conditional on the input profiles. The second measures sensitivity to starting-cell selection and stochastic trajectories under the same trained model. **Supplementary Fig.** **1** separates these comparisons. **Supplementary Table 1e** provides complementary recovery measurements for individual modules.

*Perturbation profiles and primary module construction*

The primary map comprised 14,044 genes and 42 axes: knockout (KO) and overexpression (OE) effects for 21 lineages from 11 origin populations. KO and OE were implemented as initial-state shifts along gene-associated directions in scaled-PCA space, corresponding to decreases and increases of two gene-expression standard deviations, respectively (**Methods**). For gene g, condition c, lineage l and simulation replicate r, the effect was the perturbed endpoint arrival fraction minus the corresponding wild-type (WT) fraction. Each axis in the primary map was the arithmetic mean of this difference over replicates 1–10. Perturbed and WT simulations shared starting cells and Brownian increments within a replicate (**Methods**). These replicates measured simulation variation, not variation between biological donors. All 42 axes were retained for module construction, including the four lineages near a WT floor or ceiling that were excluded from the main KO–OE correlation interpretation.

Each axis was centered and scaled across genes using the population standard deviation of that axis. A 30-nearest-neighbor graph was constructed by exact Euclidean search, with Scanpy connectivity weights. On this fixed graph, weighted, undirected Leiden clustering used the modularity objective, resolution 7, and two iterations for each seed from 0 to 1,599. Within each partition, cluster centroids were calculated from the unstandardized mean effect profiles and then standardized across clusters separately for each axis. A cluster received the axis with the largest absolute centroid z-score if that magnitude was at least 1; otherwise, the cluster entered the non-specific pool. The axis label retained the KO/OE condition and origin–target lineage, but not the sign of the z-score. Clusters with the same label were merged, and lineage-associated groups containing fewer than 30 genes received a separate dropped-group label.

Consensus was calculated after this axis-based merging. For each gene pair i and j, the co-assignment frequency was F(i,j) = (1/S) Σ_s I[L_s(i) = L_s(j)], where S is the number of seeds and L_s is the group label in seed s. The non-specific pool and dropped genes each formed one distinct group in this calculation. Average-linkage hierarchical clustering of 1 − F was cut at distance 0.5. Each resulting group received the most frequent label across member genes and all seeds. Groups with the same label were merged again, followed by the 30-gene minimum-size rule. Thus, consensus summarizes reproducibility of the named perturbation groups rather than preserving every raw Leiden community.

The adopted partition contained 27 lineage-associated groups with 8,490 genes and a non-specific pool of 5,554 genes. In the second tier, the non-specific profiles were standardized within that pool and separated by k-means (k = 4, random seed 42, 20 initializations). Numbering the resulting groups by size yielded the 31 primary modules, including M1, M2, M5 and M6 as the four diffuse modules (**Supplementary Table 1d**). The focused, intermediate and distributed categories describe concentration of lineage signatures; they are not grades of membership reproducibility.

*Clustering-seed stability and analysis settings*

Partition agreement was measured with the adjusted Rand index (ARI), using all genes and treating the non-specific pool as one group. Module overlap was measured by Jaccard similarity, |A ∩ B|/|A ∪ B|. Each lineage-associated module was matched to the module with the largest overlap in the other partition, excluding non-specific and dropped groups. Matches were calculated in both directions and pooled with equal weight per module; they were not constrained to a one-to-one correspondence. Consequently, module overlap and ARI quantify different aspects of agreement. Neither metric in this comparison evaluates the separation of the four diffuse modules.

Seed-count sensitivity was assessed using batches of 200, 400, 800 and 1,600 seeds. For each count S, three disjoint batch pairs used seed ranges 2pS to (2p + 1)S − 1 and (2p + 1)S to (2p + 2)S − 1, for p = 0, 1 and 2. The criterion required ARI ≥ 0.95 and at least 95% of the bidirectional module matches to have Jaccard similarity ≥ 0.75 in all three pairs. At resolution 7, 200 and 400 seeds did not meet this criterion; 800 and 1,600 seeds did. The primary analysis retained 1,600 seeds, which should not be interpreted as the minimum count required to pass. At 1,600 seeds, mean ARI was 0.982 (range 0.974–0.991), mean best-match Jaccard similarity was 0.989, and all module matches in each pair exceeded 0.75.

Resolution sensitivity used integers 1–10, with two batches of 1,600 seeds at each resolution (0–1,599 and 1,600–3,199). The numbers of lineage-associated modules in the first batch were 13, 17, 21, 22, 23, 26, 27, 27, 27 and 27, respectively. The second batch gave the same counts except at resolution 5, where the second batch yielded 24 modules. Resolution 7 was therefore the lowest tested resolution at which the count remained at 27 through resolution 10. At resolutions 7–10, within-resolution seed-batch ARIs were 0.991, 0.974, 0.971 and 0.965, respectively. This scan used one batch pair per resolution, whereas the seed-count assessment used three. A plateau in module count does not establish identical memberships across resolutions or biological optimality of resolution 7.

Supplementary Fig. 1a also shows the legacy single-run procedure, for which neighbor-graph and Leiden seeds varied together. The 45 pairwise comparisons gave mean ARI 0.669 (range 0.594–0.770), compared with 0.982 for the three primary consensus batch pairs. Because graph-seed handling also differed, this contrast cannot isolate the effect of consensus alone. The pairwise comparisons reused partitions; the displayed ranges summarize observed variation and are not confidence intervals.

*Sensitivity to simulation variation*

A separate comparison used the stored pool of 100 simulation profiles. Ten permutations of replicate indices 1–100 were generated with random seed 20261019. For each permutation, the first and last 50 indices defined disjoint groups A and B. Means of the first k profiles within each group were compared for k = 1, 10, 20, 30, 40 and 50. Groups were nested across k within a permutation, and replicates were reused across permutations. Thus, the ten comparisons at each k are descriptive sensitivity checks rather than ten independent experiments.

For each mean profile, the axes were restandardized, and the exact 30-nearest-neighbor graph for that profile was reconstructed. The single-seed and consensus procedures used the same graph, Leiden objective, resolution 7, and group-label rules. The single partition used seed 10,000; consensus combined seeds 10,000–10,799 and therefore included that partition. This matched 800-seed protocol addresses simulation variation without the graph-procedure difference present in the legacy comparison. This analysis does not constitute a repeat of the primary 1,600-seed assignment.

Mean ARIs for single seed versus consensus clustering were 0.258 versus 0.303 at k = 1, 0.540 versus 0.624 at k = 10, 0.616 versus 0.694 at k = 20, 0.620 versus 0.701 at k = 30, 0.641 versus 0.720 at k = 40, and 0.676 versus 0.757 at k = 50 (**Supplementary Fig. 1b**). For disjoint 10-replicate means, consensus ARI ranged from 0.522 to 0.710 and mean bidirectional best-match Jaccard similarity was 0.528. For disjoint 50-replicate means, consensus ARI ranged from 0.727 to 0.801. These comparisons show greater agreement with additional simulation averaging under the tested procedure, while retaining appreciable membership variation. No independent-sample confidence interval or *P* value was assigned, and these sensitivity profiles did not replace the primary 10-replicate map.

*Recovery of individual modules*

We assessed recovery of gene membership for each adopted lineage-associated module after resampling perturbation profiles (**Supplementary Table 1e**). The recovery analysis used 20 split-half profiles and 20 pseudo-10-replicate profiles derived from the original ten replicates, each clustered with 800 seeds (10,000–10,799). Ten distinct complementary five-versus-five splits were selected with random seed 2026. For pseudo-10 profiles, residuals were defined as E_r = D_r − D̄, where D_r is a replicate profile, and D̄ is the mean of the ten profiles. Each pseudo-profile was D̄ + (1/10) Σ_r ε_r E_r, with independently sampled signs ε_r ∈ {−1, +1} and random seed 0. The 20 profiles share their original observations and mean; they are not independently simulated ten-replicate means. Split-half profiles also share observations across splits and use fewer replicates than the primary map. Neither family provides a formal bound on independent-replicate recovery.

For each of the 27 lineage-associated reference modules, recovery was quantified as the Jaccard similarity between the reference module's gene set and the best-matching group in each comparison partition. We recorded the median similarity across the 20 partitions, the fraction with Jaccard similarity ≥ 0.6, and the fraction retaining the reference construction-axis label. This last measurement concerns the KO/OE and origin–target lineage label used during clustering. Construction-axis retention does not test preservation of the sign of the effect, the full signature, the final biological name, or the separately assigned OE-driven/KO-driven annotation. The four diffuse modules have no individual recovery estimates in this analysis and are marked as not evaluated in **Supplementary Table 1e**.

Recovery varied substantially between modules. Median pseudo-10 Jaccard similarities included 0.268 for M12 and 0.339 for M11, compared with 0.887 for M15 and 0.888 for M22. Their corresponding split-half medians were 0.230, 0.360, 0.892 and 0.894. Construction-axis retention did not imply membership stability: M11 retained the construction-axis label in all pseudo-10 partitions despite low median Jaccard similarity. Conversely, M31 had a median pseudo-10 Jaccard similarity of 0.883 but retained the construction-axis label in only 55% of partitions. These measurements support interpreting module membership and module annotation separately, rather than assigning one confidence level to every named module.

*Scope of the robustness assessment*

The high fixed-map ARI establishes reproducibility of the first-tier partition under clustering-seed changes. The lower agreement across simulation profiles and heterogeneous module recovery limit claims about exact gene membership. The 31-module catalog is a defined reference for downstream analyses, with biological annotations conditional on that partition. These checks keep the atlas, trained model, perturbation definition and simulation endpoint fixed; they do not measure uncertainty from model retraining, donor sampling or alternative biological annotations, and do not provide independent experimental validation of perturbation effects. **Supplementary Note 3. Biological annotation of perturbation-response modules**

*Lineage, expression and functional evidence*

Biological identities were assigned by integrating lineage-specific perturbation effects, the cell-type expression patterns of module genes, and Gene Ontology (GO) biological-process and MSigDB Hallmark enrichment. Lineage effects were summarized by the difference between standardized overexpression and knockout module profiles. Cell-type expression summaries averaged gene-wise min–max-scaled cell-type means across module genes.

Modules whose dominant perturbation effect was an OE-driven increase in arrival at a mature fate were named according to that fate (for example, Microglial, MGE interneuron or Oligodendroglial). Modules with progenitor-associated perturbation programs were assigned progenitor identities (for example, Cortical progenitor or Thalamic progenitor). When lineage effects were non-specific, or expression and function supported another identity, naming followed expression or function (for example, Oligodendrocyte-associated). Disease gene sets were not used for naming. Modules lacking a lineage-specific perturbation signature were designated as distributed modules. Module names and supporting annotations are provided in **Supplementary Table 1d,** and the representative transition and OE-driven/KO-driven labels are defined in **Methods**.

*Functional enrichment*

Functional enrichment used the 14,044 perturbed genes as the input universe. GO biological-process overrepresentation was calculated for each module with clusterProfiler::enrichGO and org.Hs.eg.db, using SYMBOL identifiers and gene-set sizes of 15–500. GO enrichment used Benjamini–Hochberg adjustment, a *P* value cutoff of 0.05 and a q-value cutoff of 0.2. Redundant terms were reduced using Wang semantic similarity with a cutoff of 0.7, retaining the term with the smallest adjusted *P* value.

Hallmark overrepresentation used the 50 human MSigDB Hallmark sets obtained through msigdbr and clusterProfiler::enricher, with gene-set sizes of 10–500. Hallmark terms with adjusted P < 0.05 were used as annotation evidence. Adjustment was performed within each module and gene-set collection, not jointly across all pairs of modules and terms. These enrichment analyses informed module annotation and were distinct from the mutually exclusive functional-class composition displayed in **Fig. 1e**.

**Supplementary Note 4. Simulation variability and sensitivity analyses**

Simulation variability was assessed by resampling the ten simulation runs 2,000 times. Each resample contained ten runs drawn with replacement, using matched run indices for KO, OE and WT. Endpoint counts were repooled, and gliogenic share changes recalculated relative to the resampled WT counts. Spearman correlations with dosage dominance were recomputed across genes with finite shares and across the 27 lineage-associated modules using their mean share changes. Dosage dominance and module assignments were held fixed. The 2.5th and 97.5th percentiles summarized variability across resamples. Resampling used NumPy’s default random-number generator with seed 20261110.

Share-change reliability was calculated for genes with finite shares across all ten runs. The one-way random-effects single-run intraclass correlation was ICC (1,1) = (MSbetween − MSwithin)/[MSbetween + 9MSwithin], where MSbetween and MSwithin were the between-gene and within-gene mean squares, respectively^13^. Genes were treated as subjects and runs as repeated measurements. Reliability of the ten-run mean was estimated as 10ICC/(1 + 9ICC). Across 14,044 genes, single-run ICC values were 0.911 for KO and 0.752 for OE; the estimated reliability of the ten-run mean was 0.990 for KO and 0.968 for OE. Run-resampling intervals for the gene-level share-change correlation were 0.371–0.567 for KO and −0.113–0.273 for OE. Corresponding module-level intervals were 0.579–0.686 and 0.129–0.347. These intervals describe simulation variability with dominance held fixed; gene- and module-resampling intervals are described in Methods (**Supplementary Table 2d**).

To assess sensitivity to the lineage readouts included in dosage dominance, we excluded radial glia-to-astrocyte and radial glia-to-OPC transitions and recalculated dominance using the remaining 19 lineages, while retaining the original noise scales and regularization constants. Fate measures were unchanged. Associations with share change and absolute gliogenic shift were recalculated using the same Spearman and bootstrap procedures as in the primary analysis. For absolute gliogenic shift, gene-level correlations with 19-lineage dominance were ρ = 0.542 for KO and −0.209 for OE, compared with 0.539 and −0.260 using all 21 lineages.

**Supplementary Note 5. Supporting analyses of rare and common risk across perturbation modules**

*De novo gene and variant analyses.*

Damaging de novo variants in ASD probands affected 316 genes through PTVs only, 705 through missense variants only and 125 through both. Of the 125 shared-hit genes, 77 were ASD risk genes and all had pLI > 0.9 (**Supplementary Table 6a**). Compared with the other two gene classes combined, shared-hit genes were enriched in M14, M22 and M25 (27 of 125 genes, 21.6%, versus 7.2% of the other two classes; OR = 3.58, P = 1.8 x 10^-6^) (**Extended Data Fig. 7a**)**.** These modules contained 15.17% of damaging de novo events in cases and 9.20% in siblings **(**OR = 1.76, *P* = 0.0012) (**Extended Data Fig. 7b**). Among model-represented missense-hit genes, synaptic and ion-channel genes (n = 190) had lower median dosage dominance than chromatin-remodeling and transcriptional-regulatory genes (n = 251, −0.761 versus −0.027, *P* = 2.62 x 10^-12^, Cliff's δ = −0.389). The corresponding contrast was also present across all model genes (n = 1,652 versus 2,687, medians −0.481 versus 0.015, *P* = 1.43 x 10^-63^), with no evidence that the difference was specific to missense-hit genes (interaction *P* = 0.103) (**Extended Data Fig. 7d**).

*Clinical phenotypes of de novo variant carriers.*

Among ASD probands, carriers of any damaging de novo variant had lower full-scale and nonverbal IQ, Vineland adaptive scores and DCDQ motor-coordination scores than non-carriers. They also had higher SRS scores. In comparisons with other damaging-DNV carriers, carriers in ASD risk genes and in shared-hit genes had lower IQ, Vineland and DCDQ scores. ASD risk-gene carriers also had higher SRS scores, and carriers in missense-only genes had lower SCQ, SRS and ADI-R communication scores and higher full-scale IQ and DCDQ scores. Among fate-module and fate-axis groups, M22 and neurogenic-gene carriers had lower Vineland scores, and gliogenic-gene carriers had higher Vineland composite, communication, daily living and socialization scores than other carriers (**Supplementary Table 6f**).

*Matched gene sets for module PRS*

Candidate genes were excluded if their ±100-kb windows overlapped any window of the target module. Each matched set contained distinct genes and preserved the target gene count and chromosome-specific gene counts. For each matching covariate listed in Methods, the absolute standardized mean difference between target and matched genes was required to be at most 0.1, using the square root of the mean of the two sample variances as the denominator. The number of unique SNPs and total genomic span covered by the union of gene windows were each required to lie within 10% of the target value. The summed LD score over the SNP union was also required to differ from the target by no more than 10%.

Four chains, each initialized by mixed-integer linear programming, used tempered heat-bath sampling to explore eligible gene sets. The retained chain state satisfied all matching constraints. Auxiliary states allowed temporary violations to improve movement between eligible sets. Each sampling round updated the constrained state and one randomly selected auxiliary state and attempted an exchange between adjacent states. After a module-specific burn-in of 50,000–706,000 rounds, 1,000 sets per chain were retained at intervals of 50–706 rounds, yielding 4,000 matched sets per module. Burn-in was 1,000 times the retention interval for each module. Sets could recur across retained samples. The empirical comparison used the meta-analyzed PRS transmission statistic from each retained set, as defined in **Methods**.

*Sensitivity of common-variant associations.*

Of the two modules whose PRS transmission exceeded the matched set expectation after correction, M20 remained below the correction threshold when its empirical *P* was increased by two Monte Carlo standard errors (adjusted *P* = 0.020 to 0.039), whereas M7 exceeded it after a one-standard-error increase (adjusted *P* = 0.047 to 0.054; 4,000 matched sets) (**Supplementary Table 6b**). Including MHC genes changed the corrected significance status of only SCZ–M14 among the 243 disorder–module tests, with no change for ASD. Joint adjustment for s_het and coding-sequence length removed corrected support for M22–ANX, M25–BIP/ANX and M7–BIP/MDD (**Supplementary Table 6c)**. No module passed correction in the correlated six-disorder summary (**Supplementary Table 6g**).

For AD, M11 Microglial did not pass correction across 27 modules in the European-ancestry analysis (adjusted *P* = 0.051) but did in the multi-ancestry sensitivity analysis (adjusted *P* = 0.045). Neither association passed correction after joint covariate adjustment (adjusted *P* = 0.13 and 0.11). The analyses share samples and both use European LD, so they do not provide independent replication (**Supplementary Table 6c**).

**Supplementary Figures**

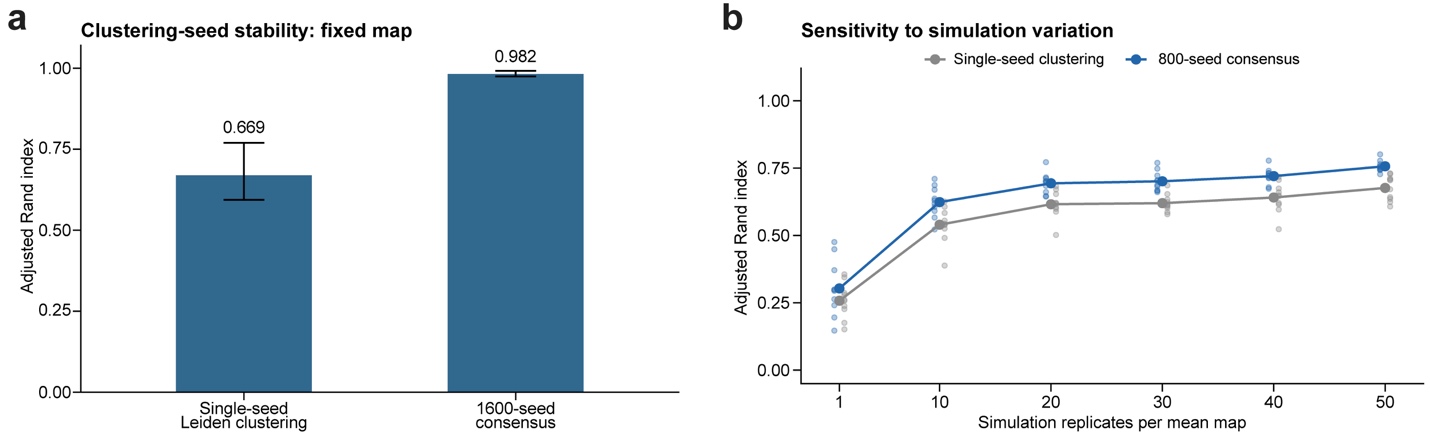

**Supplementary Fig. 1. Module-clustering stability and sensitivity to simulation variation.** **a**, Partition agreement on the fixed mean map. Bars and whiskers, mean ARI and observed range across 45 single-run pairs or three consensus-batch pairs. **b**, Agreement between maps averaged over disjoint sets of simulation replicates. Points, ten comparisons per setting; lines, means. Replicates recur across comparisons. The primary assignment uses the 1,600-seed consensus described in **Supplementary Note 2**.

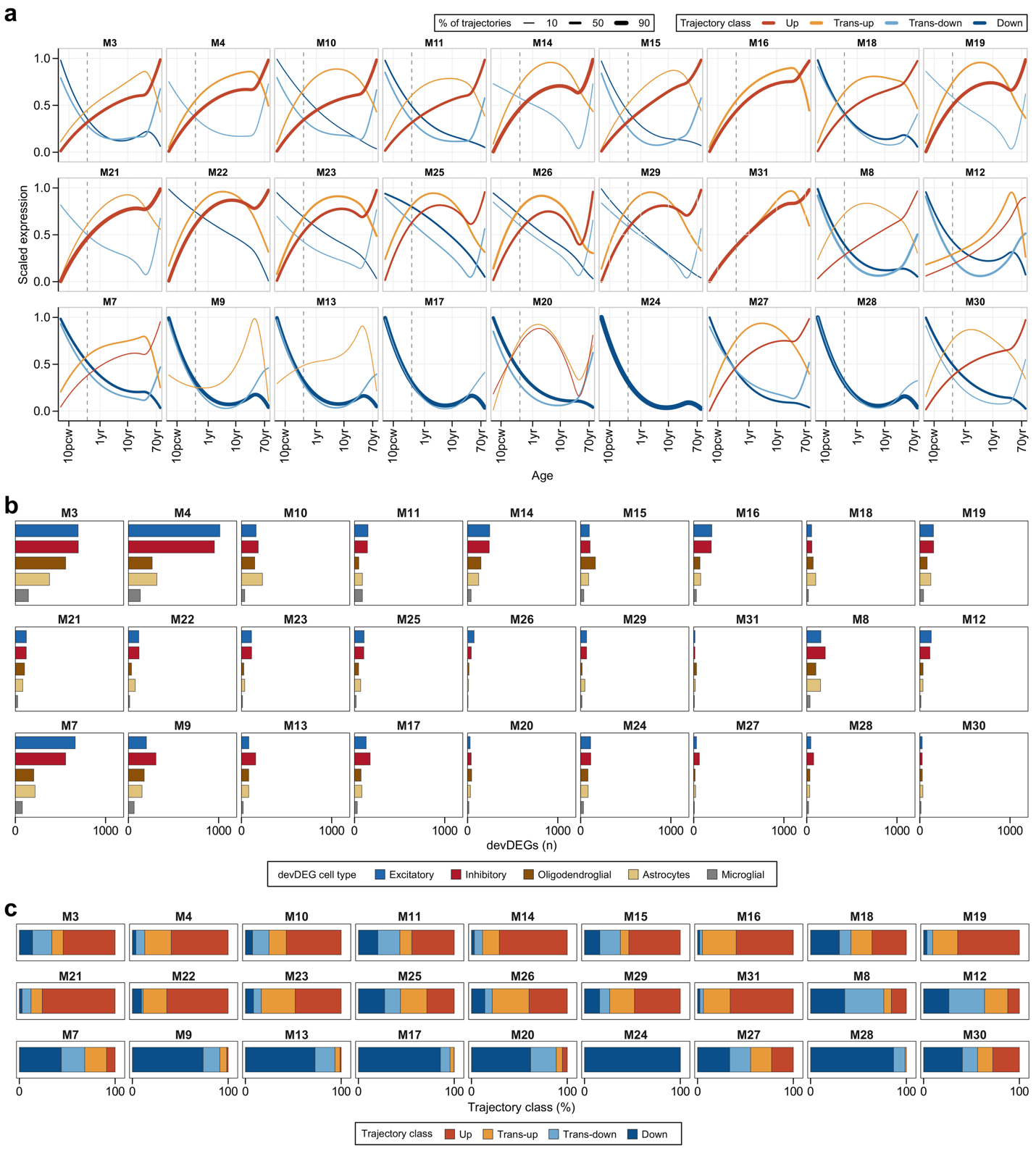

**Supplementary Fig. 2. Developmental expression trajectories and their composition across 27 lineage-associated modules.** **a**, Mean scaled expression trajectories within each trajectory class. Line width and opacity indicate the proportion of trajectories in each class. Classes representing <5% of a module’s trajectories are omitted. Dashed lines mark birth. **b**, Numbers of developmentally dynamic genes (devDEGs) in each cell group. A gene can contribute to more than one cell group. **c**, Proportions of gene–cell-group trajectories in each class, including classes omitted from a. Modules follow the same order in all three panels.

**Supplementary Tables**

**Supplementary Table 1.** Genome-wide in silico perturbation profiles and fate-regulatory module annotations.

**Supplementary Table 2**. Developmental timing, expression position and neurogenic-gliogenic allocation in relation to dosage dominance

**Supplementary Table 3.** Strength-dependent fate responses, destination discordance and the *NFIA* and M24 examples

**Supplementary Table 4.** Disease gene sets, developmental response programs and ASD analyses.

**Supplementary Table 5.** CNV intervals as dosage proxies for the knockout and overexpression perturbation map

**Supplementary Table 6.** Rare and common genetic risk across perturbation modules

**Supplementary References**

1	Pollen, A. A. *et al.* Molecular identity of human outer radial glia during cortical development. *Cell* **163**, 55-67 (2015). <https://doi.org/10.1016/j.cell.2015.09.004>

2	Pebworth, M. P., Ross, J., Andrews, M., Bhaduri, A. & Kriegstein, A. R. Human intermediate progenitor diversity during cortical development. *Proc Natl Acad Sci U S A* **118** (2021). <https://doi.org/10.1073/pnas.2019415118>

3	Eze, U. C., Bhaduri, A., Haeussler, M., Nowakowski, T. J. & Kriegstein, A. R. Single-cell atlas of early human brain development highlights heterogeneity of human neuroepithelial cells and early radial glia. *Nat Neurosci* **24**, 584-594 (2021). <https://doi.org/10.1038/s41593-020-00794-1>

4	Shi, Y. *et al.* Mouse and human share conserved transcriptional programs for interneuron development. *Science* **374**, eabj6641 (2021). <https://doi.org/10.1126/science.abj6641>

5	Kim, C. N., Shin, D., Wang, A. & Nowakowski, T. J. Spatiotemporal molecular dynamics of the developing human thalamus. *Science* **382**, eadf9941 (2023). <https://doi.org/10.1126/science.adf9941>

6	Aldinger, K. A. *et al.* Spatial and cell type transcriptional landscape of human cerebellar development. *Nat Neurosci* **24**, 1163-1175 (2021). <https://doi.org/10.1038/s41593-021-00872-y>

7	Hodge, R. D. *et al.* Conserved cell types with divergent features in human versus mouse cortex. *Nature* **573**, 61-68 (2019). <https://doi.org/10.1038/s41586-019-1506-7>

8	Marques, S. *et al.* Oligodendrocyte heterogeneity in the mouse juvenile and adult central nervous system. *Science* **352**, 1326-1329 (2016). <https://doi.org/10.1126/science.aaf6463>

9	Fard, M. K. *et al.* BCAS1 expression defines a population of early myelinating oligodendrocytes in multiple sclerosis lesions. *Sci Transl Med* **9** (2017). <https://doi.org/10.1126/scitranslmed.aam7816>

10	Xiao, L. *et al.* Rapid production of new oligodendrocytes is required in the earliest stages of motor-skill learning. *Nat Neurosci* **19**, 1210-1217 (2016). <https://doi.org/10.1038/nn.4351>

11	Yellajoshyula, D. *et al.* The DYT6 Dystonia Protein THAP1 Regulates Myelination within the Oligodendrocyte Lineage. *Dev Cell* **42**, 52-67 e54 (2017). <https://doi.org/10.1016/j.devcel.2017.06.009>

12	Bennett, M. L. *et al.* New tools for studying microglia in the mouse and human CNS. *Proc Natl Acad Sci U S A* **113**, E1738-1746 (2016). <https://doi.org/10.1073/pnas.1525528113>

13	Shrout, P. E. & Fleiss, J. L. Intraclass correlations: uses in assessing rater reliability. *Psychol Bull* **86**, 420-428 (1979). <https://doi.org/10.1037//0033-2909.86.2.420>
